# Mapping the sequence preference of the generalist class II lanthipeptide synthetase ProcM by mRNA display

**DOI:** 10.64898/2026.08.19.745792

**Authors:** Yao Ouyang, Hassan Nadeem, Yuki Goto, Diwakar Shukla, Wilfred A. van der Donk

## Abstract

The biosynthetic machineries of ribosomally synthesized and post-translationally modified peptides (RiPPs) are often substrate tolerant. A remarkable example is the class II lanthipeptide synthetase ProcM, which naturally functions as a generalist enzyme that has not evolved to use a specific substrate during its evolutionary history. Although ProcM has been studied extensively, the sequence features associated with productive modification remain underexplored. In this study, we use the ultrahigh-throughput mRNA display technique to map the sequence compatibility of ProcM across a focused library. This approach expands the landscape of ProcM reactivity beyond native substrates and individually characterized variants. Machine learning (ML) analysis revealed that the dataset contains sequence patterns associated with ProcM reactivity, with models achieving a balanced accuracy of 0.73. This moderate predictive performance contrasts with the near-perfect accuracy reported for specialized enzymes. This suggests that predicting substrate compatibility for generalist enzymes is more challenging due to uneven representation of reactive and nonreactive sequences and limitations of the available data. Our results provide a high-throughput view of ProcM reactivity and highlight differences with previous high-throughput studies on substrate selectivity of RiPP modification enzymes. Future studies will need to assess whether these differences are common when comparing generalist with specialist enzymes.

**Table of Contents Graphic:** 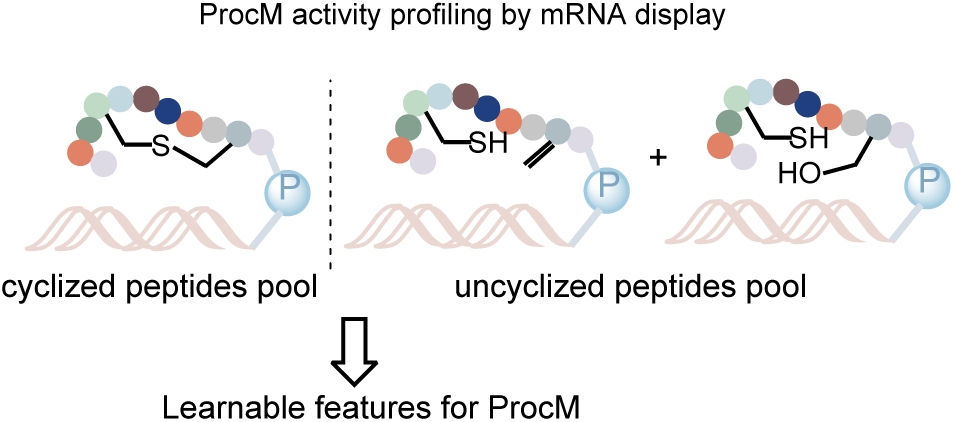

## Introduction

Lanthipeptides^1^ constitute a major class of ribosomally synthesized and post-translationally modified peptides (RiPPs).^2,3^ RiPP biosynthesis is initiated with a linear precursor peptide, containing a leader peptide that is responsible for enzyme recruitment, and a core peptide, where the post-translational modifications (PTMs) occur. The class-defining feature of lanthipeptides is the formation of lanthionine (Lan) and methyllanthionine (MeLan) crosslinks catalyzed by various classes of synthetases.^3^ These enzymes often exhibit tolerance to non-natural sequences of the core peptides.^4–11^ Thus, substantial efforts have focused on bioengineering of lanthipeptides.^12–27^

The class II lanthipeptide synthetase ProcM was originally discovered in the genome of the marine picocyanobacteria *Prochlorococcus* MIT9313 along with 29 precursor peptides (Figure S1), representing an unexpected example of extreme natural substrate tolerance.^6^ ProcM is a bifunctional lanthipeptide synthetase that sequentially dehydrates Ser or Thr to form dehydroalanine (Dha) or dehydrobutyrine (Dhb), respectively, and catalyzes the intramolecular Michael-type addition of Cys to generate Lan and MeLan (**Figure 1A**). Although the final mature products are highly divergent in sequence, ring size and ring pattern, the leader peptides that recruit ProcM are highly conserved (Figure S1).^6^ Previous research suggested that the linear precursor sequences rather than the enzyme dictate the regioselectivity of (methyl)lanthionine formation,^28^ and that the first cyclization is under kinetic control and determines the regioselectivity of the subsequently formed rings.^29^ The discovery of this highly substrate-tolerant enzyme has inspired its utilization in bioengineering applications.^18,26,28,30,31^ By randomizing the residues within the rings, ProcM was demonstrated to process unnatural sequences with leader peptides of ProcA2.8 and ProcA3.3 attached to a combinatorial library of nonnative core peptides.^28,30^ However, those studies focused on individually characterized mutants in conserved scaffolds. As a result, it is still unclear what sequence features ProcM prefers and whether this information can be extracted in a high-throughput manner.

**Figure 1.**
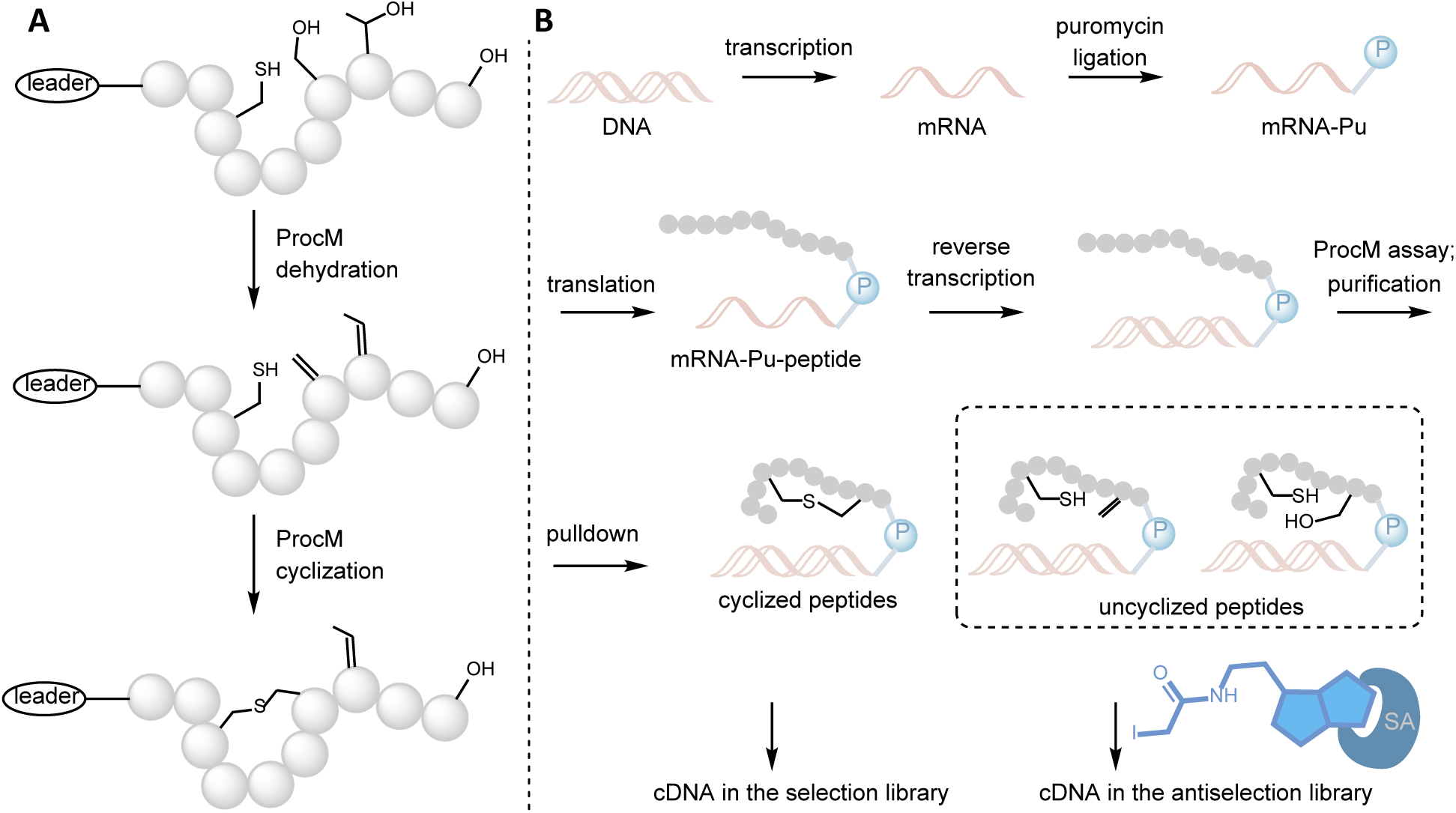
(A) Bifunctional class II lanthipeptide synthetase ProcM catalyzes a two-step reaction. (B) Design of the mRNA display workflow to map the preferences of ProcM, where the cyclized peptides will be separated from the uncyclized peptides by reaction of Cys residues with a biotinylated electrophile. SA, streptavidin.

Large language models have emerged as powerful tools for extracting latent sequence features and preferences of RiPP biosynthetic enzymes.^32–35^ Models based on deep learning frameworks such as masked language modeling can capture higher-order dependencies within peptide sequences and infer functional patterns. The performance of the models heavily relies on data accessibility and quality, requiring a clean and minimally biased training dataset to ensure robust generalization. The intrinsic substrate tolerance of several RiPP biosynthetic machineries^3,36^ has enabled high-throughput mapping of enzyme specificity and provided rich and diverse datasets for model training.^32–35,37^

mRNA display is an in vitro selection platform that couples the genotype and the phenotype through a covalent linkage between each peptide and its encoding mRNA.^38–40^ Because of its cell-free workflow, the theoretical diversity of this technique can reach up to 10^13^, substantially beyond the scope of in vivo technologies. As such, mRNA display has become a powerful platform that enables ultra-high-throughput evaluation of combinatorial peptide libraries for enzyme activity profiling and de novo binder discovery.^33,37,41,42^ The platform has also been adapted to inform on the substrate tolerance of several RiPP biosynthetic enzymes in a high throughput _manner._32,33,35,37,41,42

The previous examples of using mRNA display for developing predictive models of substrate preference have focused on RiPP biosynthetic enzymes that have evolved to make a natural product with a specific bioactivity. These types of enzymes that are the results of specific evolutionary selection have been termed specialist enzymes.^43,44^ In the context of RiPP biosynthetic enzymes, they have proven to have well defined substrate preferences and ML models are highly predictive in regards to acceptance of non-native substrates.^32,33^ ProcM and ProcM-like enzymes in contrast appear to have evolved to maintain high substrate tolerance, with this enzyme class demonstrating very high sequence conservation across the oceans whereas the substrate sequences are extremely diverse.^45^ They constitute an unusual example of a generalist enzyme^43^ that is maintained in the generalist state, with the mechanism by which this is achieved still unclear.^46^ How well mRNA display and ML will work to predict the substrate tolerance of such enzymes is an open question.

To understand the unusual flexibility of ProcM, we sought to develop a high-throughput platform that can systematically profile the inherent rules that govern any substrate preference of ProcM, which, in the long run, could guide the design of macrocyclic peptides from linear precursor peptides. In this study, mRNA display was used to generate a curated dataset of both positive substrates (cyclized) and negative substrates (non-cyclized) (**Figure 1B**). Then ML was used on the resulting datasets to generate a model that can classify the reactivity of a linear sequence. Overall, the trained model proved to have balanced accuracy in forecasting the reactivity of a precursor peptide as it needs to successfully overcome severe biochemical and mathematical constraints inherent to generalist enzyme fitness landscapes. This study shows that ProcM does not have unlimited ability for cyclization and that this generalist enzyme has preferences, but that these preferences are much less well defined than for other enzymes that have been investigated by mRNA display. Based on the findings of this investigation, we anticipate that the current learnable signal can be further improved through an expanded library design and advanced selection assays.

## Results and Discussion

### Establishment of an mRNA Display Method for ProcM Profiling

ProcM and ProcM-like enzymes pose several challenges with respect to profiling substrate specificity. First, these enzymes are unusual amongst lanthipeptide synthetases in that they naturally form rings in both directions, that is by catalyzing the reaction of Cys residues with dehydroamino acids that are located both up and downstream in the peptide sequence. The enzyme also naturally forms rings that vary greatly in size. When more than one ring is formed, the first ring determines the formation of subsequent rings. Finally, the most important property of ProcM in terms of potential utility is its macrocyclization capability, but this macrocyclization first requires an ATP-dependent dehydration step (Figure 1A) in which Ser/Thr side chains are phosphorylated followed by phosphate elimination. These features make it difficult to design a strategy that provides information on all aspects of ProcM catalysis, and in this first study we made certain choices that necessarily limit the information obtained but that allowed us to obtain a first glimpse into substrate preference.

We designed the library according to several criteria. Previous work on ProcA2.8 variants showed that randomizing residues within a ring can change the outcome of (methyl)lanthionine formation.^31^ When two rings are present, mutations in one of the rings can completely change the overall ring pattern.^28,31,47^ Because it is difficult to design a discriminator capable of distinguishing among ring topologies in a high-throughput manner, we elected to reduce the competing cyclization possibilities in the library design. Therefore, we decided to avoid the presence of any additional Cys residues in the randomized region of the substrate to guarantee formation of a single-ring pattern. We also considered removing both Ser and Thr to avoid any complications of potentially forming mixtures of ring patterns.^47^ However, the degenerate codons that remove both residues (NTT, NTK, or VWT) would result in only 4-8 amino acids in the library, which each would exclude several important physicochemical properties (NTT: IVFL; NTK: IVFLM; VWT: IVFLYHND). We therefore chose to remove only one of these two residues while also excluding the incorporation of stop codons. Based on these criteria, we chose the VDS degenerate codon, which encodes 13 residues (Asp, Glu, His, Lys, Arg, Ile, Leu, Asn, Gln, Val, Ser, Gly, and Met). We could have chosen the VMA codon, which would keep Thr but not Ser, but it would encode for 12 amino acids including Pro, which we considered less useful since Pro is believed to facilitate lanthionine/methyllanthionine ring formation in defined sequence contexts (e.g. by stabilizing turns through hydrogen bonding^48,49^). While these choices necessarily limit the information obtained, the set of amino acids contains positively and negatively charged residues as well as hydrophobic amino acids and covers a range of steric requirements. Considering that ProcA2.8 has been the model system for multiple prior bioengineering studies,^28,30,31^ we chose ProcA2.8 as the prototype to design our single-ring libraries. Accordingly, we generated three libraries named Ring 5, Ring 6, and Ring 7, with a sequenced diversity of 2.2*10^7^, 2.6*10^7^, and 1.7*10^7^ (**Figure 2A**, **Table S2**). These libraries cover three commonly observed ring sizes in natural lanthipeptides and investigate cyclization of Cys onto an upstream Dha (Ring 5) as well as a downstream Dha (Ring 6 and 7).

**Figure 2.**
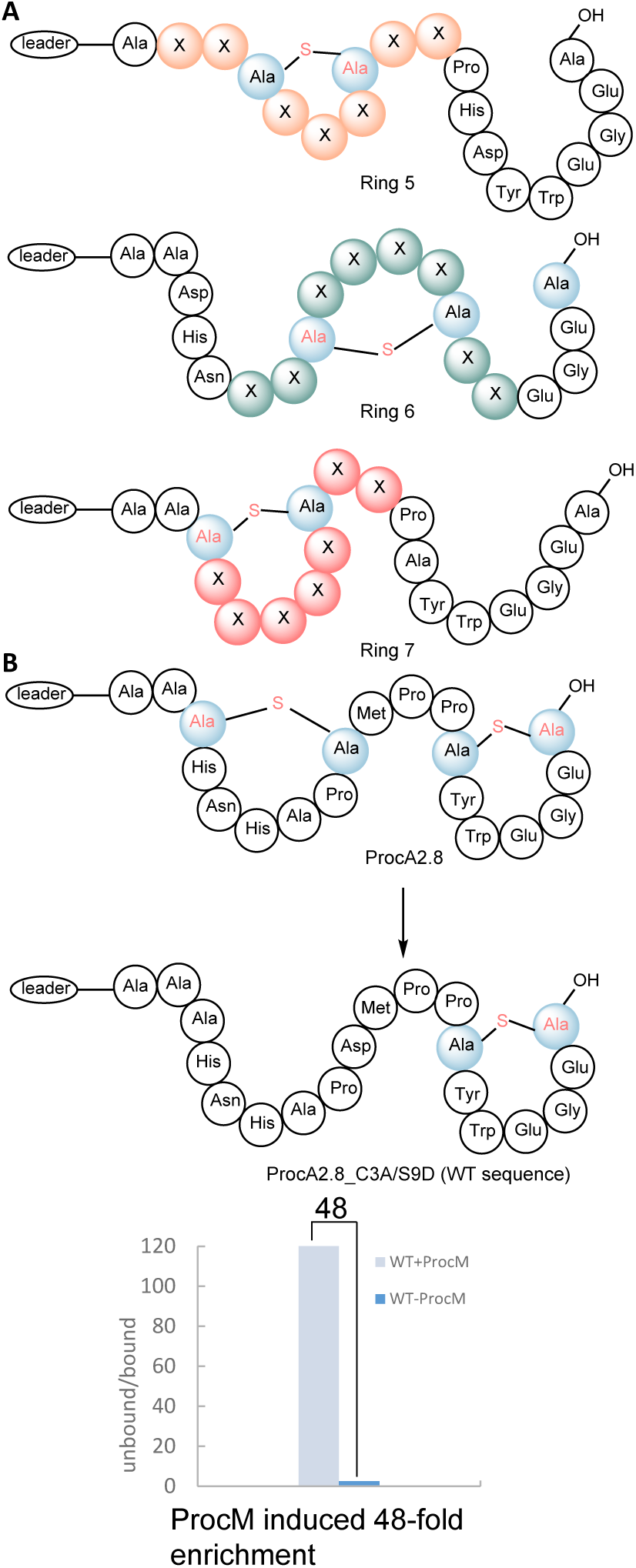
(A) Library design. Three single-ring pattern libraries, Ring 5, Ring 6, and Ring 7, were constructed and attached to the ProcA2.8 leader peptide. X denotes any of the 13 amino acids encoded by the VDS degenerate codon. (B) A WT-derived single ring sequence (ProcA2.8_C3A/S9D) was displayed as a positive control.

Lanthionine cyclization is a single enzyme-catalyzed two-step process involving two different residues on the substrate (Ser/Thr and Cys) and two active sites. Whereas more complicated selection schemes could interrogate both steps individually, for this first study we elected to use a selection strategy that would only report on sequences that are good substrates for obtaining cyclized peptides (Fig. 1B); similar approaches have worked well for other RiPP biosynthetic enzymes that catalyze multi-step processes.^32,41^ The strategy in Fig. 1B separates sequences that are poor substrates for either the dehydration or cyclization step (or both) from sequences that are good substrates for both steps, information that is most important in terms of future design of cyclic peptide libraries. In this workflow, uncyclized peptides will have a free Cys that can be modified by a biotinylated iodoacetamide (biotin-IAA). Those peptides are bound to streptavidin (SAv) beads that separate them from the cyclized peptides. Following this design, the selection focused on good substrates while the antiselection enriched for those peptides with poor reactivity for either enzymatic step (**Figure 1B**).

We first established a standard selection workflow with a peptide sequence based on a good natural substrate analog generating a single ring, ProcA2.8_C3A/S9D. Because of the lack of a readily available negative control substrate for ProcM that met our design principle, we only tested the positive control substrate showing good enrichment of cyclized over non-cyclized peptides (**Figure 2B**). Our library constructs were designed to contain an N-terminal HA tag for peptide purification, a ProcA2.8 leader peptide, followed by a core peptide. The displayed peptide library was treated with 3 μM ProcM, 0.1 μM TCEP, 5 mM ATP, and 5 mM MgCl2 in HEPES buffer (pH 7.5) for 2 h at room temperature. After anti-HA tag purification, the library was subjected to the biotin-(PEG)3-IAA assay (preincubation with 5 mM TCEP on ice for 30 min before adding 10 mM biotin-(PEG)3-IAA) at room temperature for 5 h, followed by acetone precipitation to remove any free biotin probe,^32^ which will interfere with the selection. After streptavidin (Sav) pulldown to isolate the bound fraction from the unbound fraction, the corresponding DNA was quantified by qPCR. In parallel, a ProcM-free negative control was not treated with ProcM but with all other reaction components.

With the established selection process, we performed five rounds of selection and antiselection for each library following the standard conditions described. We increased the selection stringency in the last round by incubating the library with ProcM for 30 min (instead of 2 h), while the reaction time of the antiselection was kept constant. Then the data were processed as discussed in the next section. Unlike previously reported RiPP biosynthetic enzymes profiling studies,^32^ we observed several differences in our selection. First, a continuously increased enrichment was not observed (when the enrichment is defined as the ratio of DNA recovery in the experiment versus the negative control). Instead, the enrichment fluctuated within a small range in both selection and antiselection. Second, after the first round of selection, a modest peptide level convergence was observed, as indicated by the normalized Shannon entropy (Hdataset, a measurement of diversity of peptide sequences,^50^ see SI equation 1), suggesting a portion of peptides was enriched from the naïve library. Beginning from the second round, the peptide level convergence became constant (Figure S2). We then analyzed the amino acid level convergence by computing a Log2Y* score (**Figure 3B**, see SI equation 2, Y* is an indicator of enrichment of a particular amino acid in the peptide). We plotted the heatmap of 13 residues against variable positions. In the Ring 7 library, although Cys and Ser were fixed at positions 3 and 9, Ser was enriched within the cyclization segment. We hypothesize that the possibility to generate smaller rings in the fixed 7-amino acid ring size may explain this observation; productive formation of smaller rings is more likely in larger ring constructs such as Ring 7 than in smaller ring constructs (Ring 5, 6) since 2- and 3-residue rings are seldom observed in natural lanthipeptides. Another clear feature is the observation that charged residues are disfavored when flanking the dehydration site of Ser4 in Ring 5, Ser13 in Ring 6, and Ser9 in Ring 7. The negatively charged residues Glu and Asp are particularly depleted at these positions. In Ring 6, Arg, Lys, and Gly are slightly enriched at positions flanking the invariant Ser residue in the antiselection. By contrast, the Cys-adjacent residues did not display strong enrichment. Overall, different ring sizes exhibited distinct behaviors under identical selection stringency. In particular, the preferences of positive and negative sequences were more prominent in the Ring 7 system, whereas this distinction was less evident in the Ring 5 and Ring 6 systems.

**Figure 3.**
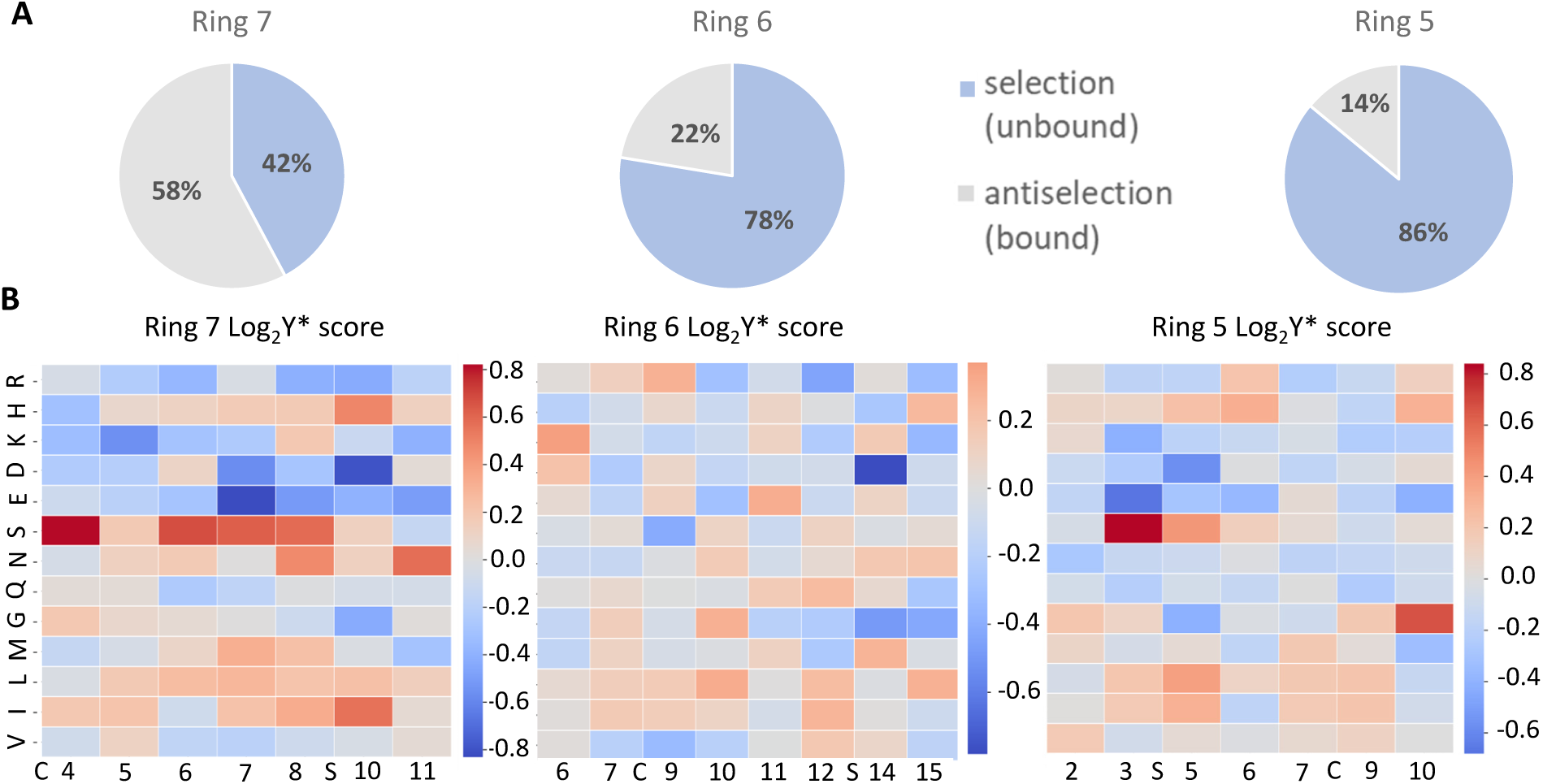
(A) Dataset distribution after five rounds of selection and after data processing as described in the text. For the data in this figure, peptides containing multiple Ser residues were included. The blue pool constitutes cyclized peptides and the grey pool non-cyclized peptides. (B) Amino acid level enrichment at specific positions of three libraries represented by Log2Y* score (see SI for details). Red fields indicate amino acids that are enriched at a certain position in the cyclized peptide pool and blue fields indicate residues that are depleted at specific positions in the cyclized peptide pool.

The distinct observations in ProcM profiling could be caused by multiple factors. In general, we obtained fewer negative peptides for ProcM even after five rounds of selection/antiselection across three different libraries (**Figure 3A**). This observation is not unexpected for an enzyme for which the evolutionary pressure has been for substrate tolerance rather than substrate specificity.^46^ Furthermore, in a typical mRNA display interrogation of substrate specificity, enzyme is in excess over substrate, which for an intrinsically substrate-tolerant enzyme like ProcM may result in even moderate substrates ending up in the good substrate pool. Our libraries contain a focused set of 13 residues, without Tyr, Phe, Pro, Trp, Ala, Cys, and Thr, which introduces some bias in the substrates. For instance, incorporation of Pro has been reported to facilitate cyclization in some sequence contexts,^48,49^ and leading to unsuccessful cyclization in other sequence contexts.^31^ Furthermore, our choice to include Ser but exclude Thr favors nonenzymatic cyclization involving Dha.^51^ Although this background reaction is not kinetically competitive with enzyme catalysis, it is sequence dependent and could contribute to cyclization in the mRNA-display conditions. Such an intrinsic chemical background process complicates the interpretation of the present selection data and may partially explain why our system did not show the smooth, continuous enrichment as reported for other enzymes.^32,33^ In particular, sequence-dependent nonenzymatic cyclization could partially obscure the distinction between enzyme-driven selection and background recovery. Preventing this background reactivity is difficult, and determining the extent of background reactivity would require a separate catalyst with just dehydration activity that displays the same sequence preference, which currently is not available. Efforts to develop the tools to address this limitation are ongoing.

We emphasize that the results across the three libraries cannot be compared directly with each other. In other words, the higher observed number of selected over non-selected peptides for the Ring 7 library does not mean that such rings are easier (faster) to form. In fact, kinetic studies in which two rings could form competitively have consistently demonstrated that the smaller rings are formed faster.^47^ Regardless of the complications for the system at hand to investigate enzyme selectivity, in terms of practical applications, successful prediction of sequences that are macrocyclized efficiently, is a valuable outcome.

### Machine Learning Analysis of Substrate Compatibility

We then investigated whether the mRNA display selection datasets contained learnable information with respect to position-specific enrichment or depletion of amino acids. In the workflow used below, the selection output was used for a binary classification.

### Data Preprocessing

The dataset for model training was compiled from the outcome with the three mRNA display libraries, Ring 5, Ring 6, and Ring 7. Prior to model training, the raw sequence data went through a two-stage preprocessing pipeline to ensure sequence quality and consistency (**Figure 4A**). In the first stage, duplicate sequences were removed, as well as sequences with multiple Ser residues that could lead to alternative ring patterns that would complicate analysis (these latter sequences were included in Figure 3 but removed for the ML analysis). This process reduced the dataset from 488,007 to 416,949 unique sequences. In the second stage, sequences containing mutations outside the allowed positions defined by each library scaffold were excluded, yielding a final dataset of 371,446 sequences. Across all libraries, sequences were assigned to one of two classes: Class 0 (streptavidin-bound, uncyclized) and Class 1 (unbound, cyclized). The final processed dataset consisted of 65,965 bound sequences and 305,481 unbound sequences, resulting in a class imbalance ratio of 4.6. Among the individual libraries, Ring 7 exhibited the most balanced distribution (imbalance ratio = 0.6), whereas Ring 5 showed the greatest imbalance (imbalance ratio = 10.4).

**Figure 4.**
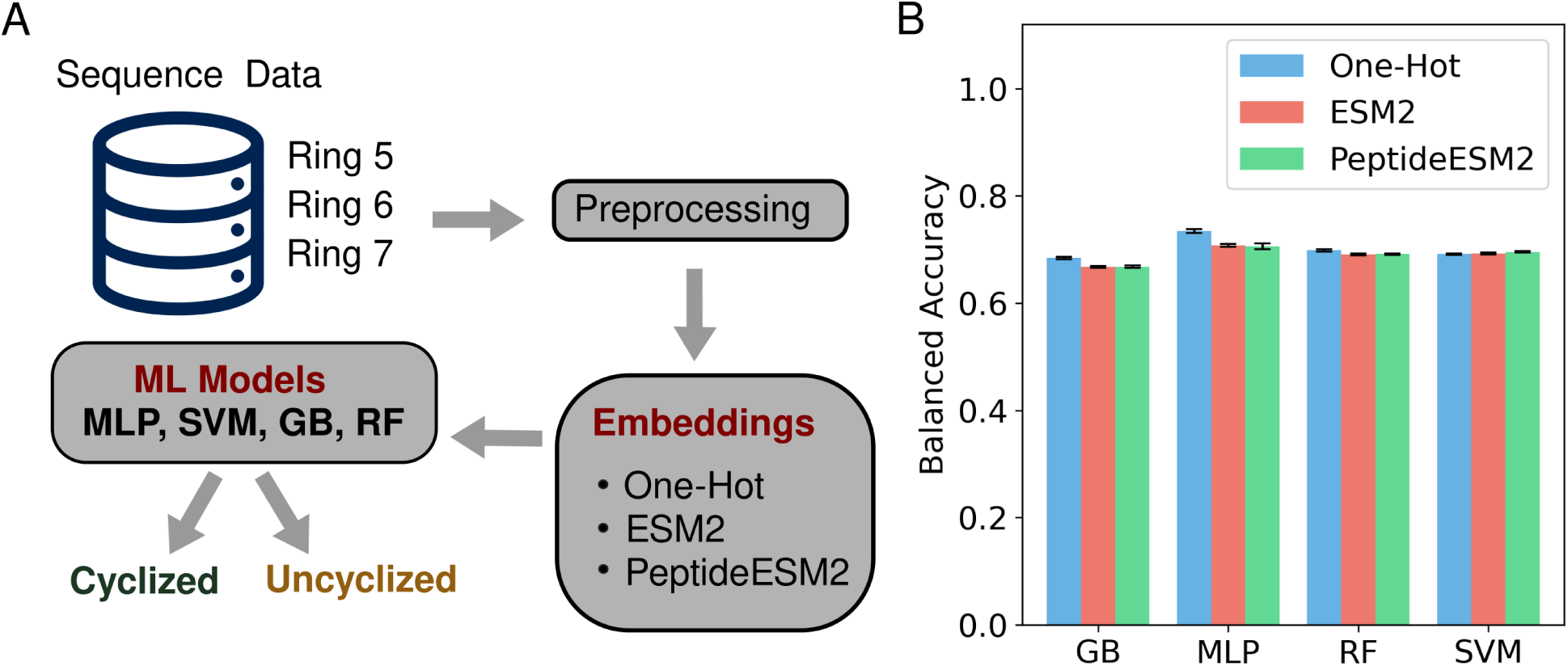
(A) Data processing and model training workflow. A variety of ML models were evaluated. MLP: Multi-Layer Perceptron, SVM: Support Vector Machine, GB: Gradient Boosting, RF: Random Forest. (B) Model performance evaluation across all classifiers and embedding combinations.

### Model Architecture and Featurization

To classify peptide sequences as cyclized or uncyclized, four machine learning classifiers were evaluated: Gradient Boosting (GB), Random Forest (RF), and Support Vector Machine (SVM) using the scikit-learn library,^52^ and a custom Multilayer Perceptron (MLP). As illustrated in **Figure 4A**, each classifier was trained using three distinct sequence representation strategies: one-hot encoding, ESM2 protein language model embeddings,^53^ and PeptideESM2 embeddings,^34^ a peptide-specialized variant of ESM2 fine-tuned on peptide sequence data. This framework enabled a systematic comparison of both conventional positional encodings and language model-derived embeddings across multiple classifier architectures. In this study, unbound sequences were designated as the positive class because they correspond to cyclized peptides in the mRNA display workflow. To mitigate the effects of class imbalance during neural network training, the MLP model was optimized using a Focal loss^54^ formulation.

### Model Performance

Model performance across all classifiers and embedding combinations is summarized in **Figure 4B** and **Table S1**. Balanced accuracy values ranged from approximately 0.67 to 0.73, while AUROC (Area Under the Receiver Operating Characteristic) values ranged from 0.74 to 0.79. Among all evaluated approaches, the MLP combined with one-hot encoding achieved the strongest overall performance, yielding a balanced accuracy of 0.73, an AUROC of 0.79, and the highest Matthews correlation coefficient (MCC) of 0.47. Because unbound sequences were defined as the positive class, sensitivity reflects the model’s ability to identify cyclized peptides correctly. Sensitivity values remained consistently high across all models (0.85-0.95), whereas specificity, corresponding to the identification of uncyclized peptides, was comparatively lower (0.38-0.56). This disparity likely arises from the substantial class imbalance within the combined dataset, which contains nearly five cyclized sequences for every uncyclized sequence. The model performance for this ProcM dataset lies within the expected range of performance metrics reported by ProteinGym^55^ benchmark for protein fitness prediction using supervised ML models.

Notably, one-hot encoding consistently matched or outperformed both ESM2 and PeptideESM2 embeddings across all classifier architectures. These results suggest that the fixed-length, position-specific characteristics of the cyclic peptide libraries are effectively captured using simple positional encodings, limiting the additional benefit provided by more complex language model-based representations. Furthermore, ESM2 and PeptideESM2 exhibited highly similar performance across all classifiers, indicating that peptide-specific pretraining conferred minimal advantage in this particular classification setting.

Pretrained language models like ESM2 are optimized on billions of diverse full-length natural protein sequences to learn global evolutionary relationships and three-dimensional structural folded constraints. When forced to embed short, synthetically randomized peptide core libraries, these models project the short peptide into a massive 1280-dimensional embedding space, introducing unnecessary noise. One-hot encoding avoids this projection error, preserving the exact, low-dimensional information needed for simple classification. Finally, generalist enzymes naturally accept a wide variety of amino acids at mutated positions, meaning the dataset has a low fraction of highly variable sites. Vieria et al. have shown that intrinsic dataset features such as fraction of highly variable sites determine the prediction accuracy of supervised ML models for variant effect predictors.^56^ In the datasets such as those obtained for ProcM where individual residue mutations do not change the fitness significantly, there is no complex epistatic relationship that needs to be learned by a transformer architecture. When the underlying sequence-function landscape lacks sharp mutational signatures, attention mechanisms fail to beat single site information provided by a simple one-hot encoding.

### Experimental Validation

To assess whether the classifier captured meaningful biochemical reactivity, we selected 60 representative sequences spanning the full range of model scores from an in silico-generated validation peptide dataset of 10^5^ sequences for experimental validation (**Figure 5A**). These peptides were produced by cell-free protein synthesis and then incubated with ProcM under the same reaction conditions used for the positive selection in rounds 1-4. Prediction accuracy was evaluated using a binary threshold of 0.5. Both classifier scores and experimental modification ratios below 0.5 were assigned as unmodified, whereas values above 0.5 were assigned as modified. A prediction was considered correct when the predicted and experimental binary classifications agreed. This classification provides a simple qualitative comparison but is inherently a very rough estimation for a system with continuous modification levels.

**Figure 5.**
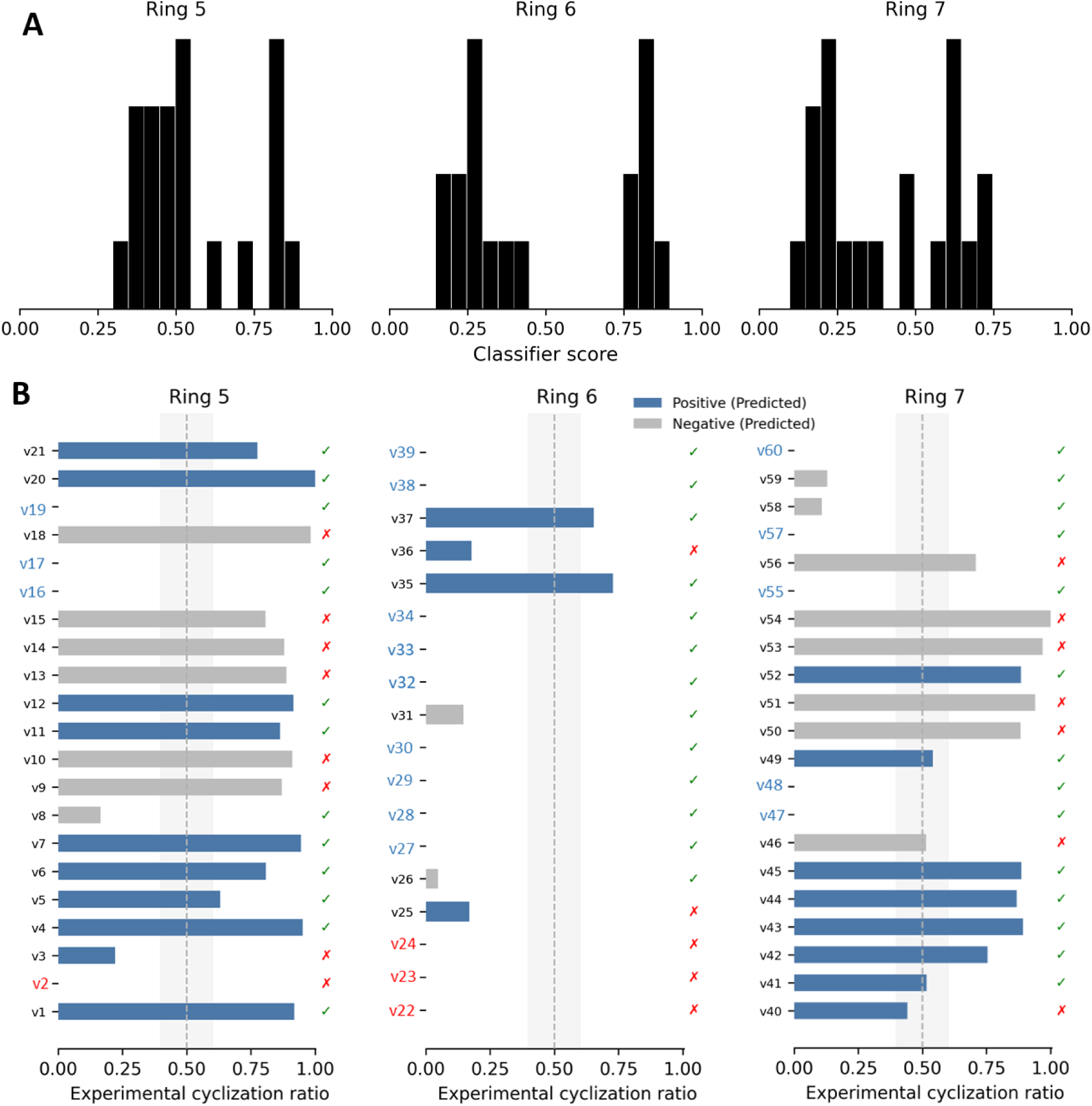
(A) Predicted score distribution of the validation dataset. The X-axis represents the predicted classifier score, while the bar height represents the relative fraction of peptides within each score range. (B) Experimental validation results of randomly sampled peptides v1-v60. The dashed line indicates a median modification ratio of 0.5. Blue bar: predicted cyclized peptides and observed cyclization ratio. Grey bar: predicted non-cyclized peptides and the observed cyclization ratio. Green checkmarks show true positives or true negatives; red crosses indicate false positives or false negatives. For peptides with an experimental cyclization ratio of 0, if this observation was consistent with the model prediction, the peptide number was colored blue on the left (e.g. v19); otherwise, it was colored red (e.g. v2). The experimental cyclization ratio was determined by LC-MS.

The overall experimental validation accuracy was 67%, which was in good agreement with the model’s balanced accuracy of 67-70% (**Figure 6A**). This consistency suggests that, despite its modest performance, the classifier captures a real and reproducible sequence-level signal in the selection data. Importantly, the validated peptides did not show a broad range of reactivity. Instead, most reactive sequences showed substantial to near-complete modification, whereas most nonreactive sequences remained largely unmodified (**Figure 5B**). This behavior, together with the lack of an additional discriminator capable of resolving more detailed reaction outcomes, suggests that the current dataset is better suited for binary classification than for regression-based modeling of relative substrate fitness. Accordingly, we interpret the model output as a measure of assay-defined substrate compatibility rather than a quantitative predictor of intrinsic ProcM reactivity. As noted above, in terms of practical applications of generating high-quality cyclic peptide libraries, this model output is still beneficial.

**Figure 6.**
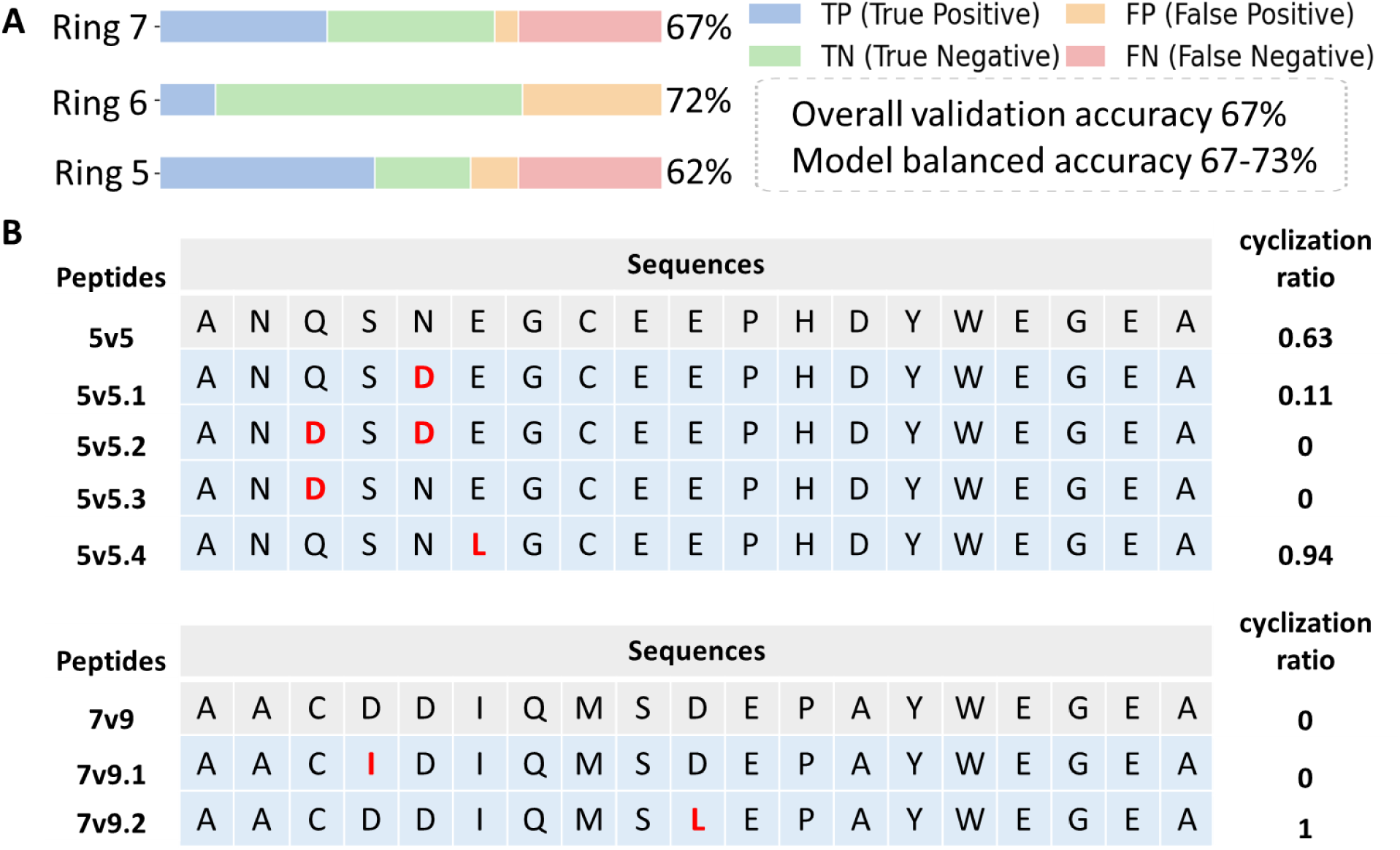
(A) Analysis of validation accuracy by ring size. (B) Evaluation of local feature predictions by single-point mutational studies. The cyclization ratio was determined by LC-MS.

When the validation results were further examined by ring size, clear differences emerged (**Figure 5B** and **6A**). Ring 5 peptides were predominantly cyclized, with modification ratios generally falling between 0.63 and 1.00, yet this subset showed the lowest validation accuracy at 62% and a relatively large number of false negatives, consistent with the strong class imbalance in the Ring 5 system (**Figure 3A** and **6A**). The Ring 7 subset showed more heterogeneous outcomes, including fully cyclized products, products that were dehydrated without cyclization, and peptides with no detectable dehydration (see SI). However, its overall agreement with the model remained comparable to that of the Ring 6 subset, approximately 67-72% (**Figure 6A**). By contrast, the Ring 6 system behaved differently from expectation. Although this ring size was initially thought to contain a broader set of modifiable substrates given the common occurrence of rings of six amino acids in natural substrates, only a small number of peptides were true positives, whereas the true negatives were better predicted (**Figure 5B** and **Figure 6A**). Taken together, these different observations suggest that the uneven model performance across ring sizes reflects that no single universal rule governs ProcM reactivity, but also highlight the limitations in the training data, including class imbalance and the relative underrepresentation of negative sequences. Biochemically, ProcM catalyzes a two-step reaction, while our selection assay only captures the cyclization step without informing on the dehydration step. Thus, to investigate the individual steps for a complex lanthipeptide synthetase like ProcM, a more advanced selection assay could further improve the outcome.

We next assessed whether the selection data contained residue-level information that could be used to guide local modulation of ProcM reactivity. Although the classifier showed only modest predictive performance, the amino acid level enrichment heatmap for the Ring 5 library in **Figure 3** revealed a clear pattern, with Asp and Glu highly enriched in the antiselection dataset at flanking positions of Ser4. To examine whether this trend was experimentally meaningful, we selected 5v5 as a representative peptide for targeted mutagenesis (**Figure 6B**). Peptide 5v5 (v5 in Figure 5) was validated as a true good substrate with a moderate cyclization ratio of 0.63. Mutation of Asn5 to Asp in 5v5.1 significantly reduced the cyclization ratio to 0.11, whereas simultaneous mutation of both flanking residues, Gln3 and Asn5 in 5v5.2, to Asp completely abolished cyclization. Notably, substitution of Gln3 with Asp alone also eliminated detectable product formation for 5v5.3, with no observable cyclization or dehydration intermediate. These results suggest that negatively charged residues adjacent to the reactive Ser are disfavored in this sequence context, with the dehydration step likely most affected. At the same time, the distinct outcomes produced by substitution on the N-terminal or C-terminal side of Ser indicate that the positional effect is not strictly equivalent. In contrast, when Glu6 was changed to a neutral residue, Leu6, the cyclization product was predominant in 5v5.4. Similarly, peptide 7v9 (v48 in Figure 5) was classified as a negative substrate with an experimental validation of 0 for the cyclization ratio. When the Ser-flanking residue Asp10 was mutated to Leu10 in 7v9.2, complete cyclization was observed instead. In contrast, the cysteine-flanking residue Asp4 does not appear to significantly contribute to the observed lack of cyclization, because the Ile4 mutant (7v9.1) remained uncyclized. This focused mutational analysis suggests that, despite the limited predictive accuracy of the classifier model, the local sequence features captured by the selection still provide useful guidance for probing ProcM reactivity. The observed depletion of Asp/Glu residues flanking Ser residues in successfully cyclized substrates in this mRNA display study is consistent with previous analysis of the flanking residues in naturally occurring class II lanthipeptides.^57,58^

### Comparison with previous investigations of RiPP biosynthetic enzymes by mRNA display

Previous studies have used the approach applied here to investigate the substrate specificity of the glutamyl-tRNA dependent dehydratase LazBF and the Cys cyclodehydratase LazDEF.^32^ Like ProcM, these enzymes have multiple active sites and catalyze multi-step processes, which has not prevented highly successful machine learning predictions. Even though the Laz enzymes have undergone natural selection to generate the structurally complex natural product lactazole that is a member of the thiopeptide antibiotics, they have been shown to display high substrate tolerance and have been used to generate thiopeptide libraries.^37,59,60^ Despite their substrate tolerance, the evolutionary pressure on these enzymes has likely optimized their activity towards the natural substrate peptide sequence. In turn, this evolutionary path has likely endowed these enzymes with well-defined substrate preferences for non-native substrates. As a consequence, the models generated by ML from the mRNA selection data for both enzymes had very high prediction accuracy (>99%).^32,37^ Clark et al. have showed that using domain-specific masked language modeling enables the construction of highly accurate LazDEF substrate specificity models even in extreme data-scarce scenarios (N∼100).^34^ This observation is aligned with the expected behavior of specialist enzymes that possess sharp sequence-function rules which allows AI models to reach near-perfect accuracy quickly. Generalist enzymes such as ProcM have evolved for extreme structural tolerance. This tolerance creates severe dataset class imbalances reported in this study along with low within-site variation that suppresses predictability. Therefore, the prediction accuracy reported in this study likely reflects the nature of sequence-function relationship rather than model or dataset limitations. To establish directly why the ProcM dataset is more challenging for classification than the Laz datasets, we compared the libraries using four statistics calculated from amino acid composition alone. These metrics measure how strongly substrates and non-substrates differ in sequence. They therefore provide a model-independent estimate of how much discriminatory information is available to a classifier. First, we calculated the Jensen–Shannon (JS) divergence between substrate and non-substrate residue distributions at each variable position. A value near zero indicates similar residue preferences in the two classes. A larger value indicates that residue identity helps distinguish substrates from non-substrates. The average JS divergence is near zero for all three ProcM libraries and more than an order of magnitude larger for the Laz libraries (Fig. 7A). Thus, residue identity at a given position carries little information about ProcM substrate acceptance, whereas the Laz datasets contain stronger position-specific signals. We next calculated the mean absolute log2 enrichment across all position–residue combinations. This metric measures how strongly individual residues are enriched or depleted among substrates. Larger values indicate stronger residue-level rules and greater discriminatory power. A typical residue is favored by only about 1.1-fold in ProcM substrates, compared with roughly 1.8-fold in the Laz libraries (Fig. 7B). ProcM therefore shows much weaker residue-specific preferences. We then compared within-class and between-class sequence distances. The distance was defined as the expected fraction of positions at which two independently drawn sequences differ. If substrates and non-substrates occupy distinct regions of sequence space, between-class distances should differ from within-class distances. Similar distances instead indicate strong overlap between the two classes. For ProcM, a substrate is about as dissimilar from a non-substrate as it is from another substrate. In contrast, the Laz libraries show consistent separation between the two classes (Fig. 7C). Thus, ProcM substrates and non-substrates are intrinsically less separable in sequence space. Finally, we measured how much sequence diversity remains after selection using the ratio of substrate entropy to the entropy of the full observed library. A ratio near unity indicates that substrates retain nearly the full diversity of the library. A lower value indicates stronger sequence restriction. This ratio remains near unity for ProcM but is lower for Laz (Fig. 7D). ProcM therefore accepts sequences across a much broader portion of the available sequence space. Together, these measures show that ProcM substrates and non-substrates are only weakly separated in sequence space compared with the Laz datasets. ProcM shows limited positional preferences, weak residue enrichment, substantial overlap between classes, and broad retention of sequence diversity. These features produce a more diffuse sequence–function landscape, with fewer clear boundaries between reactive and non-reactive sequences. As a result, the ProcM dataset contains less discriminatory signal for an ML classifier to learn. In contrast, the stronger sequence constraints of the specialist Laz enzymes create sharper class boundaries and a more readily learnable sequence–function relationship.

**Figure 7.**
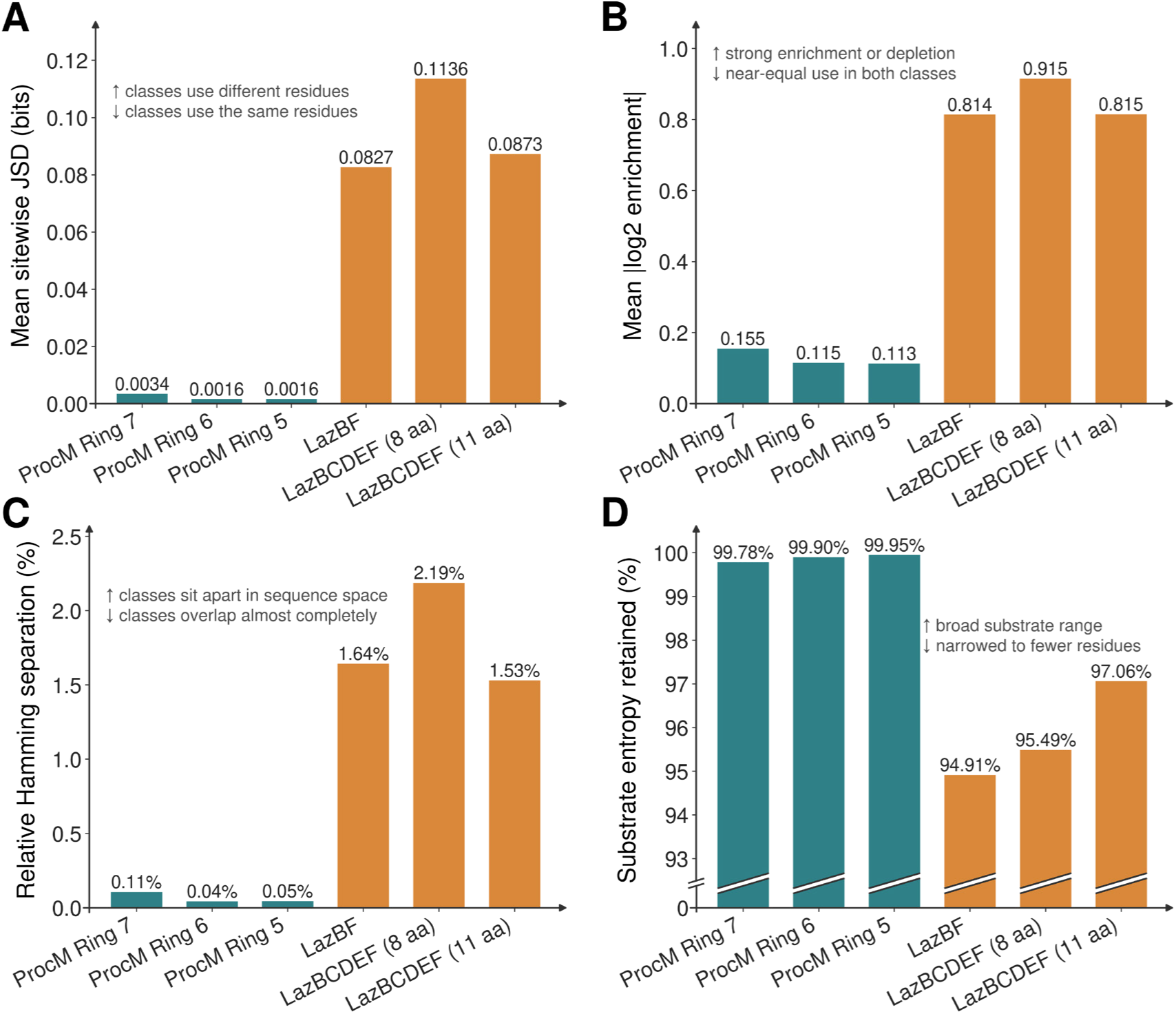
Sequence composition of substrates and non-substrates in the ProcM (teal) and Laz (orange) libraries. (A) Mean sitewise Jensen-Shannon divergence between the two residue distributions. (B) Mean absolute log2 enrichment per position and residue. (C) Excess between-class Hamming distance relative to the within-class mean. (D) Substrate entropy relative to the full library.

ProcM and other ProcM-like enzymes in cyanobacteria have been shown to have extreme divergence in the core peptide sequences of their natural substrates (e.g. Figure S1).^6,45^ In contrast, the ProcM-like enzymes have very high sequence identity across different species. These observations and others have been interpreted as strong purifying evolutionary pressure to retain the high substrate tolerance of the enzymes and at the same time maximally diversify the structures of their products.^45^ Although the functions of the resulting prochlorosins are still unclear, a suggested role is to serve as a smoke screen to prevent pattern recognition by potential predators.^46^ If correct, high conversion of the diverse sequences into single products is likely not required, and possibly less advantageous than generation of mixtures.

One of the consequences of ProcM-like enzymes having to convert thousands of ProcA peptides with diverse sequences in the marine environment^45^ is that they are much slower than other class II lanthipeptide synthetases because they have not evolved activity for individual substrate sequences.^61^ Another consequence is that they do not guide product formation, but that instead the substrate peptide sequences determine the outcome of catalysis.^28,47^ In this study, additional consequences are revealed. Compared to other RiPP enzymes that have been investigated by mRNA display, the extreme substrate tolerance of ProcM resulted in a very difference balance between the number of substrates that are cyclized and the number of poor substrates. The high substrate tolerance also resulted in a much less well-defined boundary between substrates and non-substrates. These findings reinforce the conclusion from a recent kinetic study that enzymes like ProcM may not be the ideal enzymes for cyclic peptide generation that they were once believed to be. Several studies with RiPP biosynthetic enzymes that evolved to make single bioactive peptides have demonstrated that their inherent substrate tolerance imparted by the proximity-driven activity of leader peptide dependent RiPP biosynthetic enzymes is sufficient for the generation of cyclic peptide libraries.^20,37,60,62^ Although some cyclic peptide libraries generated by ProcM were shown to have a very high success rate when based on natural substrates,^30^ for other libraries the extreme substrate tolerance like that displayed by ProcM may be an impediment for high-quality library generation rather than an advantage.

### Conclusions

In this study, we designed a high-throughput selection workflow to map the substrate selectivity of ProcM using mRNA display. Unlike similar investigations of previous specialist enzymes, this study constitutes the first analysis of a generalist enzyme using this technology. We initially designed three different ring library scaffolds to obtain insights into such a generalist enzyme with the intention to follow up with additional scaffolds to finetune the models. The resulting datasets were integrated with a deep learning approach to build a binary classifier that can distinguish a modifiable precursor peptide from an unmodifiable peptide with modest predictability. Because of class imbalance that may be a general problem for generalist enzymes and library sizes that were intentionally limited for the three different ring patterns, the overall model performance is moderate and less accurate than models generated previously for specialist enzymes. The selection did generate some learnable features that can be used to inform future cyclic peptide design, but it also revealed special challenges that may be common for generalist enzymes. Important lessons were learned for investigating this type of enzyme, and from a practical perspective the current findings will aid in cyclic library design. We anticipate that a larger library, for example, using the NNK degenerate codon, and a selection assay that can distinguish between poor dehydration and poor cyclization substrates may produce more detailed information for machine learning on individual steps in future studies.

## Supporting information

Primer sequences

Experimental validation data

## ASSOCIATED CONTENT

### Supporting Information

The Supporting Information is available free of charge at https://pubs.acs.org/doi/Experimental procedures, Figures S1-S4 showing the selection outcomes and mass spectra of peptides used for validation, and Tables S1-S2 with evaluation matrices and library dimensions (PDF).

## Notes

The authors declare no competing financial interests.

## Data availability

Raw data are deposited on Mendeley and will be released upon publication. Ouyang, Yao; van der Donk, Wilfred (2026), “Data associated with “Mapping the sequence preference of class II lanthipeptide synthetase ProcM by mRNA display””, Mendeley Data, V1, doi: 10.17632/4nyxkvht7p.1

## ACKNOWLEDGMENTS

We thank Owen Ouyang in the research group of Professor Nicholas Wu, and Guthrie Stroh in the research groups of Professor Nicholas Wu and Professor Mitchell Douglas, for their help in establishing the protocol and helpful discussions. We thank Tony Zhang for his help in troubleshooting the script for data processing.

## References

(1) Repka, L. M.; Chekan, J. R.; Nair, S. K.; van der Donk, W. A. Mechanistic understanding of lanthipeptide biosynthetic enzymes. Chem. Rev. 2017, 117, 5457–520.

(2) Arnison, P. G.; Bibb, M. J.; Bierbaum, G.; Bowers, A. A.; Bugni, T. S.; Bulaj, G.; Camarero, J. A.; Campopiano, D. J.; Challis, G. L.; Clardy, J.; Cotter, P. D.; Craik, D. J.; Dawson, M.; Dittmann, E.; Donadio, S.; Dorrestein, P. C.; Entian, K. D.; Fischbach, M. A.; Garavelli, J. S.; Göransson, U.; Gruber, C. W.; Haft, D. H.; Hemscheidt, T. K.; Hertweck, C.; Hill, C.; Horswill, A. R.; Jaspars, M.; Kelly, W. L.; Klinman, J. P.; Kuipers, O. P.; Link, A. J.; Liu, W.; Marahiel, M. A.; Mitchell, D. A.; Moll, G. N.; Moore, B. S.; Müller, R.; Nair, S. K.; Nes, I. F.; Norris, G. E.; Olivera, B. M.; Onaka, H.; Patchett, M. L.; Piel, J.; Reaney, M. J.; Rebuffat, S.; Ross, R. P.; Sahl, H. G.; Schmidt, E. W.; Selsted, M. E.; Severinov, K.; Shen, B.; Sivonen, K.; Smith, L.; Stein, T.; Süssmuth, R. D.; Tagg, J. R.; Tang, G. L.; Truman, A. W.; Vederas, J. C.; Walsh, C. T.; Walton, J. D.; Wenzel, S. C.; Willey, J. M.; van der Donk, W. A. Ribosomally synthesized and post-translationally modified peptide natural products: overview and recommendations for a universal nomenclature. Nat. Prod. Rep. 2013, 30, 108–60.

(3) Montalbán-López, M.; Scott, T. A.; Ramesh, S.; Rahman, I. R.; van Heel, A. J.; Viel, J. H.; Bandarian, V.; Dittmann, E.; Genilloud, O.; Goto, Y.; Grande Burgos, M. J.; Hill, C.; Kim, S.; Koehnke, J.; Latham, J. A.; Link, A. J.; Martínez, B.; Nair, S. K.; Nicolet, Y.; Rebuffat, S.; Sahl, H.-G.; Sareen, D.; Schmidt, E. W.; Schmitt, L.; Severinov, K.; Süssmuth, R. D.; Truman, A. W.; Wang, H.; Weng, J.-K.; van Wezel, G. P.; Zhang, Q.; Zhong, J.; Piel, J.; Mitchell, D. A.; Kuipers, O. P.; van der Donk, W. A. New developments in RiPP discovery, enzymology and engineering. Nat. Prod. Rep. 2021, 38, 130–239.

(4) Cotter, P. D.; Deegan, L. H.; Lawton, E. M.; Draper, L. A.; O’Connor, P. M.; Hill, C.; Ross, R. P. Complete alanine scanning of the two-component lantibiotic lacticin 3147: generating a blueprint for rational drug design. Mol. Microbiol. 2006, 62, 735–47.

(5) Islam, M. R.; Shioya, K.; Nagao, J.; Nishie, M.; Jikuya, H.; Zendo, T.; Nakayama, J.; Sonomoto, K. Evaluation of essential and variable residues of nukacin ISK-1 by NNK scanning. Mol. Microbiol. 2009, 72, 1438–47.

6. Li, B.; Sher, D.; Kelly, L.; Shi, Y.; Huang, K.; Knerr, P. J.; Joewono, I.; Rusch, D.; Chisholm, S. W.; van der Donk, W. A. Catalytic promiscuity in the biosynthesis of cyclic peptide secondary metabolites in planktonic marine cyanobacteria. Proc. Natl. Acad. Sci. U.S.A. 2010, 107, 10430–5.

(7) Caetano, T.; Krawczyk, J. M.; Mosker, E.; Süssmuth, R. D.; Mendo, S. Heterologous expression, biosynthesis, and mutagenesis of type II lantibiotics from *Bacillus licheniformis* in *Escherichia coli*. Chem. Biol. 2011, 18, 90–100.

(8) Field, D.; Molloy, E. M.; Iancu, C.; Draper, L. A.; PM, O. C.; Cotter, P. D.; Hill, C.; Ross, R. P. Saturation mutagenesis of selected residues of the alpha-peptide of the lantibiotic lacticin 3147 yields a derivative with enhanced antimicrobial activity. Microb. Biotechnol. 2013, 6, 564–75.

(9) Krawczyk, J. M.; Völler, G. H.; Krawczyk, B.; Kretz, J.; Brönstrup, M.; Süssmuth, R. D. Heterologous expression and engineering studies of labyrinthopeptins, class III lantibiotics from *Actinomadura namibiensis*. Chem. Biol. 2013, 20, 111–22.

(10) Zhang, Q.; Yang, X.; Wang, H.; van der Donk, W. A. High divergence of the precursor peptides in combinatorial lanthipeptide biosynthesis. ACS Chem. Biol. 2014, 9, 2686–94.

(11) Arias-Orozco, P.; Inklaar, M.; Lanooij, J.; Cebrián, R.; Kuipers, O. P. Functional expression and characterization of the highly promiscuous lanthipeptide synthetase SyncM, enabling the production of lanthipeptides with a broad range of ring topologies. ACS Synth. Biol. 2021, 10, 2579–91.

(12) Chatterjee, C.; Patton, G. C.; Cooper, L.; Paul, M.; van der Donk, W. A. Engineering dehydro amino acids and thioethers into peptides using lacticin 481 synthetase. Chem. Biol. 2006, 13, 1109–17.

(13) Levengood, M. R.; Knerr, P. J.; Oman, T. J.; van der Donk, W. A. In vitro mutasynthesis of lantibiotic analogues containing nonproteinogenic amino acids. J. Am. Chem. Soc. 2009, 131, 12024–5.

(14) Bosma, T.; Kuipers, A.; Bulten, E.; de Vries, L.; Rink, R.; Moll, G. N. Bacterial display and screening of posttranslationally thioether-stabilized peptides. Appl. Environ. Microbiol. 2011, 77, 6794–801.

(15) Zhou, L.; Shao, J.; Li, Q.; van Heel, A. J.; de Vries, M. P.; Broos, J.; Kuipers, O. P. Incorporation of tryptophan analogues into the lantibiotic nisin. Amino Acids 2016, 48, 1309–18.

(16) Montalbán-López, M.; van Heel, A. J.; Kuipers, O. P. Employing the promiscuity of lantibiotic biosynthetic machineries to produce novel antimicrobials. FEMS Microbiol. Rev. 2016, 41, 5–18.

(17) Kuthning, A.; Durkin, P.; Oehm, S.; Hoesl, M. G.; Budisa, N.; Süssmuth, R. D. Towards biocontained cell factories: an evolutionarily adapted *Escherichia coli* strain produces a new-to-nature bioactive lantibiotic containing thienopyrrole-alanine. Sci. Rep. 2016, 6, 33447.

(18) Urban, J. H.; Moosmeier, M. A.; Aumüller, T.; Thein, M.; Bosma, T.; Rink, R.; Groth, K.; Zulley, M.; Siegers, K.; Tissot, K.; Moll, G. N.; Prassler, J. Phage display and selection of lanthipeptides on the carboxy-terminus of the gene-3 minor coat protein. Nat. Commun. 2017, 8, 1500.

(19) Burkhart, B. J.; Kakkar, N.; Hudson, G. A.; van der Donk, W. A.; Mitchell, D. A. Chimeric leader peptides for the generation of non-natural hybrid RiPP products. ACS Cent. Sci. 2017, 3, 629–38.

(20) Hetrick, K. J.; Walker, M. C.; van der Donk, W. A. Development and application of yeast and phage display of diverse lanthipeptides. ACS Cent. Sci. 2018, 4, 458–67.

(21) Kakkar, N.; Perez, J. G.; Liu, W. R.; Jewett, M. C.; van der Donk, W. A. Incorporation of nonproteinogenic amino acids in class I and II lantibiotics. ACS Chem. Biol. 2018, 13, 951–7.

(22) Si, T.; Tian, Q.; Min, Y.; Zhang, L.; Sweedler, J. V.; van der Donk, W. A.; Zhao, H. Rapid screening of lanthipeptide analogs via in-colony removal of leader peptides in *Escherichia coli*. J. Am. Chem. Soc. 2018, 140, 11884–8.

(23) Schmitt, S.; Montalban-Lopez, M.; Peterhoff, D.; Deng, J.; Wagner, R.; Held, M.; Kuipers, O. P.; Panke, S. Analysis of modular bioengineered antimicrobial lanthipeptides at nanoliter scale. Nat. Chem. Biol. 2019, 15, 437–43.

(24) Fu, Y.; Xu, Y.; Ruijne, F.; Kuipers, O. P. Engineering lanthipeptides by introducing a large variety of RiPP modifications to obtain new-to-nature bioactive peptides. FEMS Microbiol. Rev. 2023, 47, fuad017.

(25) Arias-Orozco, P.; Yi, Y.; Ruijne, F.; Cebrián, R.; Kuipers, O. P. Investigating the specificity of the dehydration and cyclization reactions in engineered lanthipeptides by Synechococcal SyncM. ACS Synth. Biol. 2023, 12, 164–77.

(26) Le, T.; Zhang, D.; Martini, R. M.; Biswas, S.; van der Donk, W. A. Use of a head-to-tail peptide cyclase to prepare hybrid RiPPs. Chem. Commun. 2024, 60, 6508–11.

(27) Larsen, C. K.; Lindquist, P.; Rosenkilde, M.; Madsen, A. R.; Haselmann, K.; Glendorf, T.; Olesen, K.; Kodal, A. L. B.; Tørring, T. Using LanM enzymes to modify glucagon-like peptides 1 and 2 in *E. coli*. ChemBioChem 2024, 25, e202400201.

(28) Le, T.; Jeanne Dit Fouque, K.; Santos-Fernandez, M.; Navo, C. D.; Jiménez-Osés, G.; Sarksian, R.; Fernandez-Lima, F. A.; van der Donk, W. A. Substrate sequence controls regioselectivity of lanthionine formation by ProcM. J. Am. Chem. Soc. 2021, 143, 18733–43.

(29) Yu, Y.; Mukherjee, S.; van der Donk, W. A. Product formation by the promiscuous lanthipeptide synthetase ProcM is under kinetic control. J. Am. Chem. Soc. 2015, 137, 5140–8.

(30) Yang, X.; Lennard, K. R.; He, C.; Walker, M. C.; Ball, A. T.; Doigneaux, C.; Tavassoli, A.; van der Donk, W. A. A lanthipeptide library used to identify a protein-protein interaction inhibitor. Nat. Chem. Biol. 2018, 14, 375–80.

(31) Hegemann, J. D.; Bobeica, S. C.; Walker, M. C.; Bothwell, I. R.; van der Donk, W. A. Assessing the flexibility of the prochlorosin 2.8 scaffold for bioengineering applications. ACS Synth. Biol. 2019, 8, 1204–14.

(32) Vinogradov, A. A.; Chang, J. S.; Onaka, H.; Goto, Y.; Suga, H. Accurate models of substrate preferences of post-translational modification enzymes from a combination of mRNA display and deep learning. ACS Cent. Sci. 2022, 8, 814–24.

(33) Vinogradov, A. A.; Bashiri, G.; Suga, H. Illuminating substrate preferences of promiscuous F(420)H(2)-dependent dehydroamino acid reductases with 4-track mRNA display. J. Am. Chem. Soc. 2024, 146, 31124–36.

(34) Clark, J. D.; Mi, X.; Mitchell, D. A.; Shukla, D. Substrate prediction for RiPP biosynthetic enzymes via masked language modeling and transfer learning. Digit. Discov. 2025, 4, 343–54.

(35) Steude, E. G.; Dieckhaus, H.; Pelton, J. M.; Ren, J.; Kuhlman, B.; Bowers, A. A. Assessing Substrate Scope of the Cyclodehydratase LynD by mRNA Display-Enabled Machine Learning Models. Biochemistry 2025, 64, 2811–22.

(36) Do, T.; Link, A. J. Protein engineering in ribosomally synthesized and post-translationally modified peptides (RiPPs). Biochemistry 2023, 62, 201–9.

(37) Chang, J. S.; Vinogradov, A. A.; Zhang, Y.; Goto, Y.; Suga, H. Deep learning-driven library design for the de novo discovery of bioactive thiopeptides. ACS Cent. Sci. 2023, 9, 2150–60.

(38) Roberts, R. W.; Szostak, J. W. RNA-peptide fusions for the in vitro selection of peptides and proteins. Proc. Natl. Acad. Sci. USA 1997, 94, 12297–302.

(39) Huang, Y.; Wiedmann, M. M.; Suga, H. RNA display methods for the discovery of bioactive macrocycles. Chem. Rev. 2019, 119, 10360–91.

(40) Kamalinia, G.; Grindel, B. J.; Takahashi, T. T.; Millward, S. W.; Roberts, R. W. Directing evolution of novel ligands by mRNA display. Chem. Soc. Rev. 2021, 50, 9055–103.

(41) Vinogradov, A. A.; Nagai, E.; Chang, J. S.; Narumi, K.; Onaka, H.; Goto, Y.; Suga, H. Accurate broadcasting of substrate fitness for lactazole biosynthetic pathway from reactivity-profiling mRNA display. J. Am. Chem. Soc. 2020, 142, 20329–34.

(42) Fleming, S. R.; Himes, P. M.; Ghodge, S. V.; Goto, Y.; Suga, H.; Bowers, A. A. Exploring the post-translational enzymology of PaaA by mRNA display. J. Am. Chem. Soc. 2020, 142, 5024–8.

(43) Noda-Garcia, L.; Liebermeister, W.; Tawfik, D. S. Metabolite-enzyme coevolution: from single enzymes to metabolic pathways and networks. Annu. Rev. Biochem. 2018, 87, 187–216.

(44) Noda-Garcia, L.; Tawfik, D. S. Enzyme evolution in natural products biosynthesis: target- or diversity-oriented? Curr. Opin. Chem. Biol. 2020, 59, 147–54.

(45) Cubillos-Ruiz, A.; Berta-Thompson, J. W.; Becker, J. W.; van der Donk, W. A.; Chisholm, S. W. Evolutionary radiation of lanthipeptides in marine cyanobacteria. Proc. Natl. Acad. Sci. USA 2017, 114, E5424–E33.

(46) Le, T.; van der Donk, W. A. Mechanisms and evolution of diversity-generating RiPP biosynthesis. Trends Chem. 2021, 3, 266–78.

(47) Desormeaux, E. K.; Barksdale, G. J.; van der Donk, W. A. Kinetic analysis of cyclization by the substrate-tolerant lanthipeptide synthetase ProcM. ACS Catal. 2024, 14, 18310–21.

(48) Li, B.; Yu, J. P.; Brunzelle, J. S.; Moll, G. N.; van der Donk, W. A.; Nair, S. K. Structure and mechanism of the lantibiotic cyclase involved in nisin biosynthesis. Science 2006, 311, 1464–7.

(49) Tang, W.; Jiménez-Osés, G.; Houk, K. N.; van der Donk, W. A. Substrate control in stereoselective lanthionine biosynthesis. Nat. Chem. 2015, 7, 57–64.

(50) Medina-Franco, J. L.; Martínez-Mayorga, K.; Bender, A.; Scior, T. Scaffold diversity analysis of compound data sets using an entropy-based measure. QSAR Comb. Sci. 2009, 28, 1551–60.

(51) Mukherjee, S.; van der Donk, W. A. Mechanistic studies on the substrate-tolerant lanthipeptide synthetase ProcM. J. Am. Chem. Soc. 2014, 136, 10450–9.

(52) Pedregosa, F.; Varoquaux, G.; Gramfort, A.; Michel, V.; Thirion, B.; Grisel, O.; Blondel, M.; Prettenhofer, P.; Weiss, R.; Dubourg, V.; Vanderplas, J.; Passos, A.; Cournapeau, D.; Brucher, M.; Perrot, M.; Duchesnay, É. Scikit-learn: Machine Learning in Python. J. Mach. Learn. Res. 2011, 12, 2825–30.

(53) Lin, Z.; Akin, H.; Rao, R.; Hie, B.; Zhu, Z.; Lu, W.; Smetanin, N.; Verkuil, R.; Kabeli, O.; Shmueli, Y.; dos Santos Costa, A.; Fazel-Zarandi, M.; Sercu, T.; Candido, S.; Rives, A. Evolutionary-scale prediction of atomic-level protein structure with a language model. Science 2023, 379, 1123–30.

54. Lin, T.-Y.; Goyal, P.; Girshick, R.; He, K.; Dollár, P. Focal loss for dense object detection. arXiv 2017, arXiv:1708.02002

(55) Notin, P.; Kollasch, A. W.; Ritter, D.; van Niekerk, L.; Paul, S.; Spinner, H.; Rollins, N.; Shaw, A.; Weitzman, R.; Frazer, J.; Dias, M.; Franceschi, D.; Orenbuch, R.; Gal, Y.; Marks, D. S. ProteinGym: Large-Scale Benchmarks for Protein Design and Fitness Prediction. bioRxiv 2023, doi:10.1101/2023.12.07.570727.

56. Vieira, L. C.; Lin, S.; Wilke, C. O. Intrinsic dataset features drive mutational effect prediction by protein language models. bioRxiv 2026, doi:10.64898/2026.03.08.710389.

(57) Dufour, A.; Hindre, T.; Haras, D.; Le Pennec, J. P. The biology of lantibiotics from the lacticin 481 group is coming of age. FEMS Microbiol Rev 2007, 31, 134–67.

(58) Rink, R.; Kuipers, A.; de Boef, E.; Leenhouts, K. J.; Driessen, A. J.; Moll, G. N.; Kuipers, O. P. Lantibiotic structures as guidelines for the design of peptides that can be modified by lantibiotic enzymes. Biochemistry 2005, 44, 8873–82.

(59) Vinogradov, A. A.; Shimomura, M.; Goto, Y.; Ozaki, T.; Asamizu, S.; Sugai, Y.; Suga, H.; Onaka, H. Minimal lactazole scaffold for in vitro thiopeptide bioengineering. Nat. Commun. 2020, 11, 2272.

(60) Vinogradov, A. A.; Zhang, Y.; Hamada, K.; Chang, J. S.; Okada, C.; Nishimura, H.; Terasaka, N.; Goto, Y.; Ogata, K.; Sengoku, T.; Onaka, H.; Suga, H. De Novo discovery of thiopeptide pseudo-natural products acting as potent and selective TNIK kinase inhibitors. J. Am. Chem. Soc. 2022, 144, 20332–41.

(61) Thibodeaux, C. J.; Ha, T.; van der Donk, W. A. A price to pay for relaxed substrate specificity: a comparative kinetic analysis of the class II lanthipeptide synthetases ProcM and HalM2. J. Am. Chem. Soc. 2014, 136, 17513–29.

(62) King, A. M.; Anderson, D. A.; Glassey, E.; Segall-Shapiro, T. H.; Zhang, Z.; Niquille, D. L.; Embree, A. C.; Pratt, K.; Williams, T. L.; Gordon, D. B.; Voigt, C. A. Selection for constrained peptides that bind to a single target protein. Nat. Commun. 2021, 12, 6343.

