## Supporting Information for "Mapping the sequence preference of the generalist class II lanthipeptide synthetase ProcM by mRNA display"

### Materials and methods

Molecular biology experiments were performed using reagents from New England Biolabs, Thermo Fisher Scientific, CHEMIMPEX or Sigma-Aldrich. Anti-HA magnetic beads were from Pierce Thermo Fisher Scientific (#88837). Magnetic beads with immobilized streptavidin (SAv) were from Invitrogen (Dynabeads Streptavidin C1, # 65002). MALDI-TOF-MS data acquisition was performed using a Bruker UltrafleXtreme mass spectrometer (Bruker Daltonics) in reflector positive mode at the University of Illinois School of Chemical Sciences Mass Spectrometry Laboratory. A commercial mixture of 9:1 2,5-dihydroxybenzoic acid (DHB) and 2-hydroxy-5-methoxybenzoic acid (Super-DHB or SDHB) solution with a stock concentration of 25 mg/mL was routinely mixed with samples in a 1 to 1 ratio and air-dried before MALDI-TOF-MS analysis. MS1 and MS2 data acquisition was performed using an Agilent qTOF instrument equipped with a UPLC system. Ni-NTA resin for peptide and protein purifications was obtained from ThermoFisher.
Oligonucleotides for library assembly (HPLC purification grade for regular and randomized sequences, respectively) were purchased from IDT. All oligonucleotides were used as received. PCR amplifications were carried out in a C1000 Touch PCR thermal cycler. qPCR analysis was performed using the StepOnePlus™ Real-Time PCR System.
Expression and purification of ProcM and LahT150 were done as previously described.[1, 2]

**Library construction.** DNA libraries were assembled by two-step PCR from oligonucleotide primers (primer sequences are summarized in Supplementary Table 1) with Q5 high-fidelity 2x master Master mix. The assembly process was carried out separately for three libraries following the NEB Q5 protocol. Step 1: Overlapping primers were annealed and extended. Step 2: The crude extension product from step 1 was used as a template and added to a master mixture containing forward and reverse primers. The final DNA product was purified by gel extraction.

**Step 1: 50 μL PCR reaction**

A. Initial denature step 95 °C 2 min

B. Annealing 61 °C 1 min
C. Elongation 72 °C 1 min
10× between B and C.

**Step 2: 50 μL crude from step 1 was used as a template to setup a 500-μL PCR reaction**

A. Initial denature step 95 °C 2 min
B. Denature 95 °C 40 s
C. Annealing 61 °C 40 s
D. Elongation 72 °C 40 s

10× between B -D

Final elongation 72 °C for 2 min

**In vitro transcription**. MEGAscript™ T7 Transcription Kit and MEGAclear™ Transcription Clean-Up Kit were used following the manual provided by the manufacturer. To achieve maximum yield, the transcription was performed at 37 °C for 16 h. For the first round, a 60-μL transcription reaction was performed. Starting from the 2nd round, a 40-μL transcription reaction was performed.

| | |
|---|---|
| Nuclease-free Water | To a final 20 μL |
| NTPs | 2 μL each |
| template DNA | X μL (corresponding to 250-300 ng DNA) |
| T7 RNA Polymerase Mix | 2 μL |
| 10× buffer | 2 μL |

**Puromycin (Pu) ligation**. 1 μM mRNA library, 1.5 μM Pu linker (see Supplementary tables), 1 μM T4 RNA ligase (1 unit/μL) in 1× ligation buffer 100 μL (20% DMSO), 25 °C, 45 min. Ligated mRNA was purified by ethanol precipitation: The reaction was stopped after 45 min by adding a 1 × volume of quenching solution (0.6 M NaCl and 10 mM EDTA). Ligated mRNA was precipitated by adding 2× volume of 100% ethanol, and 0.02× volume of 5 mg/mL glycogen. This mixture was briefly vortexed and then centrifuged at 16,000 rpm for 15 min to pellet the puromycin-linked RNA. The supernatant was discarded, and the resultant pellet was washed with 70% ethanol and centrifuged again 16,000 × g for 5 min. The pellet was collected and air dried before resuspending in 1/10 the volume of water as the original T4 RNA ligation reaction (the final Pu-mRNA conc. 10 μM). This product was stored at -80 °C and directly used in PURExpress translation reaction. The efficiency of the reaction was determined by Urea-PAGE.

**In vitro translation.** The translation was carried out with PurExpress kit from NEB (E6850S). The first round was performed at a 20 μL scale, and starting from the second round, 10 μL of translation was performed.

**Assemble the reaction on ice in a new tube in the following order (for a 20 μL reaction):**

Solution A: 8 μL
Solution B: 6 μL
Murine RNase Inhibitor: 0.8 μL
Nuclease-free $H_2O$: 0.2 μL
Template (10 uM): 5 μL

After translation, incubation at 25 °C for 15 min was performed to facilitate mRNA-peptide complexes, EDTA (pH 8) was added to a final concentration of 17 mM, and the reaction was incubated at 37 °C for 30 min to remove the mRNA-peptide complexes from the ribosomes. A final concentration of 17 mM $MgCl_2$ was added to quench EDTA.

**Reverse transcription**. Complementary DNA was appended by a reverse transcription reaction containing all the translation products according to the protocol of the Superscript IV kit.
For a 20 μL reaction:

| | |
|---|---|
| Translation product | 10 μL |
| 100 μM Reverse primer | 0.8 μL |
| 10 mM dNTPs | 1 μL |
| Nuclease-free $H_2O$ | 2.2 μL |

The above reaction mixture was incubated at 65 °C for 5 min and cooled down on ice for 3 min. Then, 5x SIVV buffer in the Superscript IV kit (4 μL), 100 mM DTT (1 μL), and RTase (1 μL) were added. The resulting mixture was incubated at 50 °C for 10 min.

**ProcM assay.** The library was incubated with ProcM (final 3 μM), ATP (final 5 mM), $MgCl_2$ (final 5 mM), and TCEP (final 0.1 mM) in HEPES (100 mM, pH 7.5), at room temperature for 2 h. For round 5, the reaction time was shortened to 30 min in the selection, while the antiselection was performed for 2 h constantly. Accordingly, a no-enzyme control was carried out for both selection and antiselection.

**Biotin-IAA assay**. After ProcM assay, the library was incubated with TCEP (final 5 mM) on ice for 30 min before Biotin-(PEG)3-IAA (final 10 mM) was added in HEPES (100 mM, pH 7.5) at room temperature for 5 h.

**Library purification**. The library was diluted to 100 μL and added to pre-washed 35-μL anti-HA beads. The bead slurry was rotated at 4 °C for 1 h, after which the solution was loaded on a magnetic rack, the supernatant discarded and the beads were washed 3× with wash buffer (50 mM Tris HCl, 150 mM NaCl, pH 7.6, 0.05-0.1% Tween-20). Then 30 μL of HA peptide eluent (2.5 mg/mL in 50 mM Tris-HCl buffer, pH 7.6) was added to the above mixture and incubated at 37 °C for 20 min. The elution step was performed twice to achieve maximum yield.

**Acetone precipitation.** To remove excess free biotin probe completely, acetone precipitation of cDNA/mRNA-peptide was carried out. To one volume of the sample, three volumes of cold acetone were added, and the mixture was incubated at –20 °C for 30 min. The peptide–mRNA/cDNA conjugates were recovered by centrifugation (15300 *g* for 15 min at 4 °C). The supernatant was discarded. The pellets were washed with acetone and redissolved in wash buffer.

**Streptavidin pulldown.** The cyclized peptides (without biotinylation) were separated from the uncyclized peptide (biotinylated) by streptavidin pulldown. To one volume of the sample from above, one volume of blocking buffer (wash buffer with BAS additive) was added. Streptavidin C1 Dynabeads (~40 μL of 10 mg/mL bead slurry per library) were washed twice with the wash buffer, once with 1× blocking buffer, and added to the sample. Incubation at 4 °C for 60 min was carried out, after which the supernatant was separated as the unbound fraction, and the beads were washed twice with wash buffer containing 1 M urea and then once with wash buffer. Elution of the bound cDNA was carried out by heating the beads at 95 °C for 5 min suspended in nuclease-free water.

**qPCR Amplification.** The eluted unbound and bound cDNA were quantified by qPCR with SsoAdvanced Universal SYBR® Green Supermix following the protocol provided by the manufacturer.

20 μL qPCR reaction:
2× Supermix 10 μL
10 μM fwd 0.5 μL
10 μM rev 0.5 μL

Template sample 5 μL
MQ $H_2O$ 4 μL

qPCR cycle:
1. 95 °C, 2 min
2. 95 °C, 15 s
3. Anneal/extension, 60 °C 30 s (Cycles 40×, steps 2-3)

**NGS sequencing**. Tailing PCR was used to install Rd1 and Rd2 adapter sequences to library 5' and 3'-ends. The resulting DNA was sequenced by Azenta using Illumina paired-end 150 bp sequencing. cDNA was PCR-amplified using appropriate primers and 1-2 µL of the recovered fraction as the PCR template for the following round.
**Note**: For each library, total reads were at or above 10^7, and after Q30 filtering and constant-flanking-sequence filtering, the remaining reads were still around 10^7. Across the three initial libraries, the total number of unique sequences (in the naive library) was about 6.7 × 10^7. However, after the first round of selection, the unique sequence number dropped to the 10^5 level, with many repeated sequences being sequenced multiple times. We attribute this loss of diversity to a PCR recovery bottleneck.

**Peptide validation.** Linear double-stranded DNA templates encoding a T7 promoter sequence as well as the ORFs were translated following the PurExpress protocol. Each reaction was carried out at a 5-μL reaction scale. The resulting peptide was treated with ProcM under the same assay conditions described above. After ProcM assay, LahT digestion was performed to remove the leader peptide. The resulting mixture was preincubated with TCEP (final 5 mM) on ice for 10 min before NEM (final 10 mM) was added. The NEM assay was performed at 37 °C for 10 min. The reaction was desalted and analyzed by MALDI-ToF and qToF-LC-MS.

**High-resolution mass spectrometry**
The desalted LahT-digested peptides were injected onto an Agilent 1290 LC-MS QToF for ESI-HRMS analysis. LC separation was conducted at 50 ºC on a 5%-95% gradient of ACN/water (+0.1% formic acid) over 10 min at 1 mL/min flow rate on a Phenomenex C18-XB WIDEPORE column (part nr. 00D-4482-E0). Mass spectra were collected in positive mode at 10 spectra/s and 100 ms/spectrum.
Product distributions were quantified by integrating areas under peptide-derived peaks. Individual product areas, represented by A, were calculated by summing areas under extracted ion chromatogram (EIC) peaks, and cyclization ratios were calculated using this equation:

**Cyclization ratio** = $A_{cyclization}/A_{total}$

$A_{cyclization}$ is the area corresponding to the cyclization product in the EICs. $A_{total}$ is the total area corresponding to substrate-derived peaks, including cyclized peptide, non-dehydrated peptide, and dehydrated peptide.

**Definitions.** Normalized Shannon entropy ($H_{dataset}$) and Y* score were defined as previously reported.[3]
$f_{pep}$ is the peptide frequency, $C_{total}$ is the total number of reads in the dataset.

$$H_{dataset} = -\sum_{pep} \frac{f_{pep}^{*} \log_2 f_{pep}}{\log_2 c_{total}} \quad \text{equation 1}$$

$$Y^{*}_{aa,\,pos} = \frac{f^{selection}_{aa,\,pos}}{f^{antiselection}_{aa,\,pos}} \quad \text{equation 2}$$

**Enrichment** = cDNA recovery (with ProcM)/cDNA recovery (without ProcM)    (equation 3)

For selection, cDNA recovery = unbound/bound    (equation 4)

For antiselection, cDNA recovery = bound/unbound    (equation 5)

**Computational Methods for Binary Classification**

**S1. Dataset Preprocessing**

Following initial dataset aggregation, preprocessing was performed in two sequential filtering stages to improve dataset quality and consistency across peptide libraries (Ring 5, Ring 6, and Ring 7). The original combined dataset contained 488,007 peptide sequences, consisting of 112,795 bound (Class 0) and 375,212 unbound (Class 1) samples, corresponding to an overall class imbalance ratio of 3.3. In the first preprocessing stage, duplicate peptide sequences were removed, reducing the dataset to 416,949 total sequences, including 77,266 bound and 339,683 unbound samples. In the second stage, sequences violating library-specific sequence constraints were excluded, yielding the final dataset of 371,446 peptide sequences composed of 65,965 bound and 305,481 unbound samples, with a final overall class imbalance ratio of 4.6. Dataset statistics varied across libraries after preprocessing: Ring 5 decreased from 248,164 to 198,236 sequences, Ring 6 from 177,589 to 127,930 sequences, and Ring 7 from 62,254 to 45,280 sequences. The final processed dataset was subsequently used for all model training, validation, and evaluation experiments. Training and validation datasets were generated using stratified random sampling with a default 90:10 split.

**S2. Sequence Featurization**

Sequences were represented using either one-hot encoding over the 20 standard amino acids or precomputed embeddings from protein language models (ESM-2, PeptideESM2). Optional positional encodings, including sinusoidal and Fourier representations, were appended to improve sequence-aware learning but these did not substantially alter the performance.

**S4. Model Architectures**

The primary deep learning architecture was a multilayer perceptron (MLP) consisting of fully connected layers with batch normalization, ReLU activation, and dropout regularization. Classical machine learning models included k-nearest neighbors, AdaBoost, logistic regression, support vector machines, random forests, and gradient boosting classifiers.

**S5. Imbalance Handling**

Because the dataset was class imbalanced, Focal Loss[4] was used during neural network training to emphasize difficult samples and improve minority-class performance. The optimized focal loss parameters were $\alpha = 0.2016$ and $\gamma = 2.003$.

**S6. Training Procedure**

Models were trained using the Adam optimizer with weight decay regularization. Training was performed for up to 500 epochs with early stopping based on validation balanced accuracy. Performance metrics included loss, overall accuracy, per-class accuracy, and balanced accuracy. Random seeds were fixed to ensure reproducibility.

**S7. Hyperparameter Optimization**

Bayesian optimization was performed primarily using Optuna[5] with Tree-structured Parzen Estimator (TPE) sampling. Search parameters included learning rate, batch size, dropout, weight decay, focal loss parameters, and hidden-layer dimensions. The final optimized architecture used hidden dimensions [512, 256, 128, 64, 32] with a learning rate of

1.028 × $10^{-4}$.

**S8. Evaluation Metrics**

Model performance was evaluated using balanced accuracy, Matthews correlation coefficient (MCC), AUROC. Balanced accuracy was used as the primary model-selection metric because of class imbalance. Detailed performance metrics for all model architectures and embeddings type are shown in Table S1.

Table S1: Model Performance Comparison Across Different Embeddings

| Model | Embedding | AUROC | Bal. Acc. | MCC | Sens. | Spec. |
|---|---|---|---|---|---|---|
| GB | One-Hot | 0.7526±0.0030 | 0.6844±0.0023 | 0.4351±0.0025 | 0.9463±0.0012 | 0.4225±0.0056 |
| | ESM2 | 0.7431±0.0021 | 0.6680±0.0017 | 0.4169±0.0020 | 0.9535±0.0009 | 0.3825±0.0042 |
| | PeptideESM2 | 0.7442±0.0015 | 0.6680±0.0022 | 0.4171±0.0038 | **0.9537±0.0007** | 0.3823±0.0048 |
| MLP | One-Hot | **0.7890±0.0041** | **0.7343±0.0035** | **0.4703±0.0056** | 0.9072±0.0060 | **0.5614±0.0115** |
| | ESM2 | 0.7548±0.0032 | 0.7080±0.0028 | 0.3949±0.0099 | 0.8697±0.0120 | 0.5464±0.0126 |
| | PeptideESM2 | 0.7553±0.0051 | 0.7061±0.0054 | 0.3942±0.0083 | 0.8735±0.0086 | 0.5387±0.0165 |
| RF | One-Hot | 0.7607±0.0030 | 0.6987±0.0022 | 0.4271±0.0031 | 0.9221±0.0022 | 0.4753±0.0060 |
| | ESM2 | 0.7509±0.0022 | 0.6907±0.0016 | 0.4313±0.0019 | 0.9362±0.0011 | 0.4452±0.0041 |
| | PeptideESM2 | 0.7515±0.0030 | 0.6915±0.0012 | 0.4339±0.0011 | 0.9371±0.0008 | 0.4459±0.0032 |
| SVM | One-Hot | 0.7480±0.0021 | 0.6914±0.0014 | 0.3591±0.0020 | 0.8578±0.0006 | 0.5251±0.0035 |
| | ESM2 | 0.7479±0.0017 | 0.6927±0.0019 | 0.3560±0.0029 | 0.8492±0.0005 | 0.5363±0.0041 |
| | PeptideESM2 | 0.7487±0.0028 | 0.6955±0.0013 | 0.3612±0.0027 | 0.8502±0.0009 | 0.5408±0.0018 |

**S9. *In Silico* Sequence Generation for Experimental Validation**

To generate peptide candidates for prospective experimental validation, *in silico* peptide libraries were constructed for the Ring 5, Ring 6, and Ring 7 scaffolds using constrained stochastic sequence generation. Each library followed a predefined sequence template containing fixed framework residues and variable positions. Conserved residues required for scaffold integrity and library identity were retained, while designated variable positions were randomized to explore sequence space within experimentally allowable constraints.

Randomization was restricted to a subset of permitted amino acids to maintain compatibility with library-specific design requirements. Amino acids C, A, F, T, W, Y, and P were excluded from variable positions because these residues were disallowed by sequence-design constraints. Sequence generation therefore sampled only from the remaining allowed amino acid set at randomized positions.

To maximize sequence novelty and minimize redundancy, generated candidates were filtered using a minimum Hamming-distance criterion. Each candidate sequence was required to differ by at least two residue substitutions (minimum Hamming distance = 2) from all sequences in the original experimental datasets as well as from all previously accepted generated sequences within the same library. Candidate peptides failing this criterion were rejected and regenerated. This filtering step prevented near-duplicate sequences and promoted broader exploration of sequence space while preserving scaffold-specific structural constraints.

A total of 100,000 candidate sequences were generated for each library following diversity filtering. Generated sequences were subsequently evaluated using the trained machine learning classifier to obtain predicted binding probabilities for each peptide.

To construct an experimentally balanced validation set spanning the full prediction-confidence range, generated peptides were uniformly subsampled across the predicted probability distribution. The probability range was divided into five equal-width bins, and approximately equal numbers of sequences were sampled from each interval. This strategy ensured representation of high-confidence, intermediate-confidence, and low-confidence predictions during downstream experimental testing, enabling systematic evaluation of model calibration and predictive performance across the full spectrum of predicted activities.

**Table S2**. Library diversity of Ring 5, Ring 6, and Ring 7 before ligation with puromycin.

| Library | Core peptide | Theoretical diversity | Actual diversity | NGS reads |
|---|---|---|---|---|
| **Ring 5** | AXXSXXXCXXPHDYWEGEA | 6.2*10^7 | 2.2*10^7 | 5.8*10^7 |
| **Ring 6** | AADHNXXCXXXXSXXEGEA | 8.1*10^8 | 2.6*10^7 | 4.7*10^7 |
| **Ring 7** | AACXXXXXSXXPAYWEGEA | 6.2*10^7 | 1.7*10^7 | 4.9*10^7 |

**Figure S1**. (A) Sequences of 29 precursor peptides encoded by the genome of *Prochlorococcus* MIT 9313.[6] These peptides contain highly conserved leader peptides and diverse core peptides. (B) Structures of several representative prochlorosins after ProcM maturation showing the diverse ring patterns. For Lan/MeLan, atoms originating from Ser/Thr are shown in blue, and from Cys in red, dehydrated residues and residues after hydrolysis of N-terminal dehydrobutyrine are in purple. Pcn = prochlorosin.

## A

```
                          Leader peptide                                                         Core peptide
1.1  MSEEQLKAFIAKVQADTSLQEQLKA--EGADVVAIAKAAGFSITTEDLE-KEHRQ-------TLSDDDLEGVAGG FFCVQGTANRFTINVC---------------
1.2  MSEEQLKAFIAKVQADPSLQEQLKA--EGADVVSIAKAAGFSITTEDL--NSHIT----TKLNLSEEELEGVAGA MDCVSSTAQQTECRPGGPRASYCWDDLR----
1.3  MSEEQLKGFLSKVQSDASLQEQLKV--EGADVVAIAKAAGFSITTEDL--NSHRQ-------NLSEDELEGVAGG GLCTLTSNLAAVCCGGCRRATSE---------
1.4  MSEEQLKAFLTKVQADTSLQEQLKA--EGADPVAIAKAAGFAITTEDL--NSHRQ-------NLSDDELEGVAGG GSSYRNGKCTFGPACPS--------------
1.5  MSEEQLKAFIAKVQADTSLQEQLKV--EGADVVAIAKAAGFSITTEDL--NTHRQ-------NLSDDELEGLHGA GPGCTGGWWAFTDTAGGGSCEG----------
1.6  MSEEQLKAFIAKVQADTSLQEQLKV--EGADVVAIAKAAGFSITTDDFERNTHRQ-------TLSDDELEGVAGG KSTNGCGCKPGHTLSSFLCTLECWL-------
1.7  MSEEQLKAFIAKVQADTSLQEQLKV--EGADVVAIAKASGFAITTEDL--KAHQAN---SQKNLSDAELEGVAGG TIGGTIVSITCETCDLLVGKMC----------
2.1  MSEEQLKAFIAKVQADSSLQEQLKA--EGADVVAIAKAAGFSITTEDWDQRPVR--------TLSDEELEGAAGG CCITGESPGSAPTNDYKCTKGRGPGGCY----
2.2  MSEEQLKAFIAKVQADPSLQEQLKA--EGADTVAIAKAAGFSITTEDL--KEHRQ-------TLSDDELESVAGG GNDTVITKEYSCYVTSDKGCC-----------
2.3  MSEEQLKAFLEKVKADTSLKEKLKAAKSPEDVVGIAKEHGHEFTADKI------S-------QLSEEELEGVAGG MQAGSCNWICFVNGVYINDGRMANKAI-----
2.4  MSEEQLKAFLAKVQADASLQEQLKA--EGADVVAIAKAAGFSITTEDL--NSHRQ------IEMTDDELEGVAGG GCGLGARRETAQCWLSH--------------
2.5  MSEEQLKAFIAKVQGDTSLQEQLKA--EGADVVAIAKAAGFAITEAEV--KAYQT------RNLSDEELDEVAGG APCRPFTDPIYCWRKGEQTIIGRGRSCLYPE-
2.6  MSEEQLKAFIAKVQADTSLQEQLKA--EGADVVAIAKAAGFAISTEDL--NNHRQ-------NLSDDELEGVAGG GICVYVNCVLSIRETPSVI-------------
2.7  MSEEQLKAFIAKVQADTSLQEQLKA--EGADVVAIAKAAGFSIATEDL--KTHRQ-------TLSDDDLEGVAGG AGCYPICDWTSPTRS-----------------
2.8  MSEEQLKAFLTKVQADTSLQEQLKI--EGADVVAIAKAAGFSITTEDL--NSHRQ-------NLSDDELEGVAGG AACHNHAPSMPPSYWEGEC-------------
2.9  MSEEQLKAFIAKVQADPSLQEQLKA--EGADVVAIAKAAGFTITTEDL--KTARQ-------TLSDDDLEGVAGG YEDGDYTKSISIVVACCRF-------------
2.10 MSEEQLKAFIAKVQADSSLQEQLKA--EGADPVSIAKAAGFTITTEDL--NSHRQ-------NLSDEELEGAAGG AGGTIPSLMTGCGWLTGLCVR-----------
2.11 MSEEQLKAFIAKVQADTSLQEQLKA--EGADVVAIAKAAGFAITKEDL--NSHRQ-------TLSEDELESVAGG GRIDTCPAGGGTSEQTGTCC------------
3.1  MSEEQLKAFIAKVQADASLQEQLRT--EGADVVAIAKAAGFSITTEDL--NSHRQ-------NLSDDELEGVAGG GGKMTVRGRDMSCGCQDYWEDDY---------
3.2  MSEEQLKAFIAKVQADASLQEQLRT--EGADVVAIAKAAGFSITTEDL--NSHRQ-------NLSDDELEGVAGG GGGCDGIRITDKQTVADNTIVPCSCFHQ----
3.3  MSEEQLKAFIAKVQGDSSLQEQLKA--EGADVVAIAKAAGFTIKQQDL--NAAAS-------ELSDEELEAASGG GDTGIQAVLHTAGCYGGTKMCRA---------
3.4  MSEEQLKAFIAKVQGDTSLQEQLKA--EGADVVAIAKAAGFSITTEDL--NTHRQ-------TLSDRELEGVAGG TTAFTGVDTESIAFCCS---------------
3.5  MSEEQLKAFLEKVKADTSLQEKLKAAADSDAVLVIAKDAGFSISADDL--KNAQS-------EISEEELESVAGG AGVTEATIDAGGGCTFNPCCR-----------
4.1  MSEEQLKALIAKVQADTSLQEQLKA--EGADVVAIAKAAGFSITTEDL--KEHRQTLSVGRQTLSESELEGLAGG GGGARTKTANVPSDLPVRAPAMSTFAENQT--
4.2  MSEEQLKAFIAKVQADTSLQEQLKA--EGADPVAIAKAAGFSITTEDL--KEHRQ-------ALSDDDLEGVAGG TIVTVTGALISIAAEC----------------
4.3  MSEEQLKAFIAKVQADTSLQEQLKA--EGADVVAIAKAAGFTITTEDL--NSHRQ-------NLTDDELEGVAGG TASGGCDTSMFCY-------------------
4.4  MSEEQLKAFLEKVKGDTSLQEKLQAAADSDAVLAIAKEAGFMISADEL--KKAQS-------EISEEELESAAGG RLKSGCHCGTVIRSYSKYC-------------
s.1  MSEEQLKAFLEKVKADTSLQEKLKAAADSDAVLVIAKDAGFSISADDL--KNAQS-------EISEEELESVAGG AQSAGGCGICECDNRQSTSCHYPSHG------
s.2  MSEEQLKAFLEKVKADTSLQEKLKAAAGSDAVLAIAKAAGLMISADDL--TKAQS-------EISDAELEDAAGG GAQGPACCAAMESSDTRCGWVSWVLSEVVPPQ
```

## B

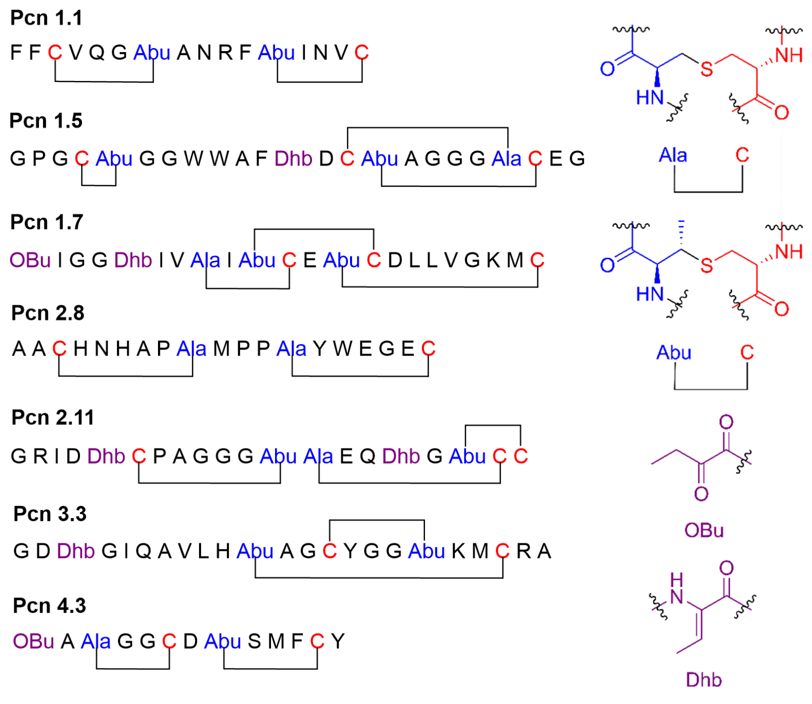

**Figure S2**. Normalized Shannon entropy was calculated using equation 1, which indicated the peptide level convergence. NGS sequencing was not performed for round 1 of Ring 5 and Ring 6.

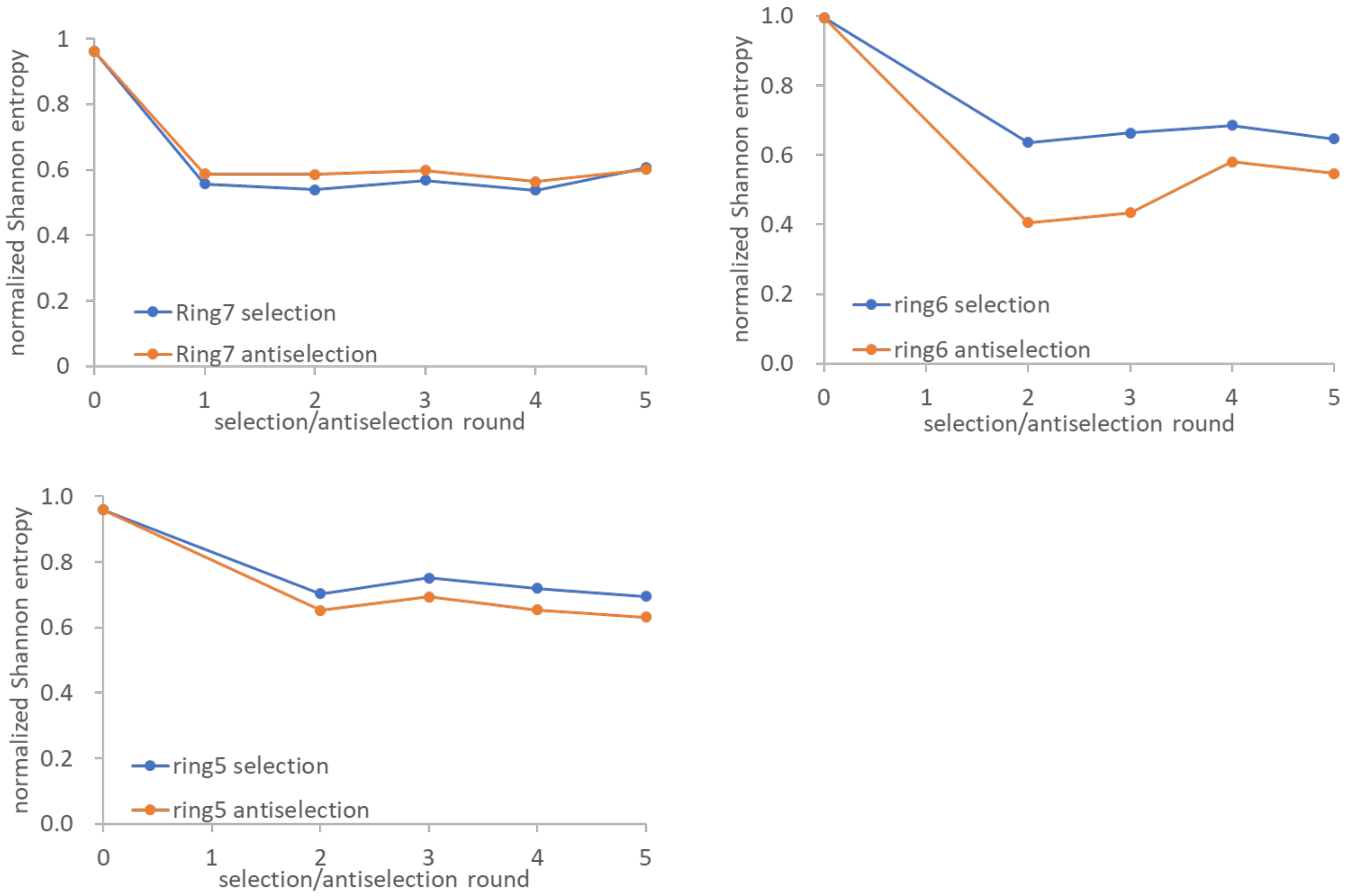

**Figure S3**. Summary of the selection experiments. The cDNA recovery and enrichment were calculated by qPCR based on equations 3-5.

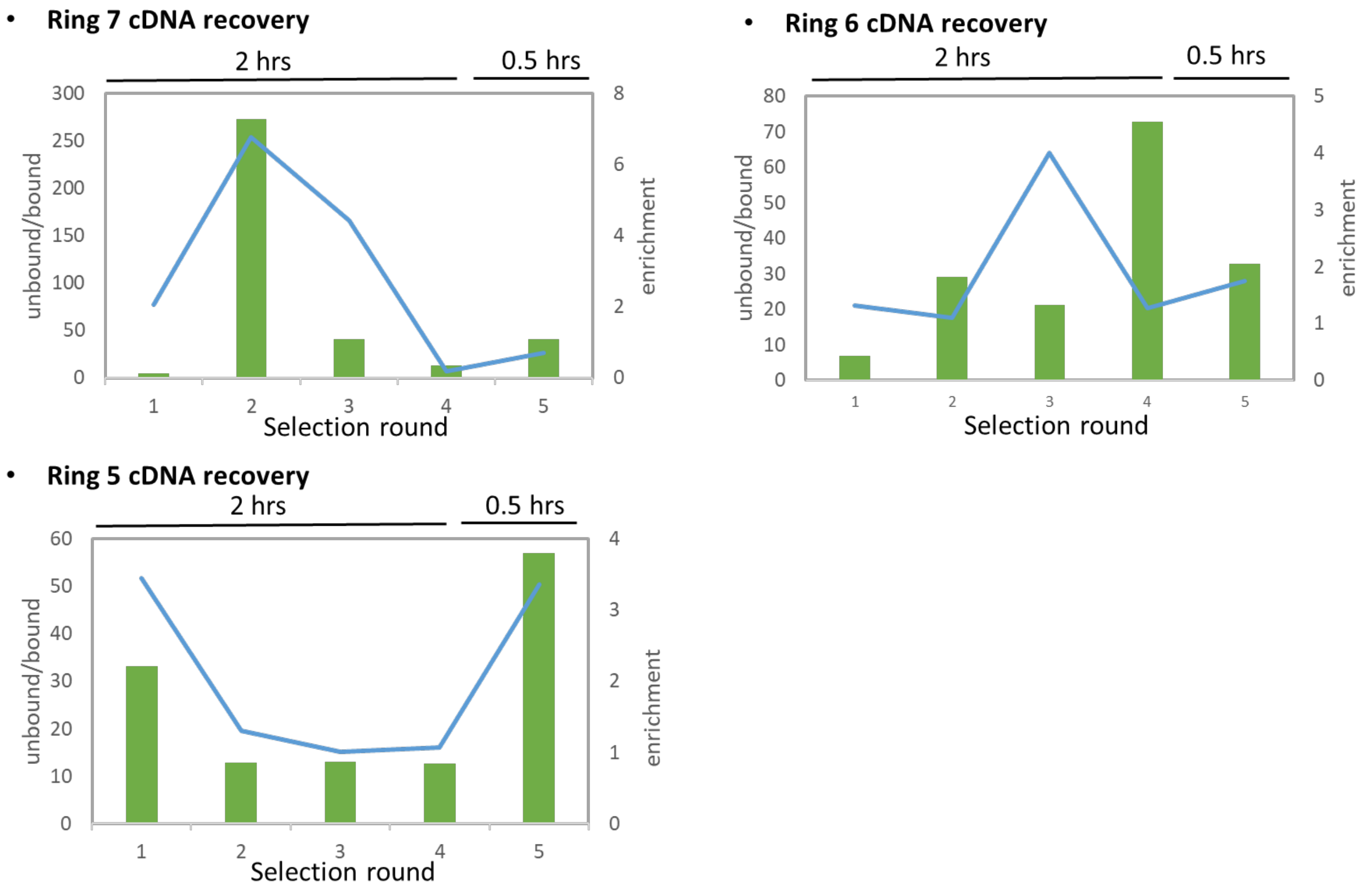

**Figure S4**. Summary of the antiselection experiments. The cDNA recovery and enrichment were calculated by qPCR based on equations 3-5.

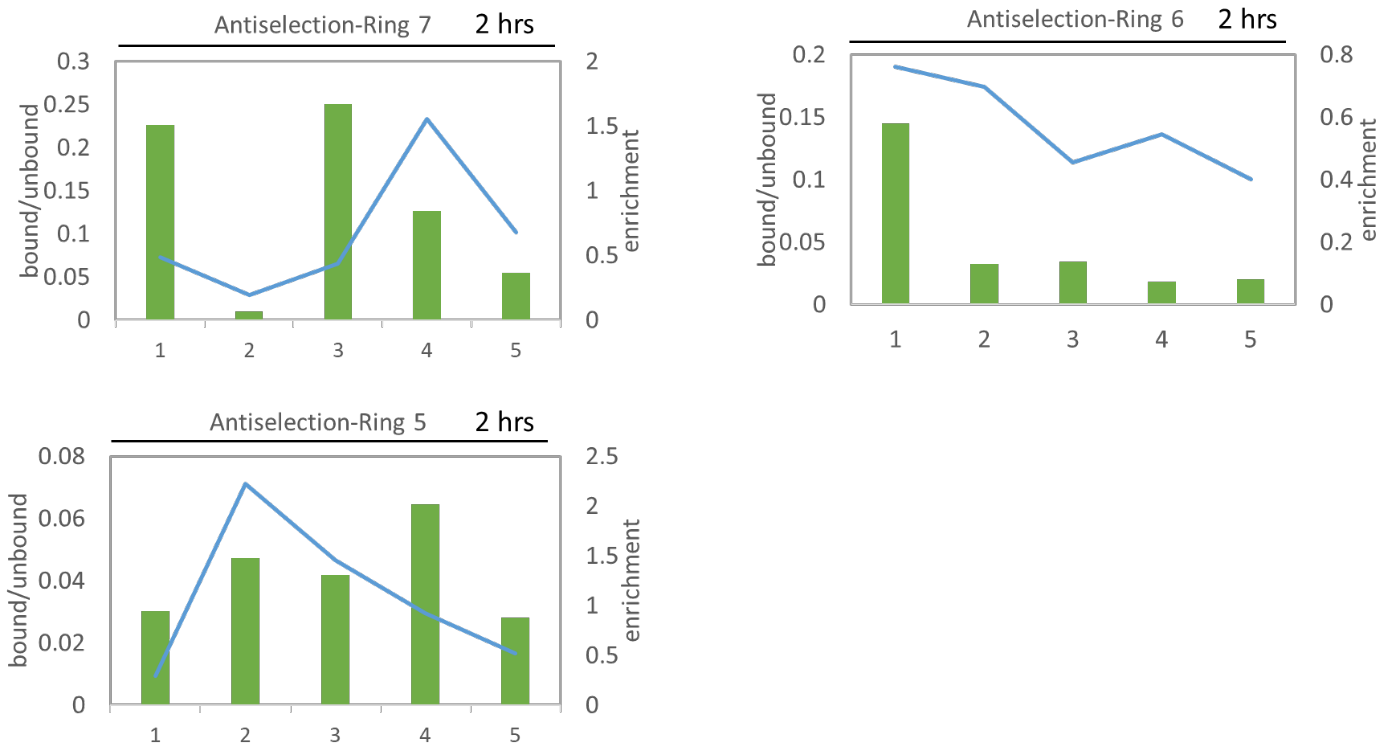

**Figure S5**. HR-LC-MS or MALDI-ToF spectra of validation peptides **v1-v60** after ProcM assay, LahT digestion, and NEM assay. Peptides **v1-v60** were prepared with PurExpress. Except for **v24** and **v33**, all peptides were analyzed by LC-MS. **v24** and **v33** peptides were not visible on LC-MS and were therefore analyzed by MALDI-ToF MS. The exact mass, predicted probability, and cyclization ratio are summarized in the Supplementary file of validation peptides.

**v1**

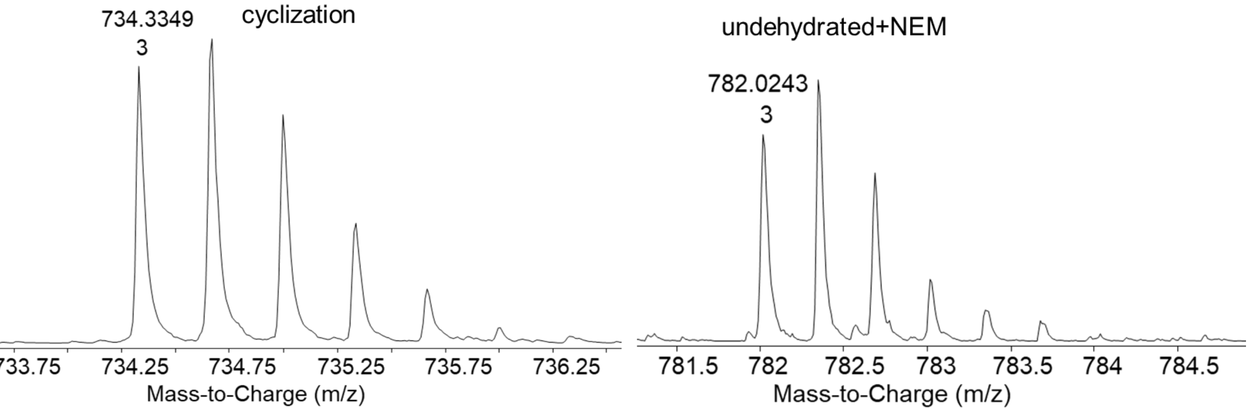


**v2**

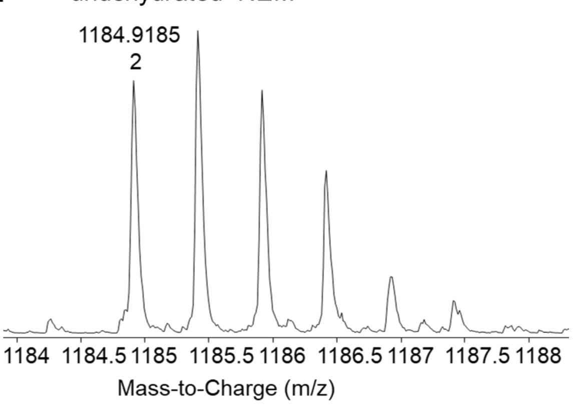

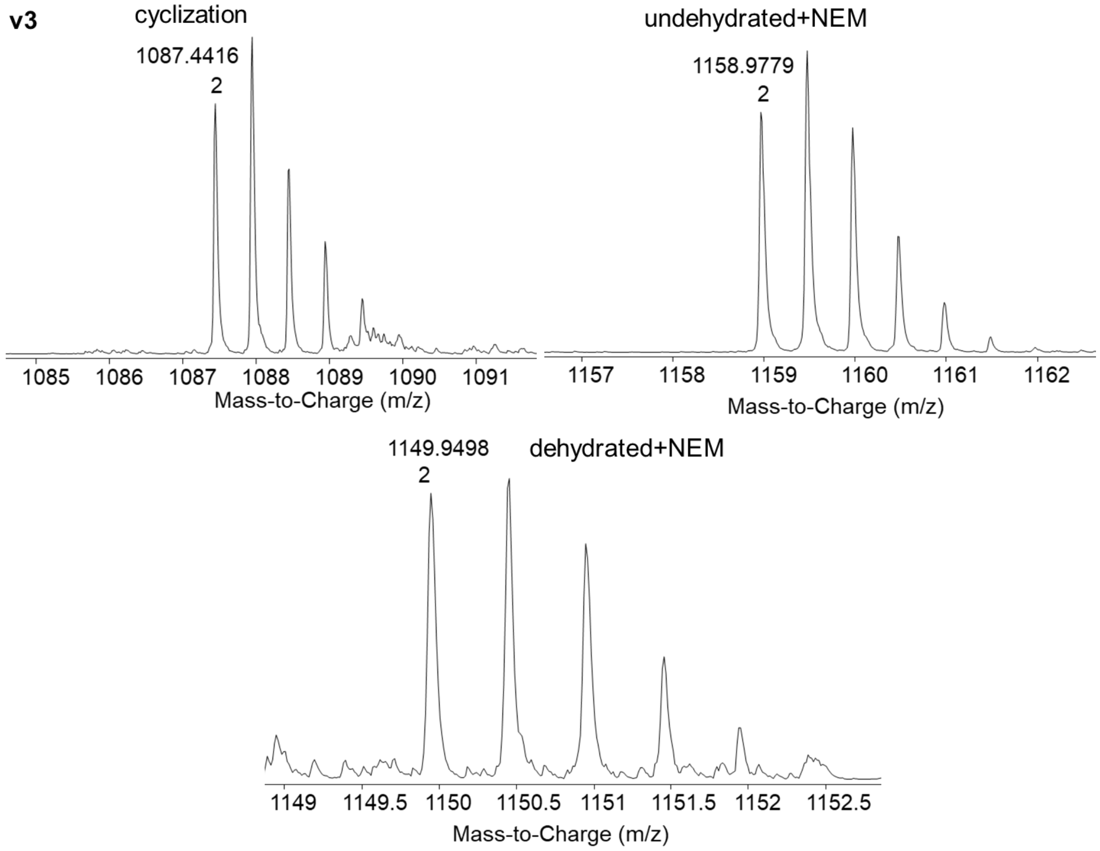
v3
cyclization
1087.4416
2
undehydrated+NEM
1158.9779
2
Mass-to-Charge (m/z)
Mass-to-Charge (m/z)
1149.9498
2
dehydrated+NEM
Mass-to-Charge (m/z)

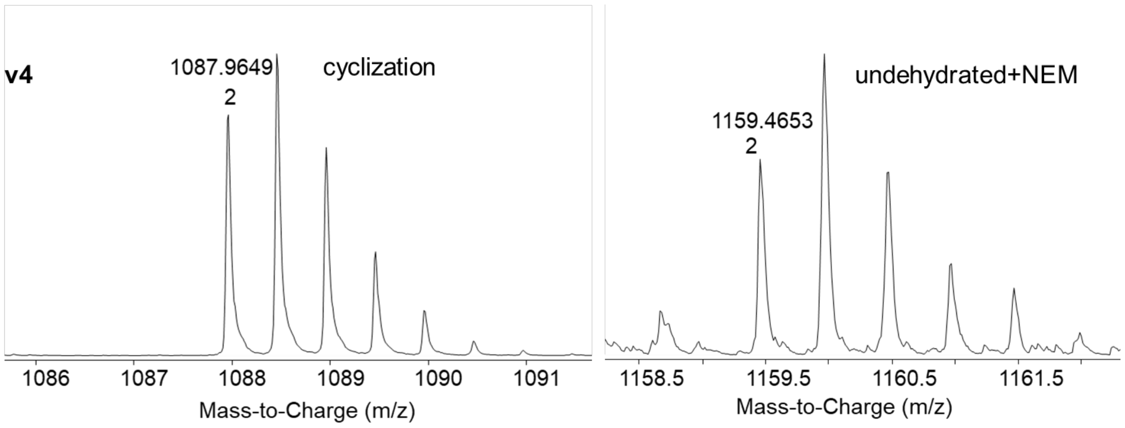
v4
1087.9649
2
cyclization
Mass-to-Charge (m/z)
undehydrated+NEM
1159.4653
2
Mass-to-Charge (m/z)

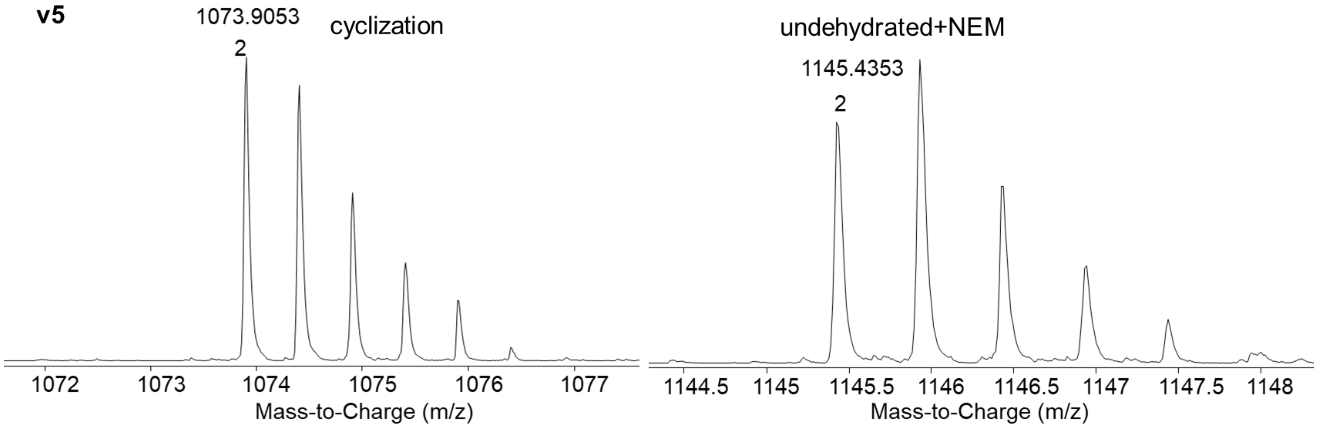
v5
1073.9053
2
cyclization
Mass-to-Charge (m/z)
undehydrated+NEM
1145.4353
2
Mass-to-Charge (m/z)

**v6**

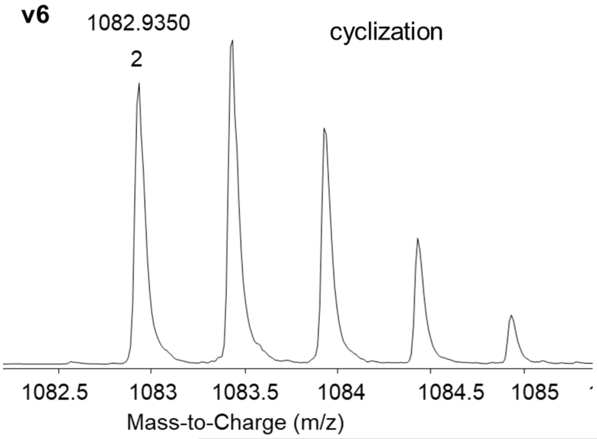
1082.9350
2
cyclization
1082.5
1083
1083.5
1084
1084.5
1085
Mass-to-Charge (m/z)

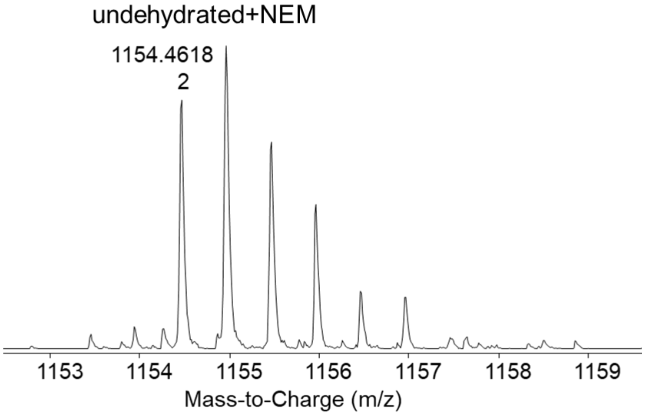
undehydrated+NEM
1154.4618
2
1153
1154
1155
1156
1157
1158
1159
Mass-to-Charge (m/z)

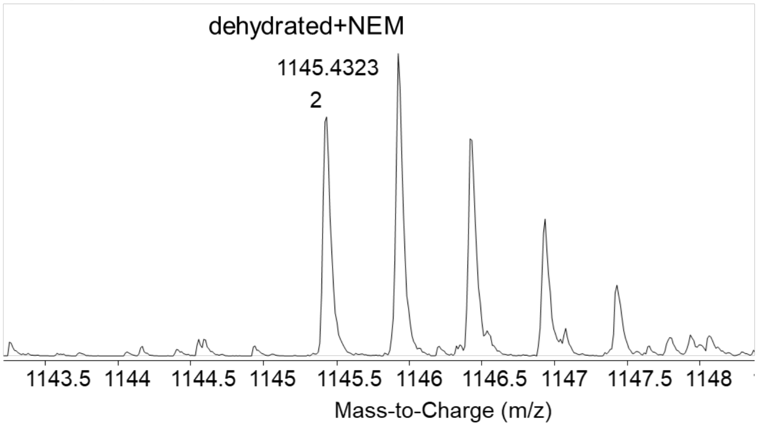
dehydrated+NEM
1145.4323
2
1143.5
1144
1144.5
1145
1145.5
1146
1146.5
1147
1147.5
1148
Mass-to-Charge (m/z)

**v7**

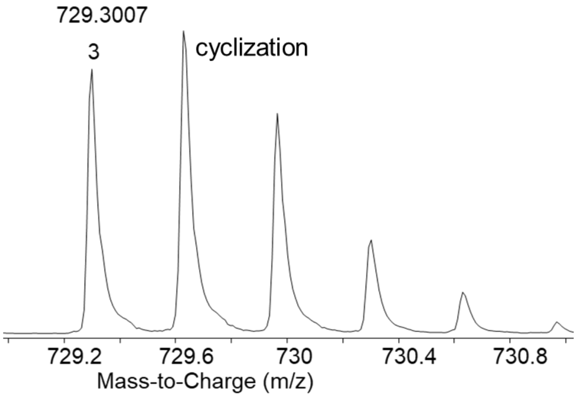
729.3007
3
cyclization
729.2
729.6
730
730.4
730.8
Mass-to-Charge (m/z)

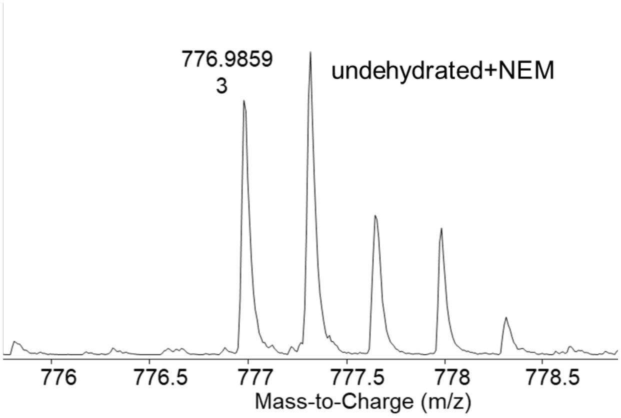
776.9859
3
undehydrated+NEM
776
776.5
777
777.5
778
778.5
Mass-to-Charge (m/z)

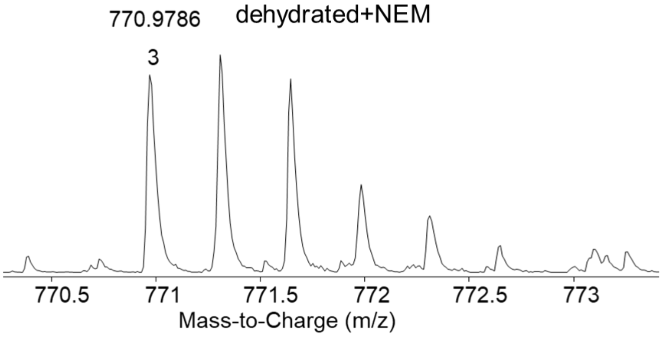
770.9786
3
dehydrated+NEM
770.5
771
771.5
772
772.5
773
Mass-to-Charge (m/z)

**v8**

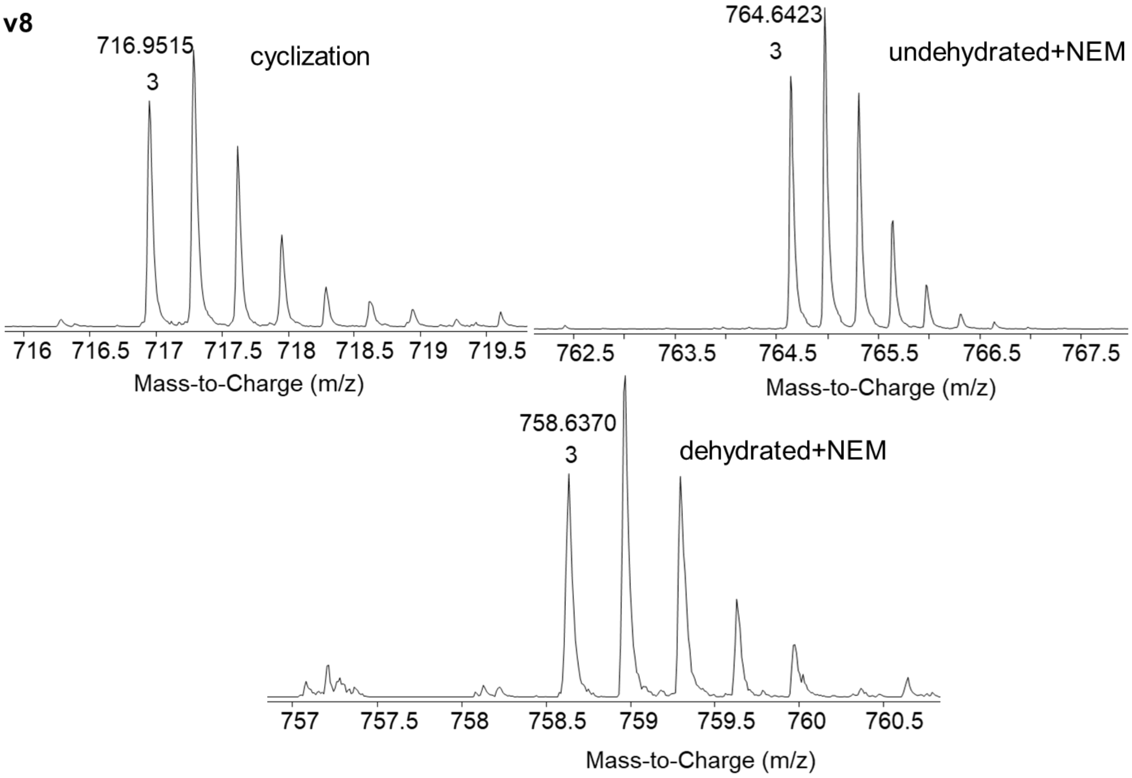
716.9515
3
cyclization
Mass-to-Charge (m/z)
764.6423
3
undehydrated+NEM
Mass-to-Charge (m/z)
758.6370
3
dehydrated+NEM
Mass-to-Charge (m/z)

**v9**

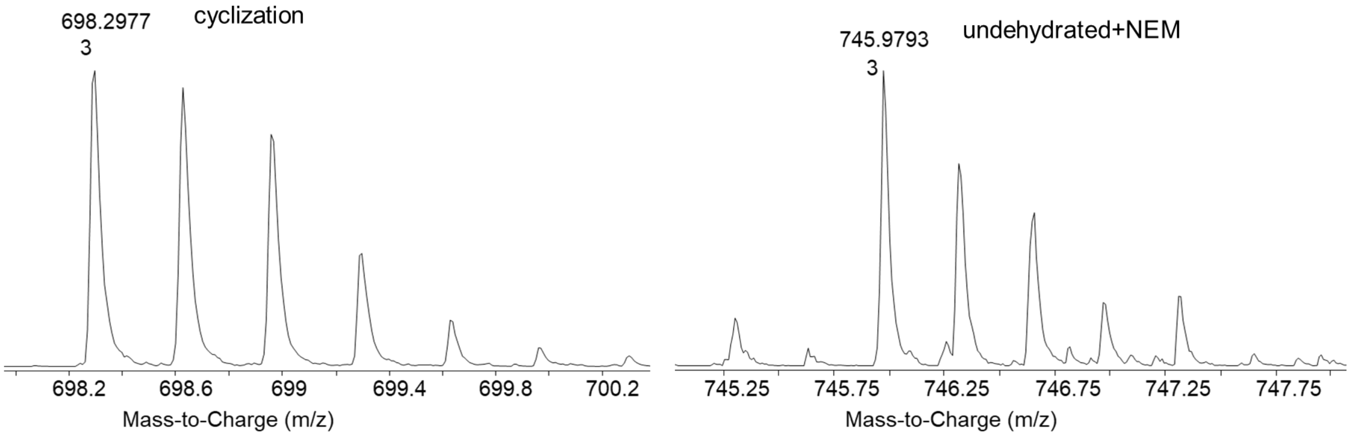
698.2977
3
cyclization
Mass-to-Charge (m/z)
745.9793
3
undehydrated+NEM
Mass-to-Charge (m/z)

**v10**

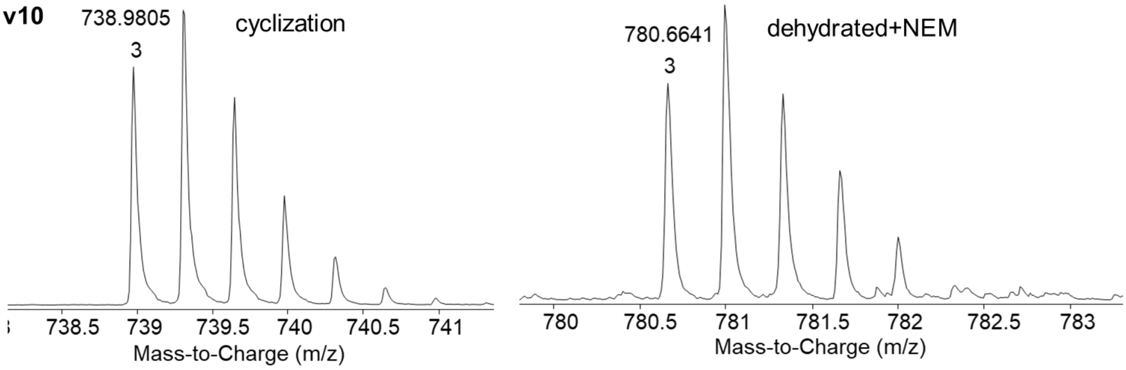
738.9805
3
cyclization
Mass-to-Charge (m/z)
780.6641
3
dehydrated+NEM
Mass-to-Charge (m/z)

**v11**

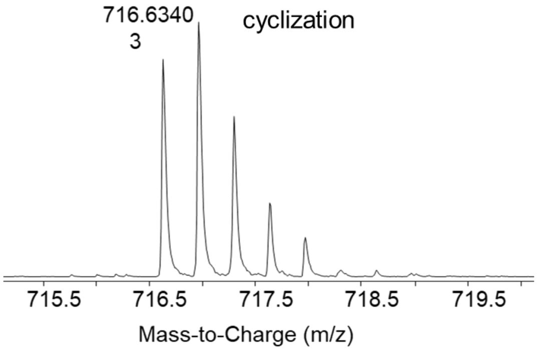
716.6340
3
cyclization
715.5 716.5 717.5 718.5 719.5
Mass-to-Charge (m/z)

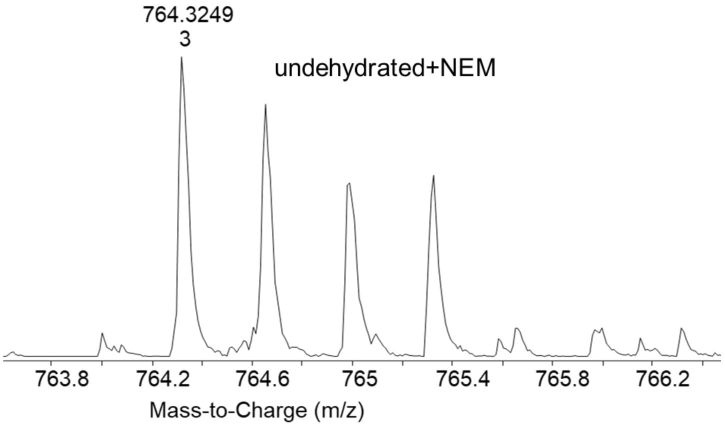
764.3249
3
undehydrated+NEM
763.8 764.2 764.6 765 765.4 765.8 766.2
Mass-to-Charge (m/z)

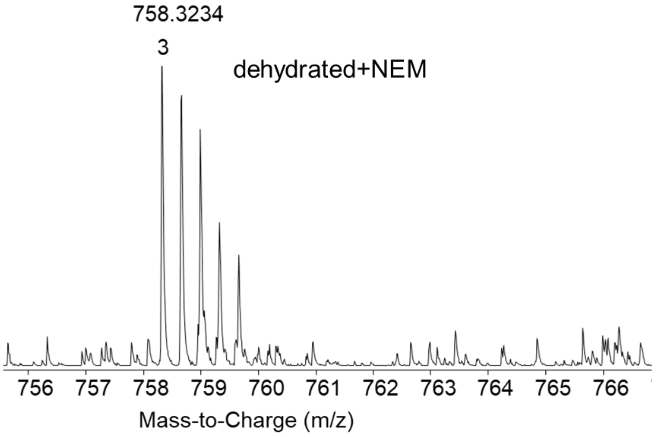
758.3234
3
dehydrated+NEM
756 757 758 759 760 761 762 763 764 765 766
Mass-to-Charge (m/z)

**v12**

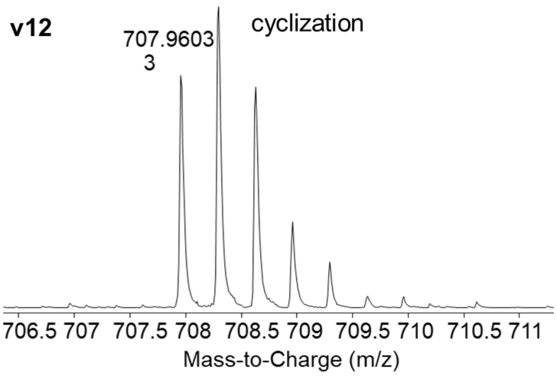
707.9603
3
cyclization
706.5 707 707.5 708 708.5 709 709.5 710 710.5 711
Mass-to-Charge (m/z)

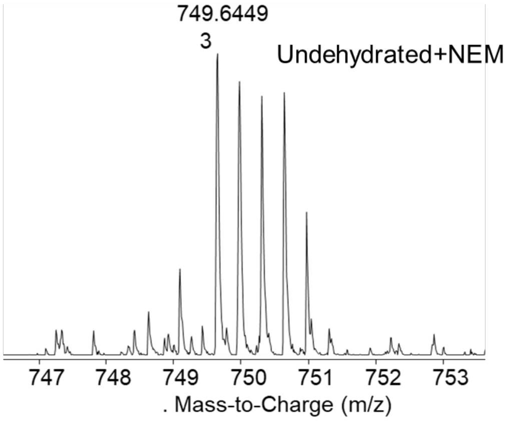
749.6449
3
Undehydrated+NEM
747 748 749 750 751 752 753
. Mass-to-Charge (m/z)

Mass-to-Charge (m/z)

**v13**

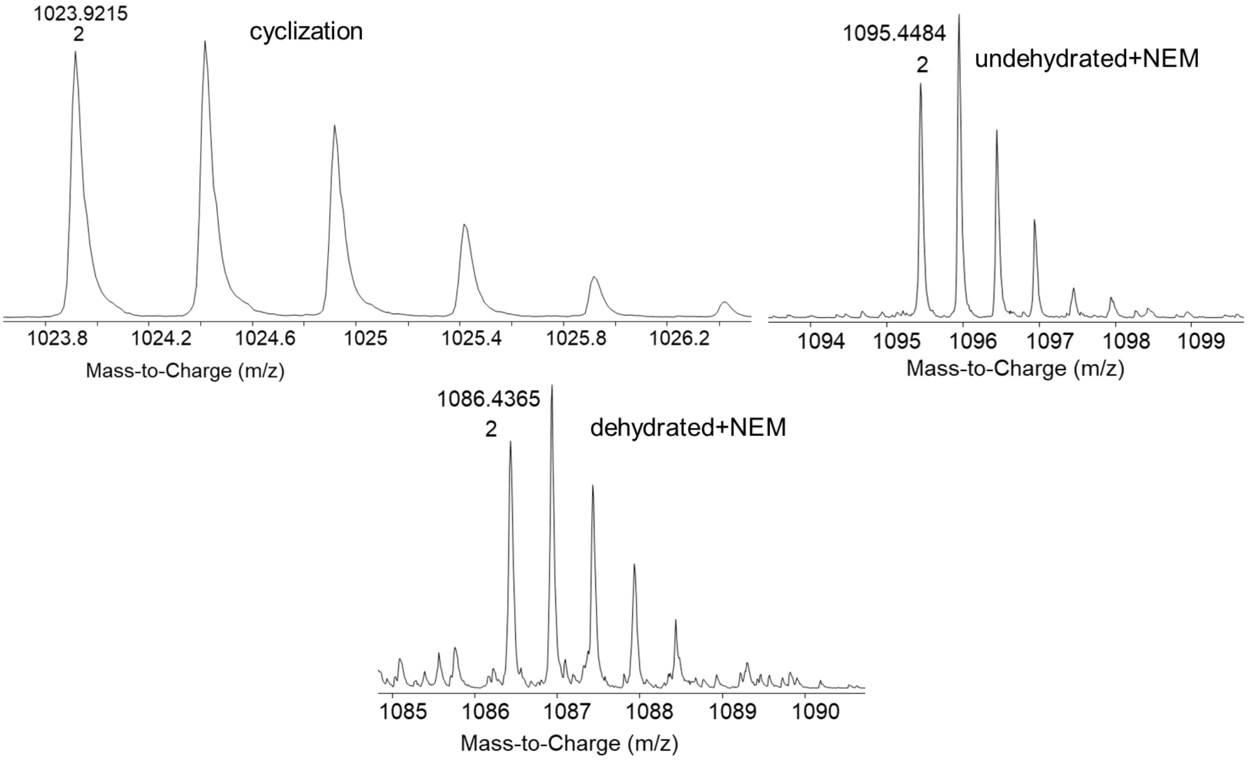


**v14**

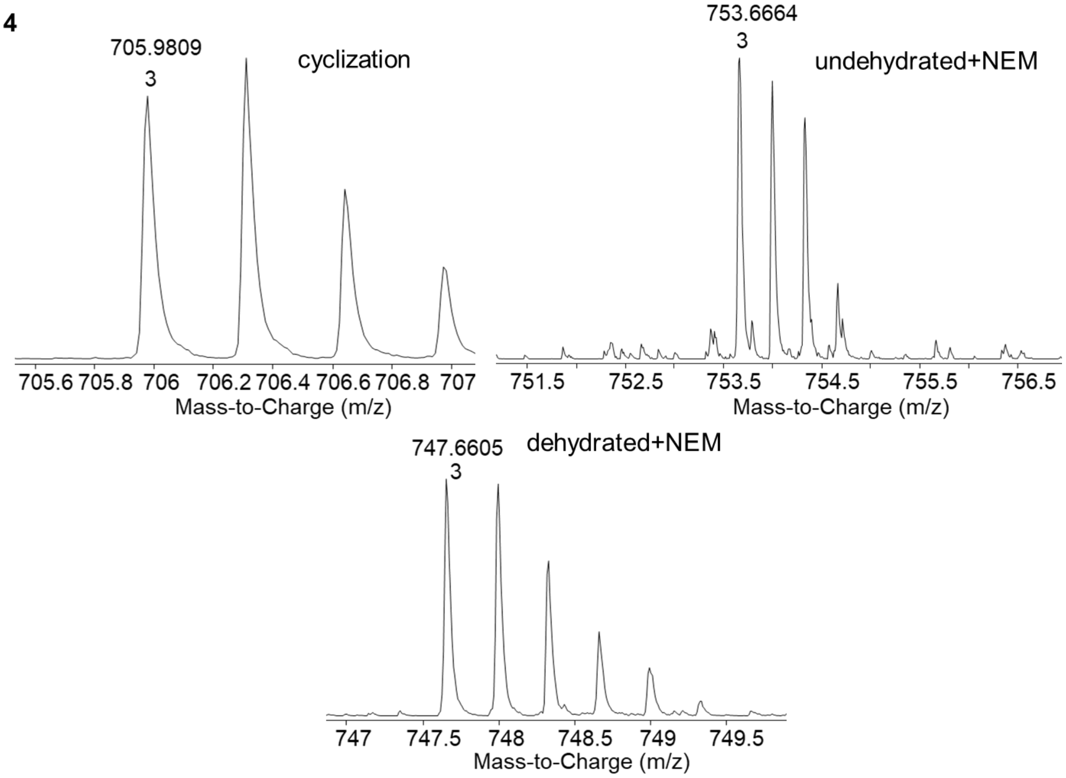

**v15**

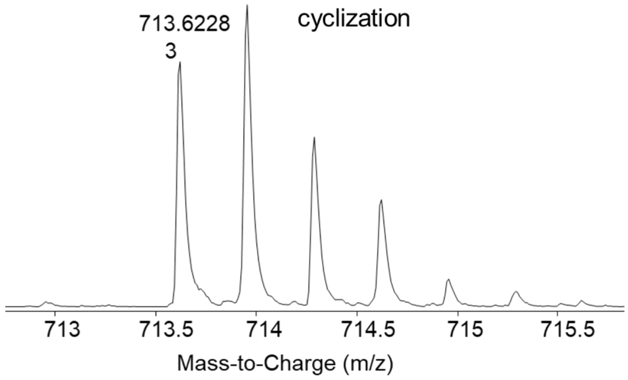
713.6228
3
cyclization
713
713.5
714
714.5
715
715.5
Mass-to-Charge (m/z)

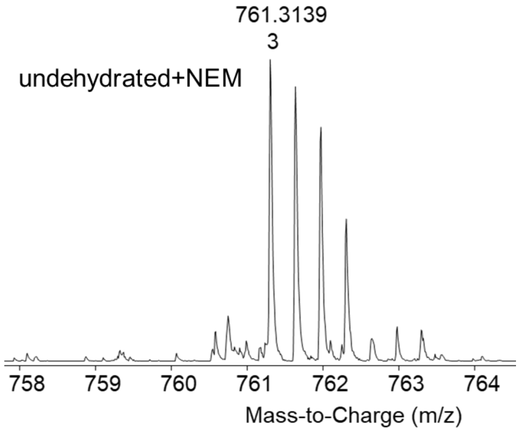
761.3139
3
undehydrated+NEM
758
759
760
761
762
763
764
Mass-to-Charge (m/z)

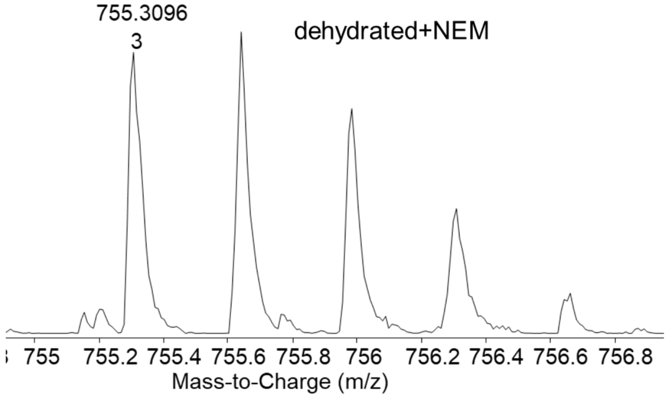
755.3096
3
dehydrated+NEM
755
755.2
755.4
755.6
755.8
756
756.2
756.4
756.6
756.8
Mass-to-Charge (m/z)

**v16**

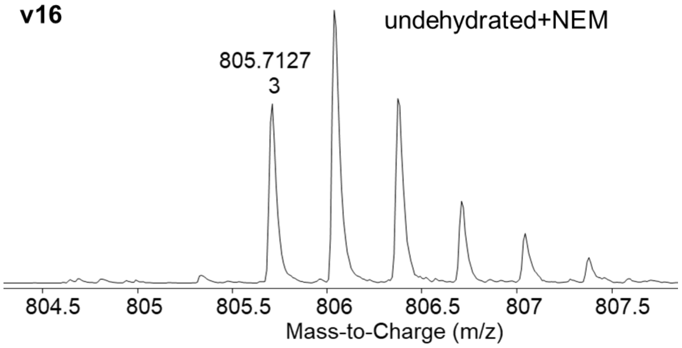
805.7127
3
undehydrated+NEM
804.5
805
805.5
806
806.5
807
807.5
Mass-to-Charge (m/z)

**v17**

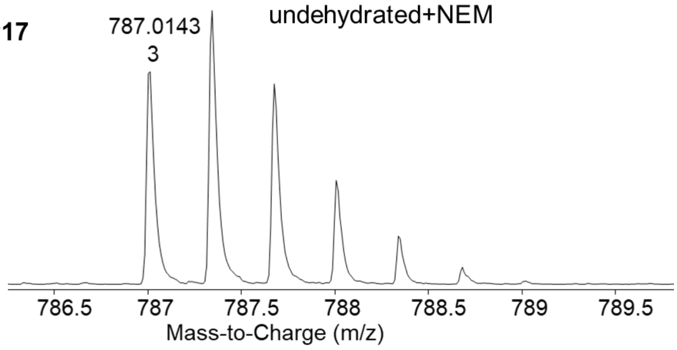
787.0143
3
undehydrated+NEM
786.5
787
787.5
788
788.5
789
789.5
Mass-to-Charge (m/z)

**v18**

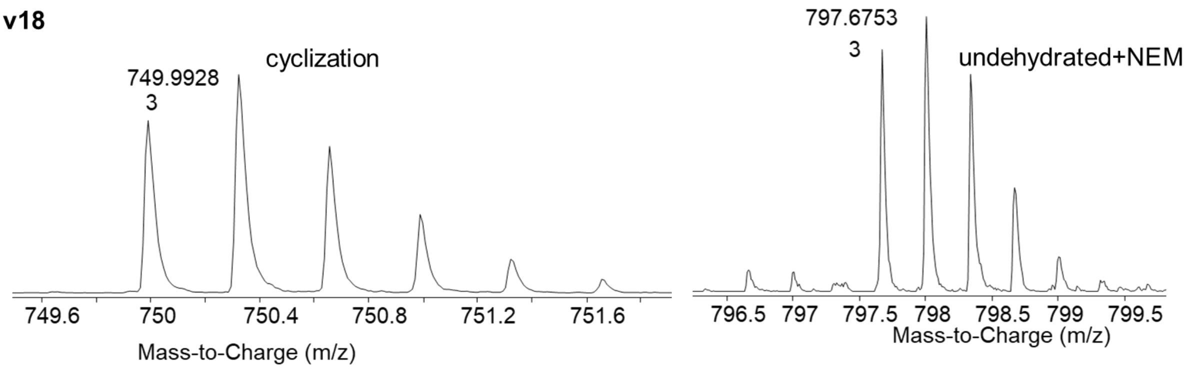


**v19**

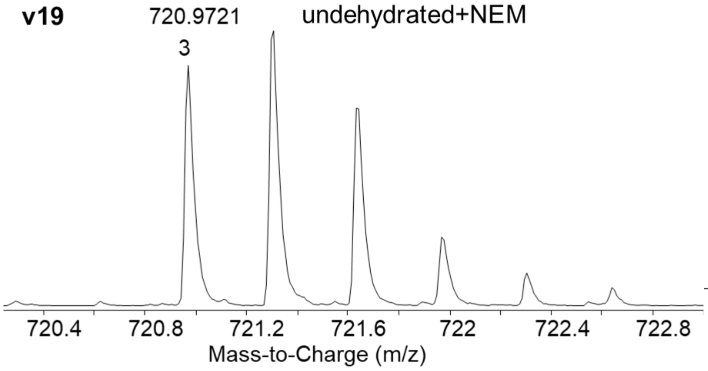


**v20**

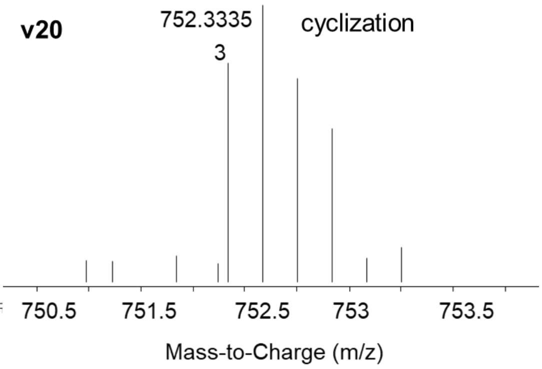


**v21**

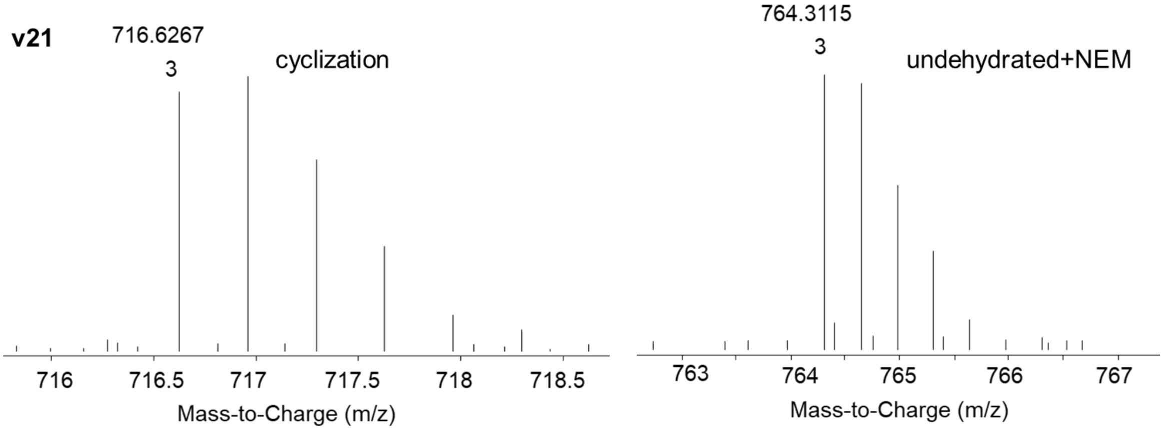


**v22**

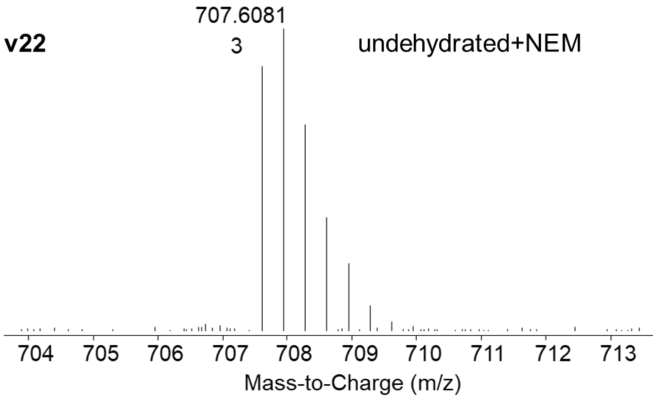


**v23**

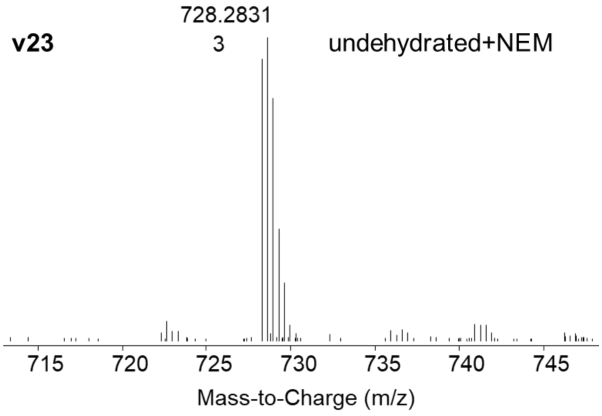

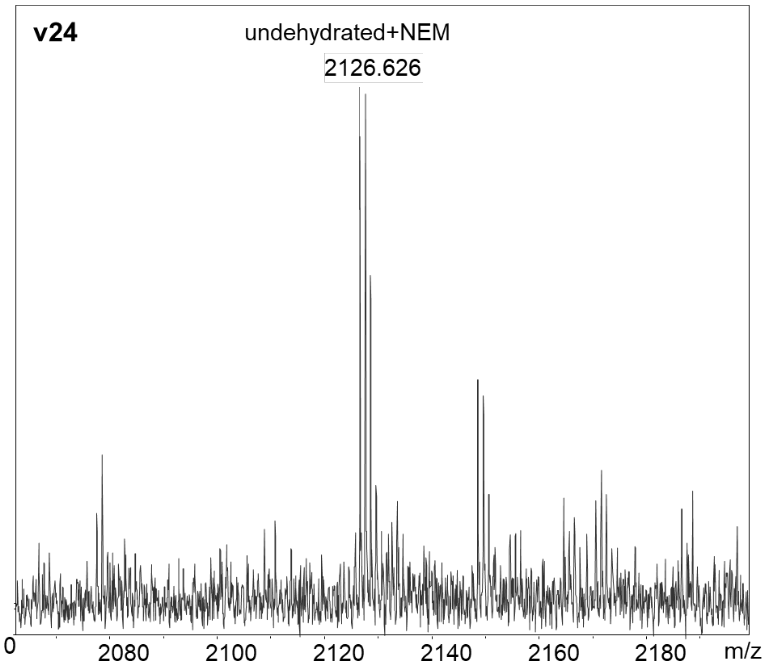
v24
undehydrated+NEM
2126.626
0
2080
2100
2120
2140
2160
2180
m/z

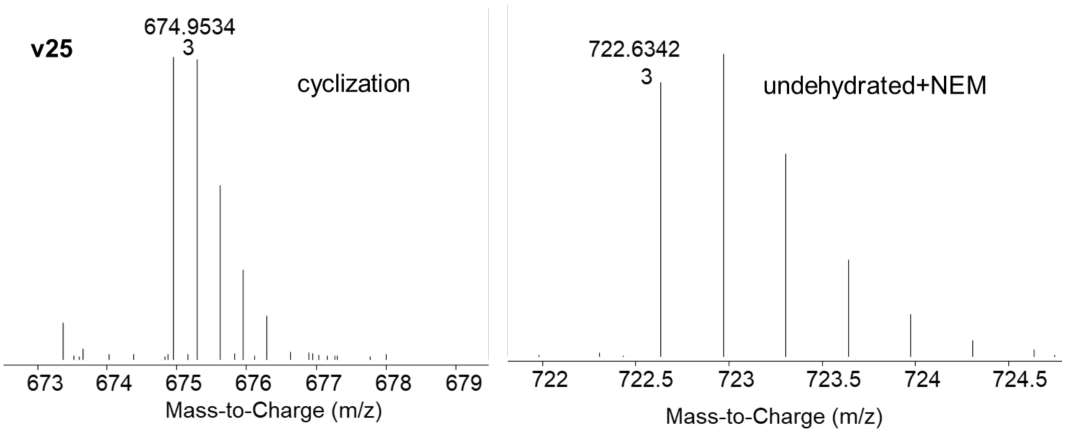
v25
674.9534
3
cyclization
673
674
675
676
677
678
679
Mass-to-Charge (m/z)
722.6342
3
undehydrated+NEM
722
722.5
723
723.5
724
724.5
Mass-to-Charge (m/z)

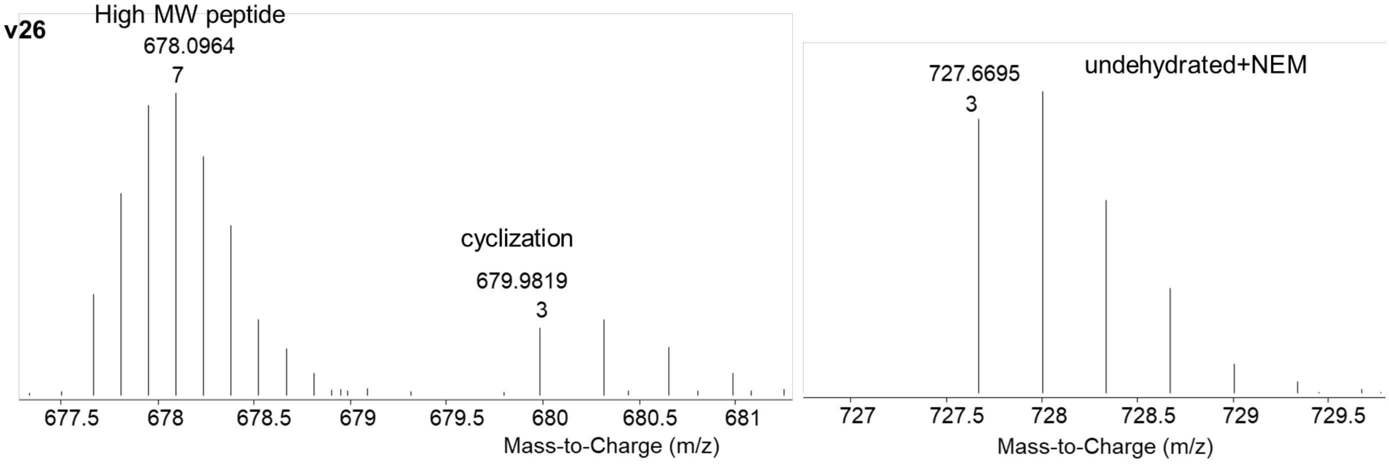
v26
High MW peptide
678.0964
7
cyclization
679.9819
3
677.5
678
678.5
679
679.5
680
680.5
681
Mass-to-Charge (m/z)
727.6695
3
undehydrated+NEM
727
727.5
728
728.5
729
729.5
Mass-to-Charge (m/z)

**v27**

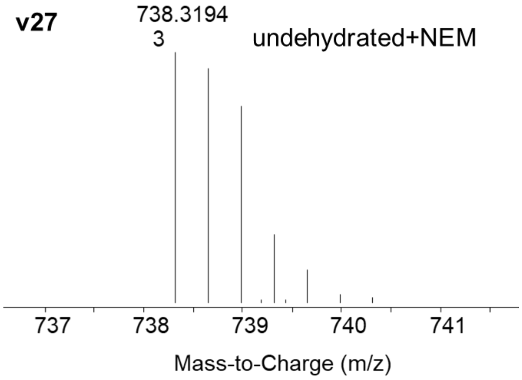


**v28**

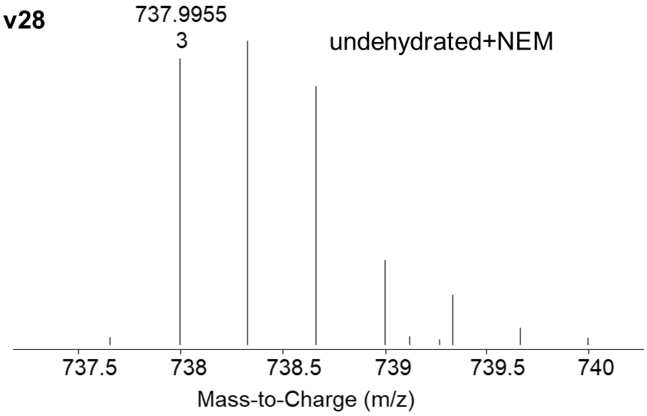


**v29**

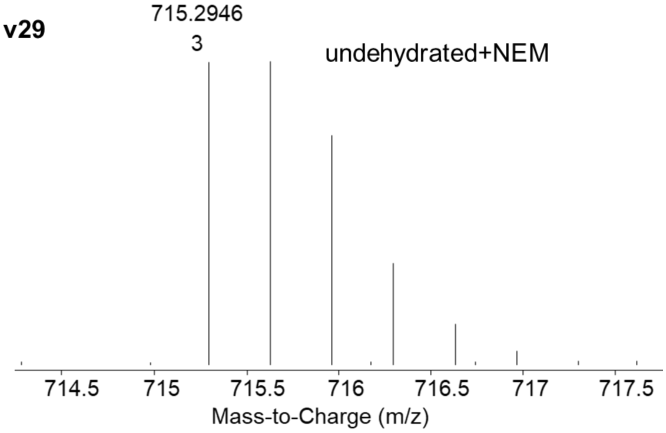


**v30**

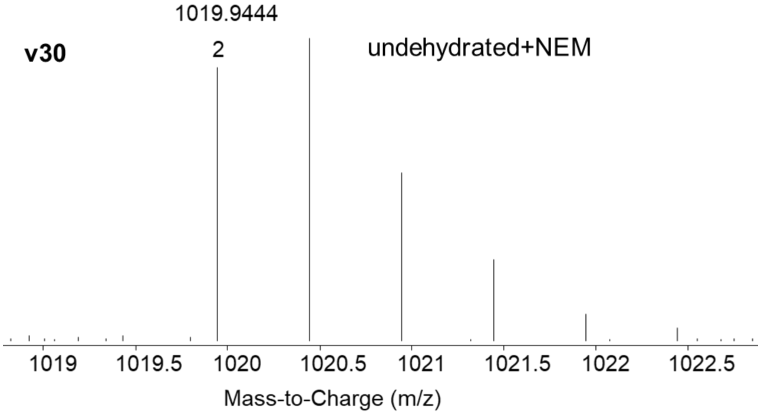


**v31**

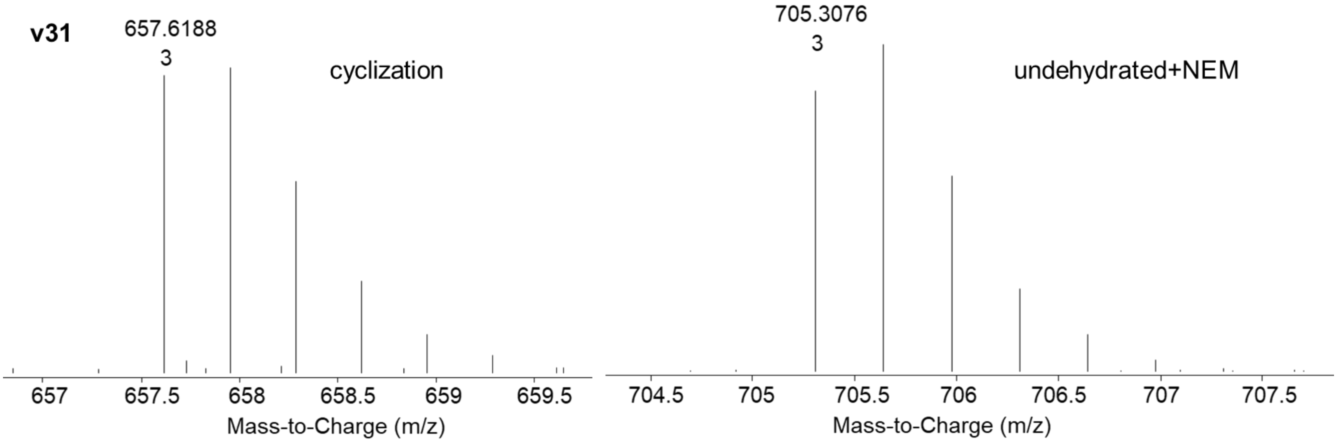

**v32**

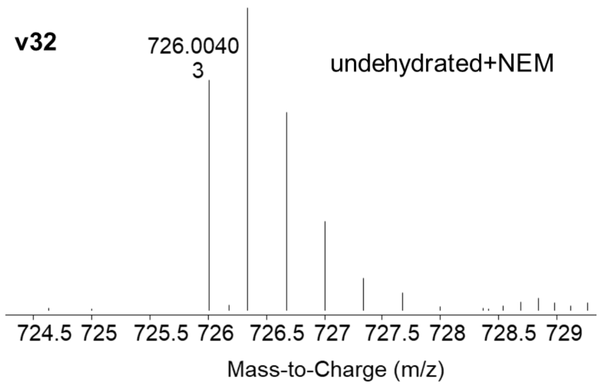
726.0040
3
undehydrated+NEM
724.5 725 725.5 726 726.5 727 727.5 728 728.5 729
Mass-to-Charge (m/z)

**v33**

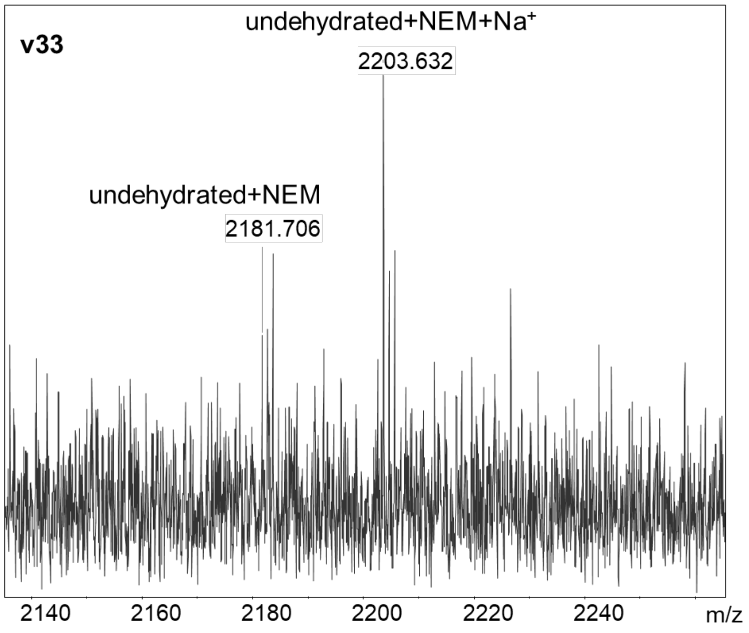
undehydrated+NEM+Na+
2203.632
undehydrated+NEM
2181.706
2140 2160 2180 2200 2220 2240 m/z

**v34**

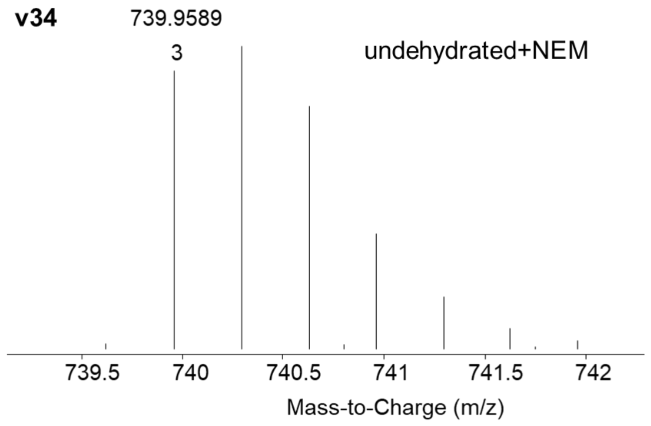
739.9589
3
undehydrated+NEM
739.5 740 740.5 741 741.5 742
Mass-to-Charge (m/z)

**v35**

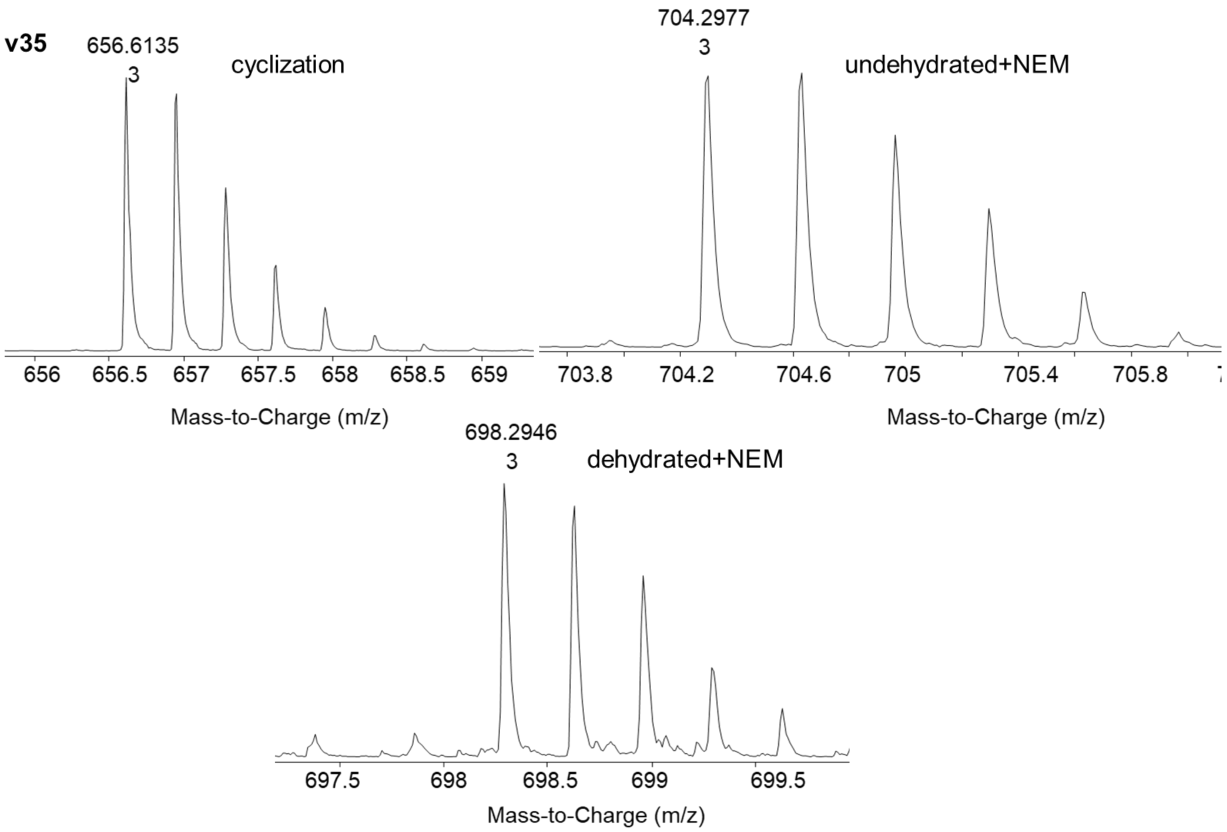
656.6135
3
cyclization
Mass-to-Charge (m/z)
704.2977
3
undehydrated+NEM
Mass-to-Charge (m/z)
698.2946
3
dehydrated+NEM
Mass-to-Charge (m/z)

**v36**

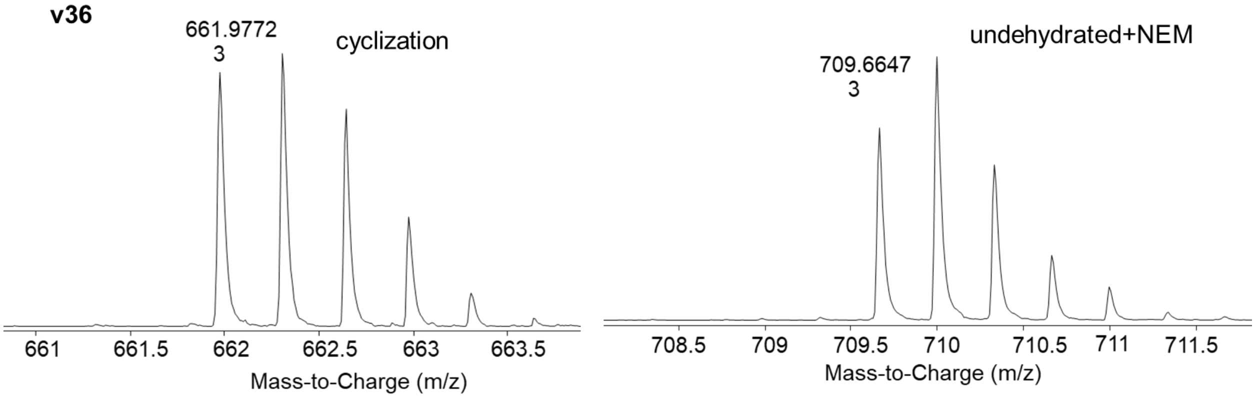
661.9772
3
cyclization
Mass-to-Charge (m/z)
709.6647
3
undehydrated+NEM
Mass-to-Charge (m/z)

**v37**

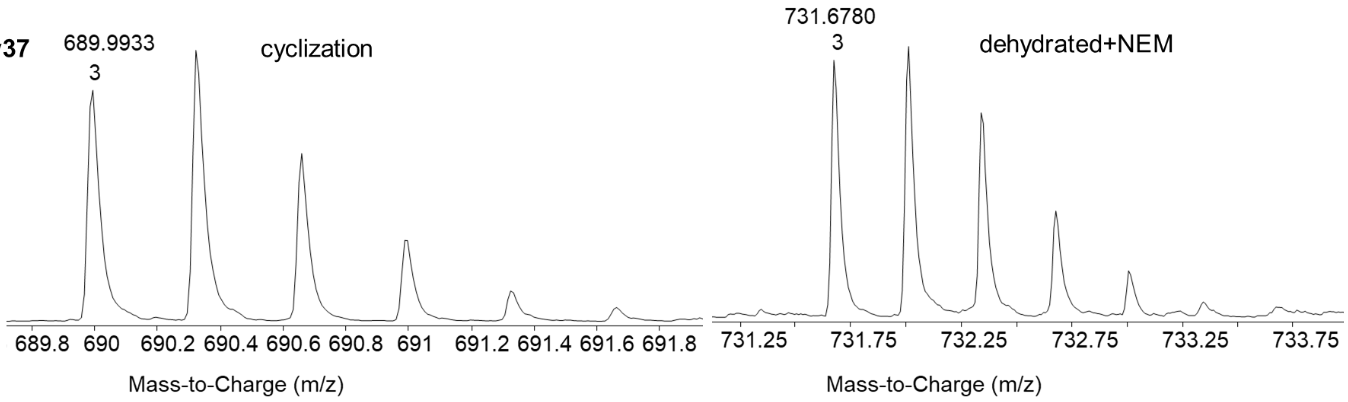
689.9933
3
cyclization
Mass-to-Charge (m/z)
731.6780
3
dehydrated+NEM
Mass-to-Charge (m/z)

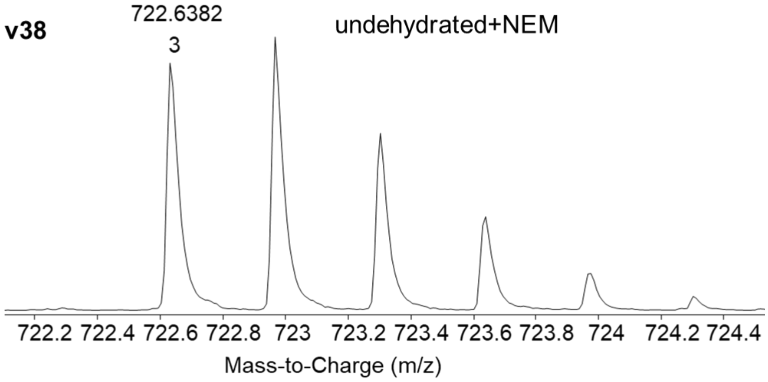
v38
722.6382
3
undehydrated+NEM
722.2 722.4 722.6 722.8 723 723.2 723.4 723.6 723.8 724 724.2 724.4
Mass-to-Charge (m/z)

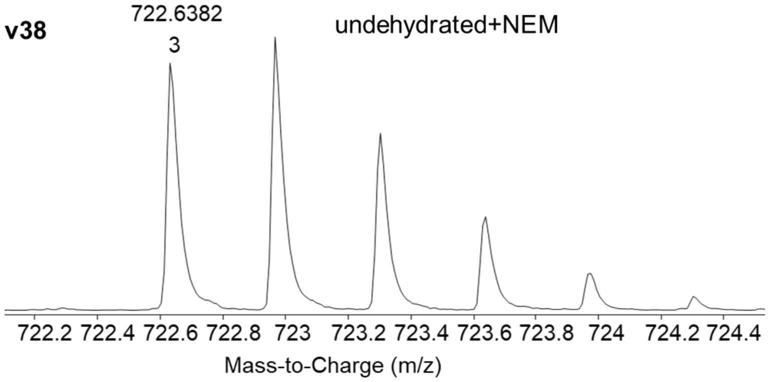
v38
722.6382
3
undehydrated+NEM
722.2 722.4 722.6 722.8 723 723.2 723.4 723.6 723.8 724 724.2 724.4
Mass-to-Charge (m/z)

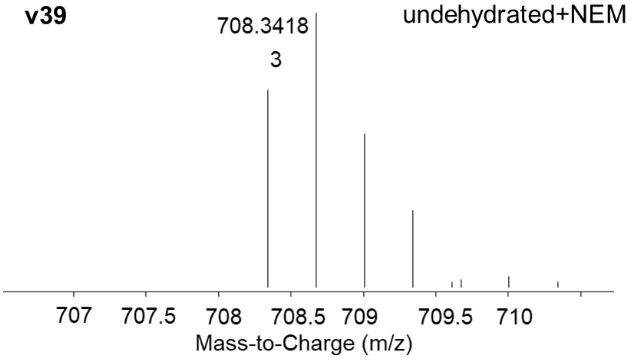
v39
708.3418
3
undehydrated+NEM
707 707.5 708 708.5 709 709.5 710
Mass-to-Charge (m/z)

**v40**

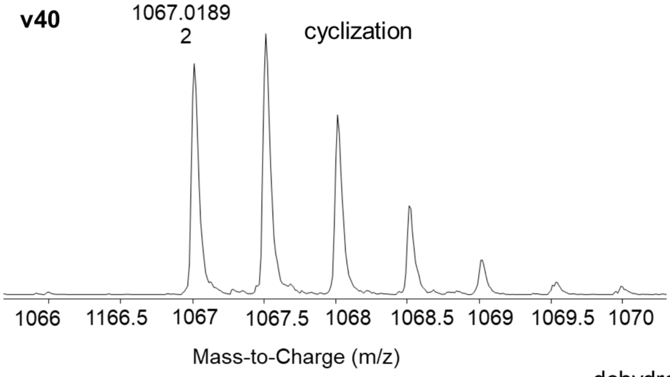
1067.0189
2
cyclization
1066
1166.5
1067
1067.5
1068
1068.5
1069
1069.5
1070
Mass-to-Charge (m/z)

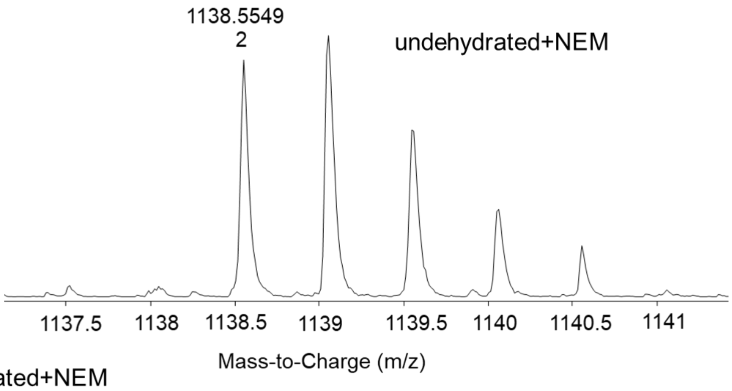
1138.5549
2
undehydrated+NEM
1137.5
1138
1138.5
1139
1139.5
1140
1140.5
1141
Mass-to-Charge (m/z)

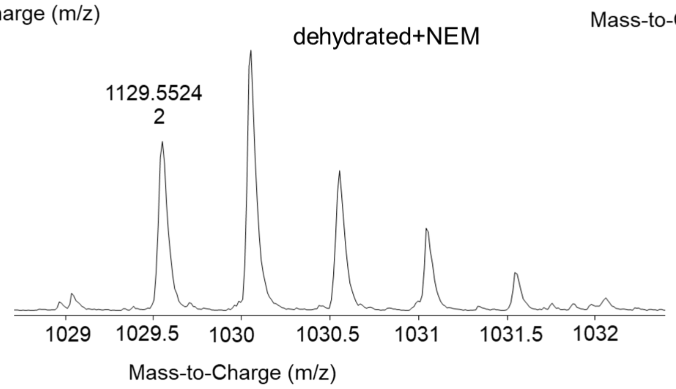
dehydrated+NEM
1129.5524
2
1029
1029.5
1030
1030.5
1031
1031.5
1032
Mass-to-Charge (m/z)

**v41**

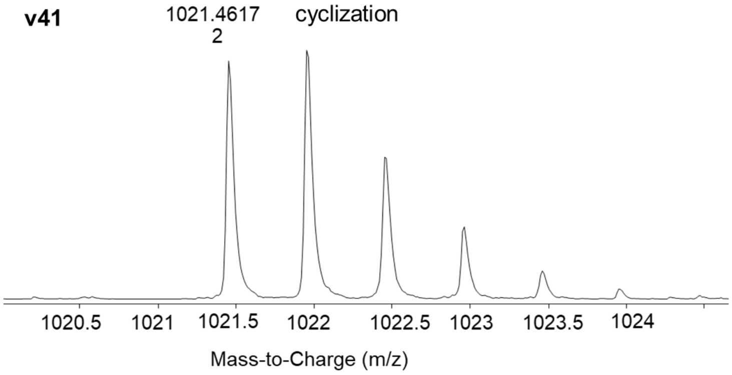
1021.4617
2
cyclization
1020.5
1021
1021.5
1022
1022.5
1023
1023.5
1024
Mass-to-Charge (m/z)

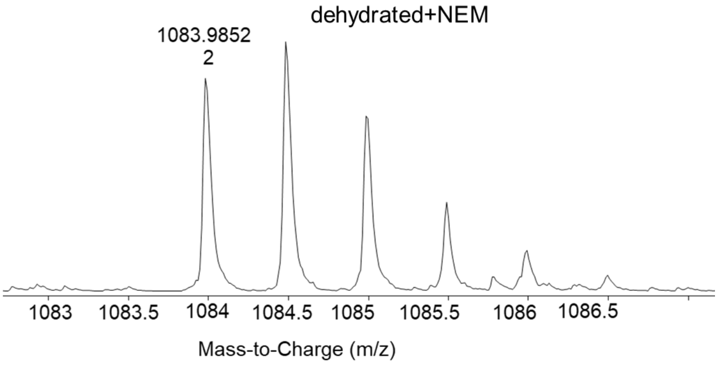
dehydrated+NEM
1083.9852
2
1083
1083.5
1084
1084.5
1085
1085.5
1086
1086.5
Mass-to-Charge (m/z)

**v42**

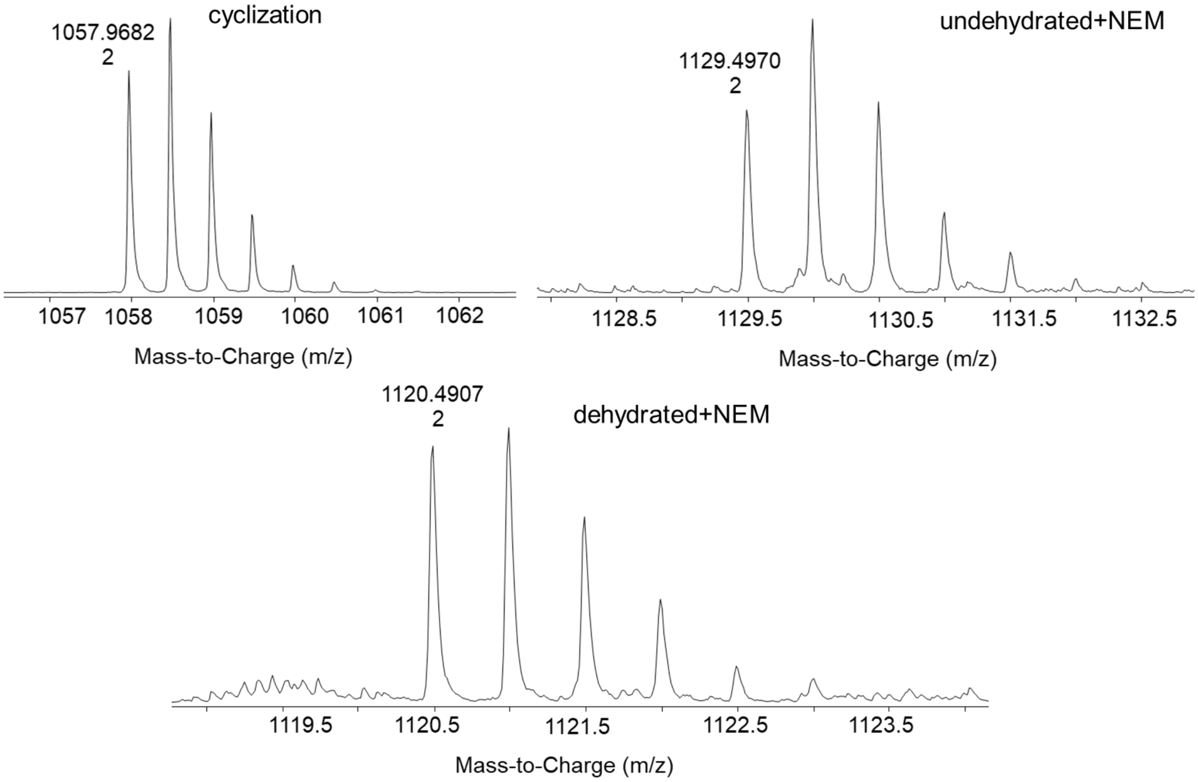
1057.9682
2
cyclization
1057
1058
1059
1060
1061
1062
Mass-to-Charge (m/z)
1129.4970
2
undehydrated+NEM
1128.5
1129.5
1130.5
1131.5
1132.5
Mass-to-Charge (m/z)
1120.4907
2
dehydrated+NEM
1119.5
1120.5
1121.5
1122.5
1123.5
Mass-to-Charge (m/z)

**v43**

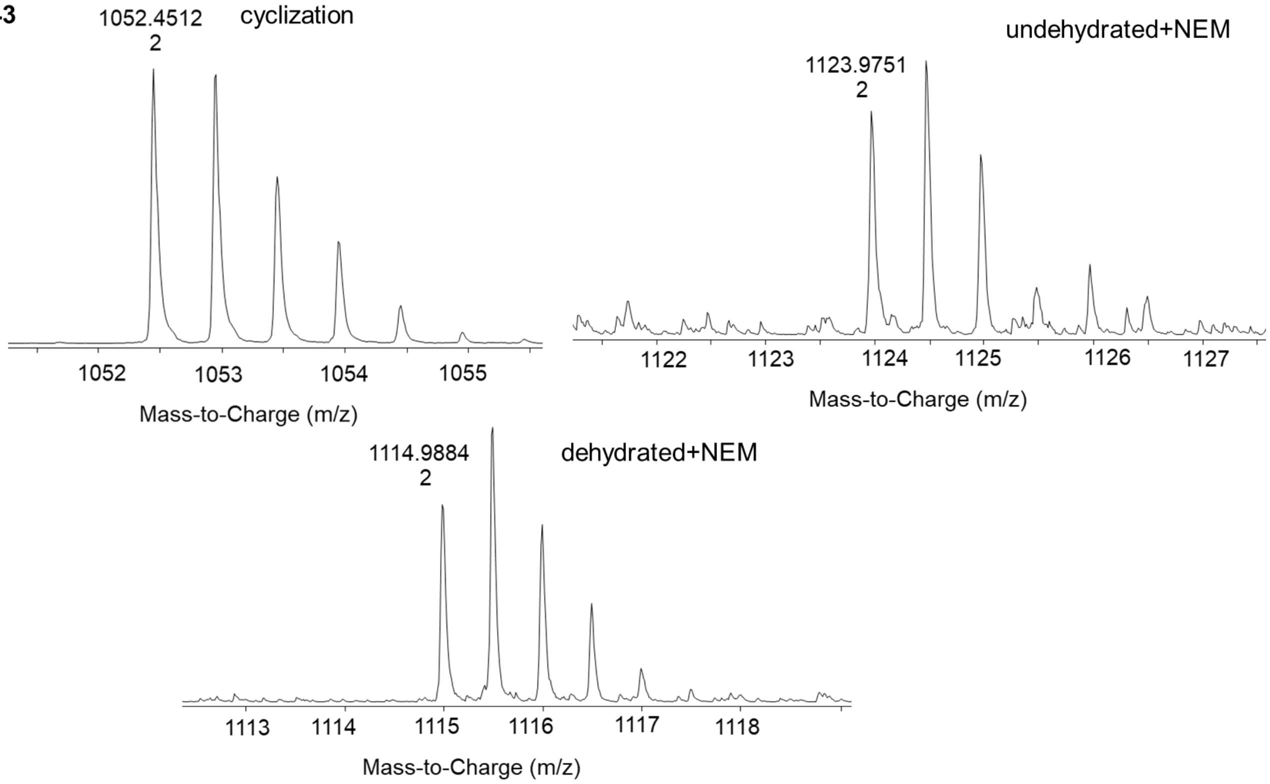
1052.4512
2
cyclization
1052
1053
1054
1055
Mass-to-Charge (m/z)
1123.9751
2
undehydrated+NEM
1122
1123
1124
1125
1126
1127
Mass-to-Charge (m/z)
1114.9884
2
dehydrated+NEM
1113
1114
1115
1116
1117
1118
Mass-to-Charge (m/z)

**v44**

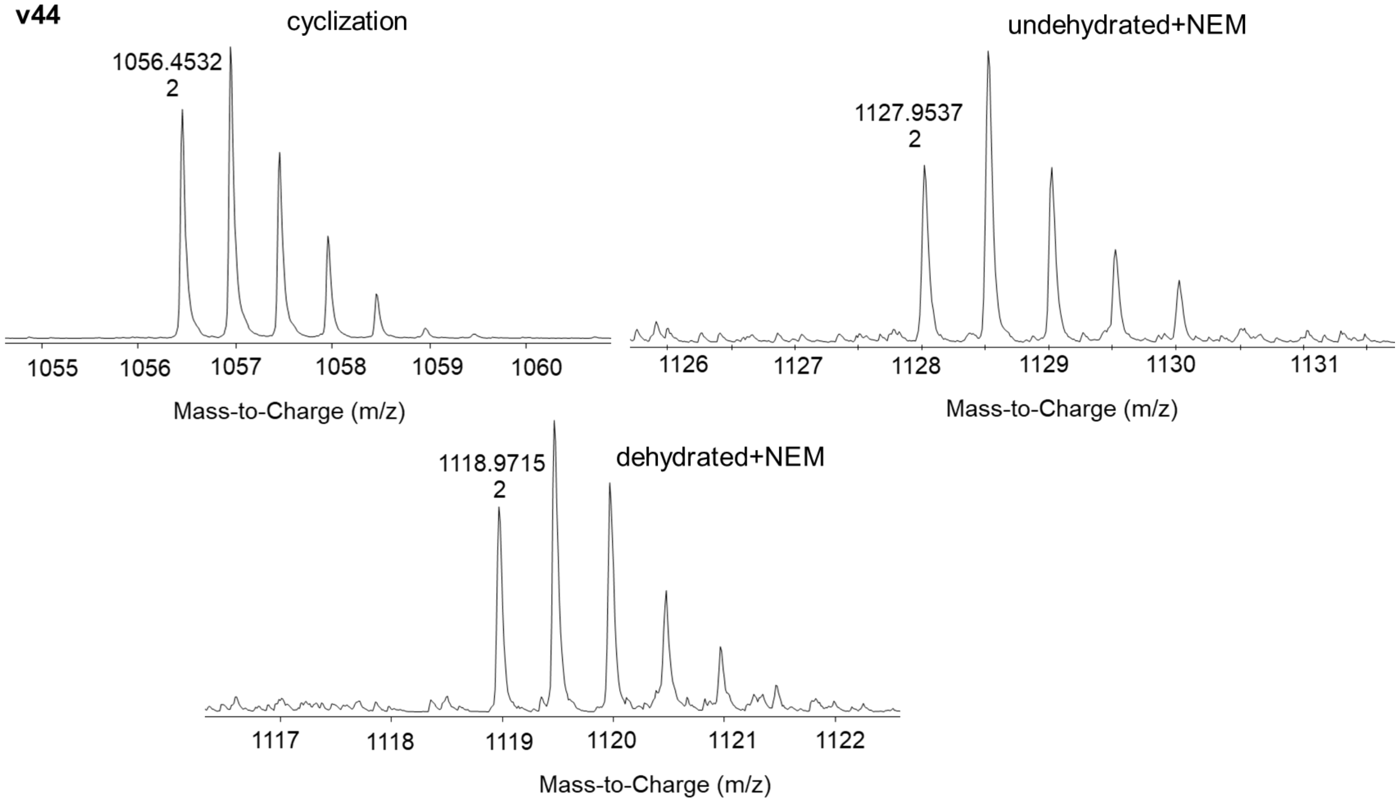
cyclization
1056.4532
2
1055
1056
1057
1058
1059
1060
Mass-to-Charge (m/z)
undehydrated+NEM
1127.9537
2
1126
1127
1128
1129
1130
1131
Mass-to-Charge (m/z)
1118.9715
2
dehydrated+NEM
1117
1118
1119
1120
1121
1122
Mass-to-Charge (m/z)

**v45**

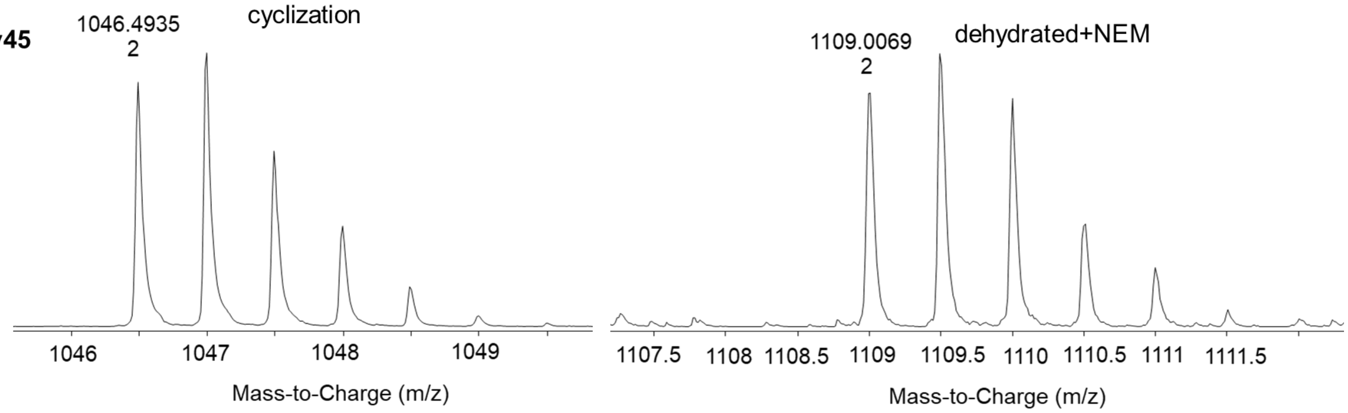
1046.4935
2
cyclization
1046
1047
1048
1049
Mass-to-Charge (m/z)
1109.0069
2
dehydrated+NEM
1107.5
1108
1108.5
1109
1109.5
1110
1110.5
1111
1111.5
Mass-to-Charge (m/z)

**v46**

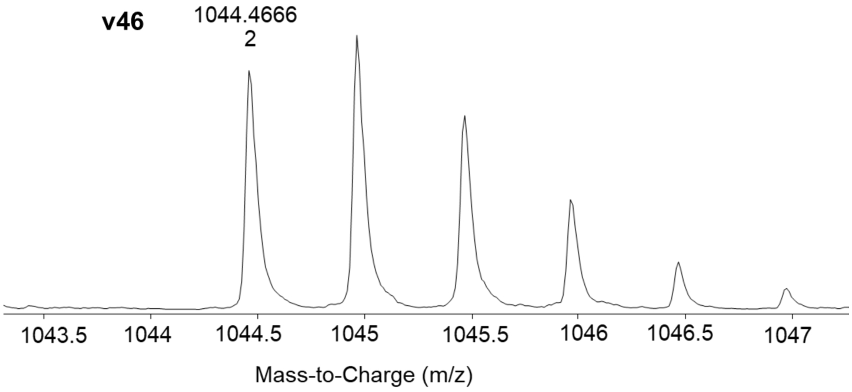
1044.4666
2
1043.5
1044
1044.5
1045
1045.5
1046
1046.5
1047
Mass-to-Charge (m/z)

**v47**

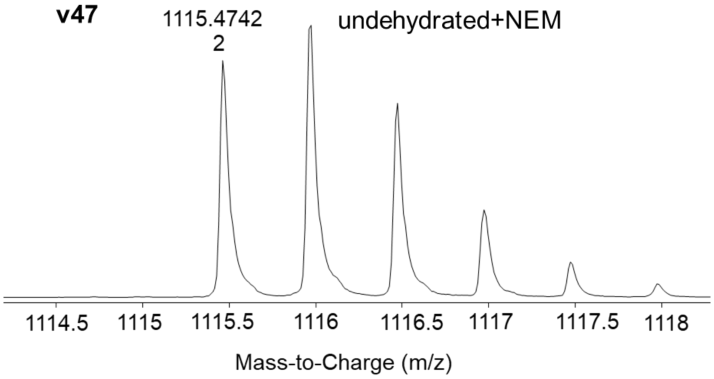


**v48**

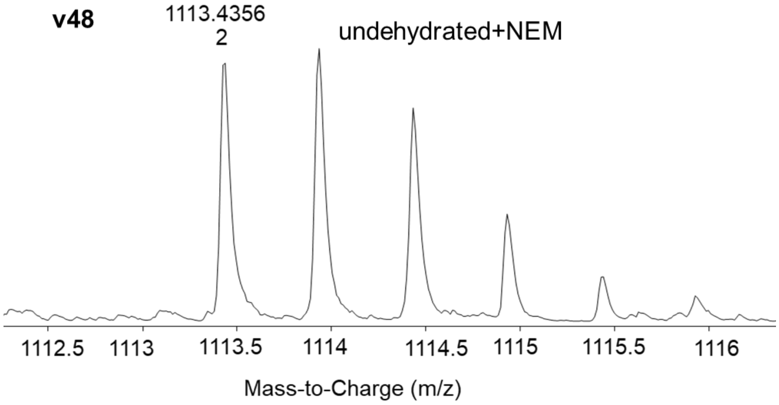


**v49**

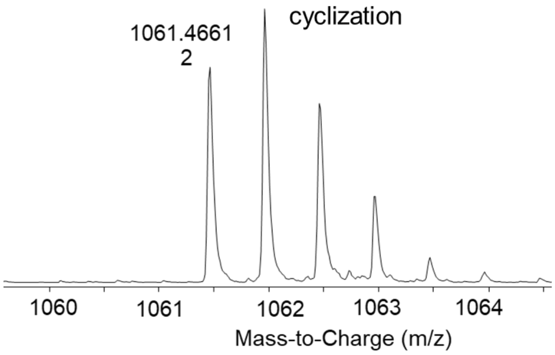


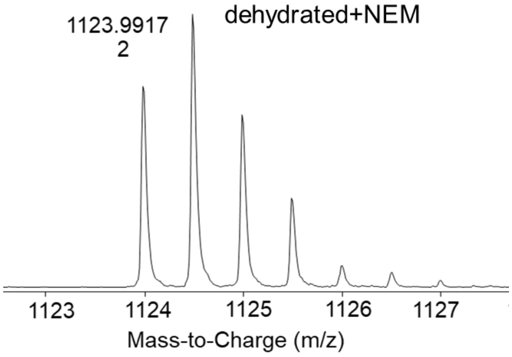


**v50**

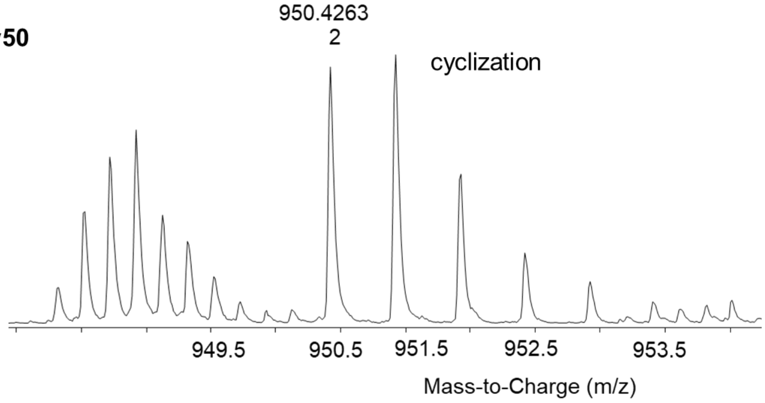


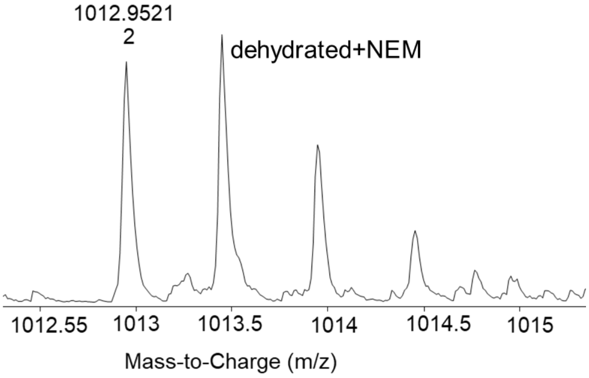

**v51**

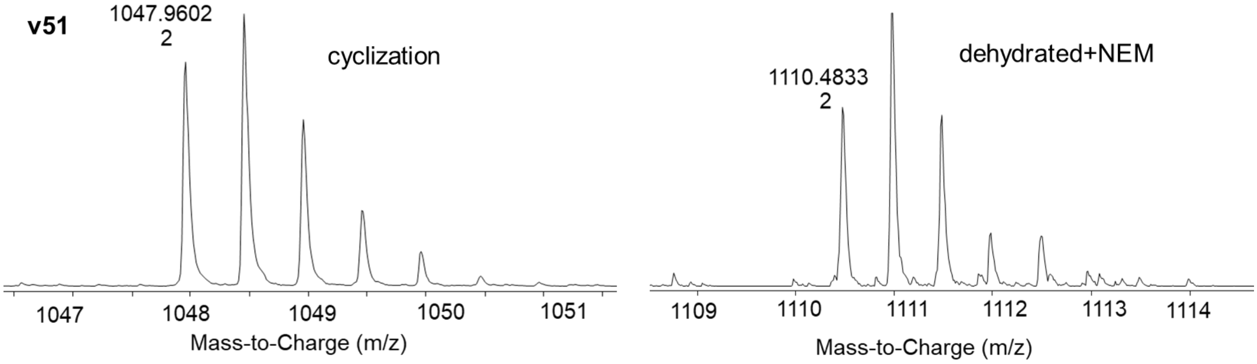


**v52**

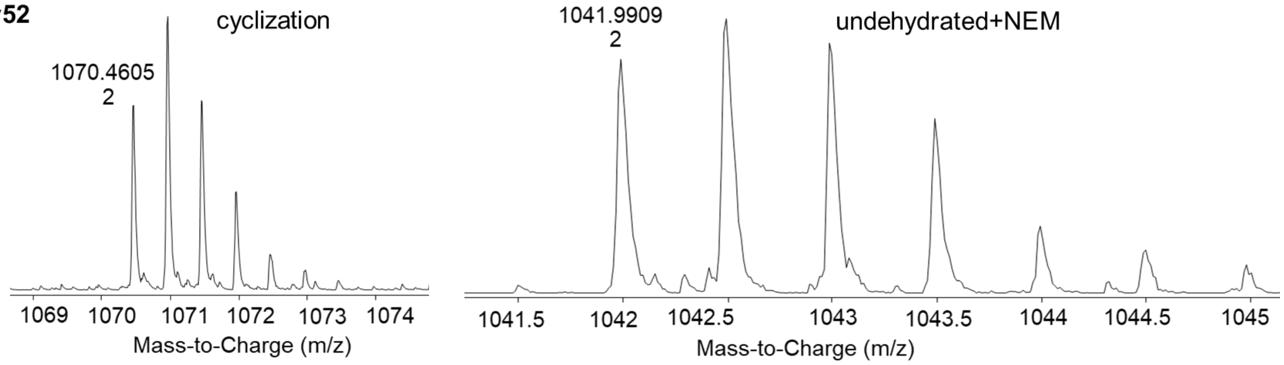


**v53**

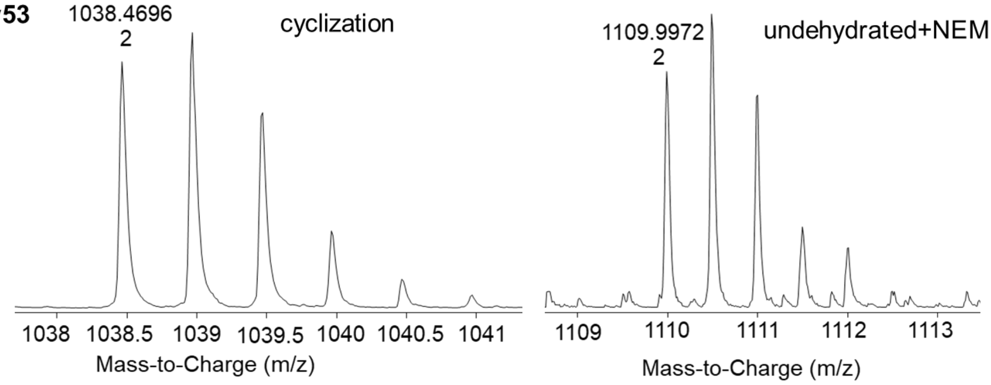


**v54**

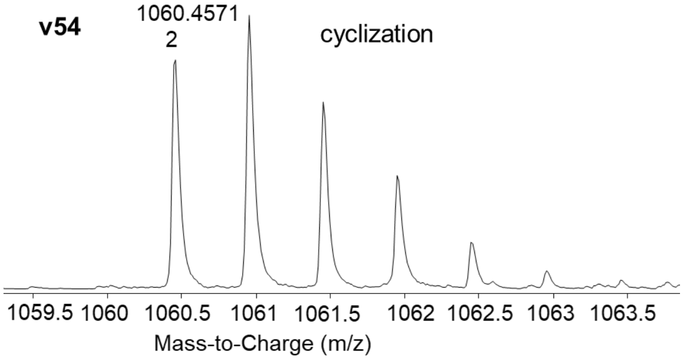

**v55**

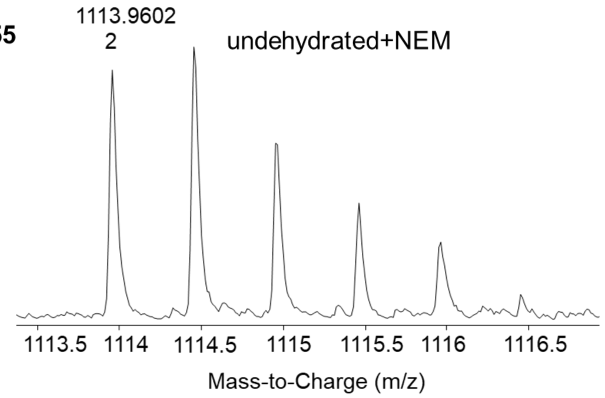


**v56**

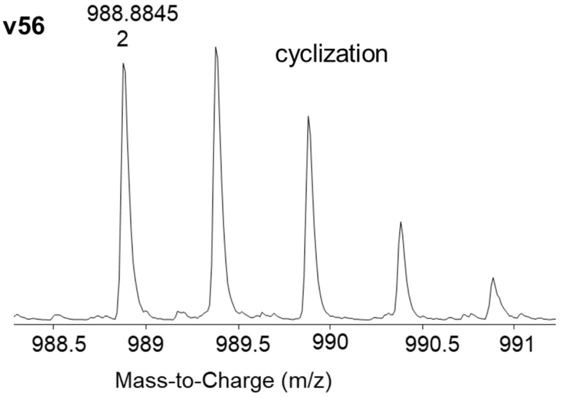


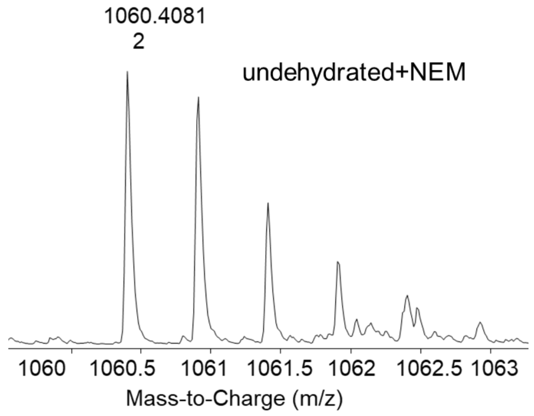


**v57**

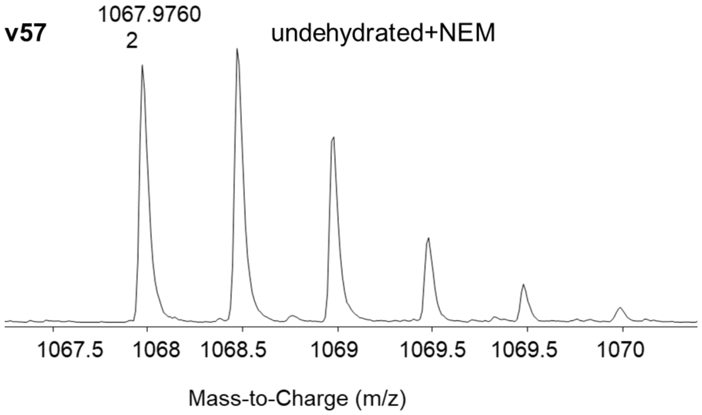


**v58**

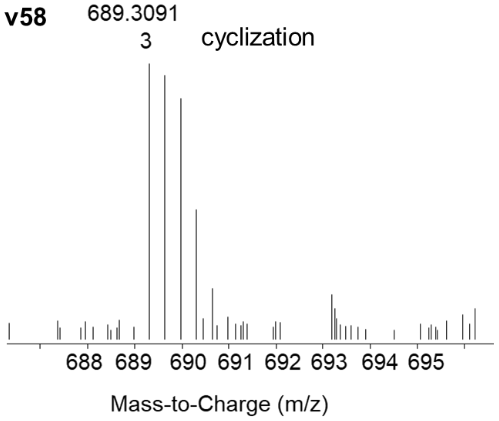


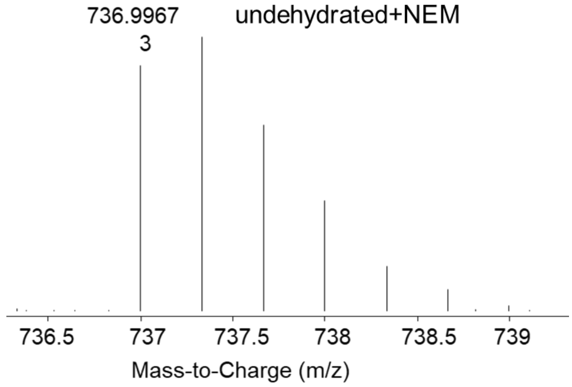

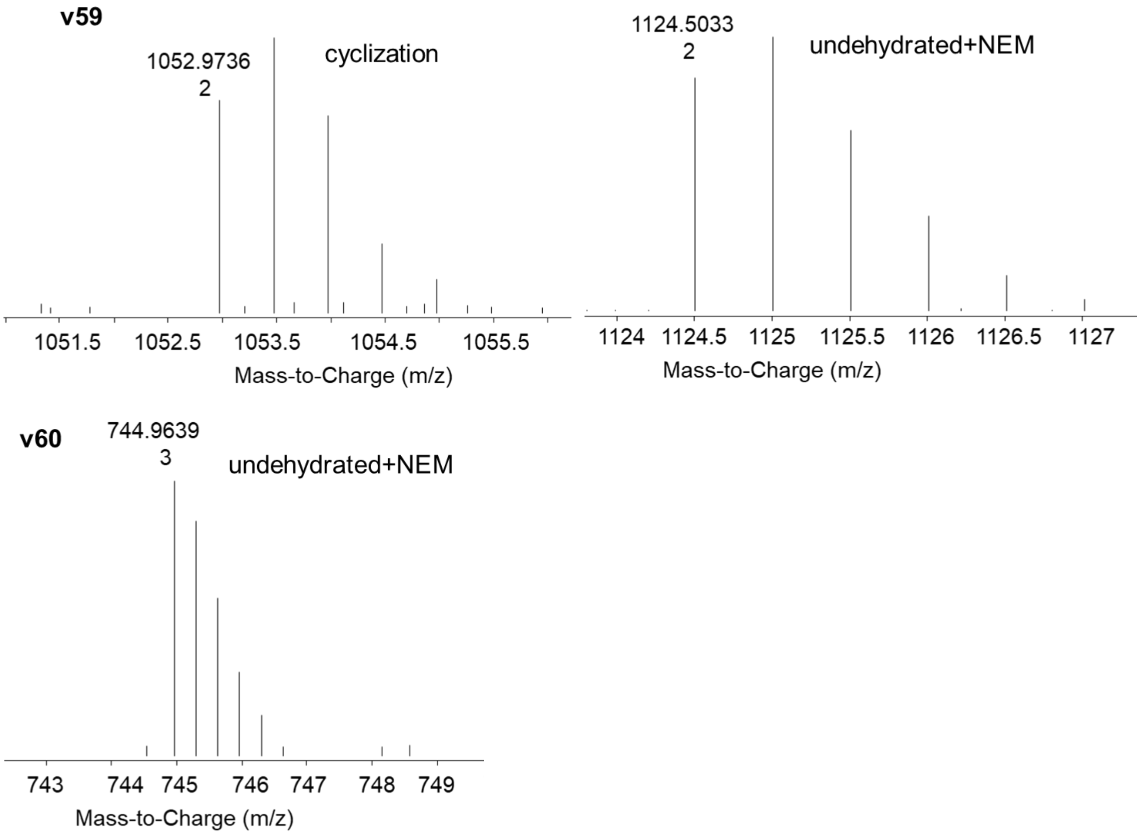


**References**

(1) Mukherjee, S.; van der Donk, W. A. Mechanistic studies on the substrate-tolerant lanthipeptide synthetase ProcM. *J. Am. Chem. Soc*. **2014**, *136*, 10450–10459.

(2) Bobeica, S. C.; Dong, S. H.; Huo, L.; Mazo, N.; McLaughlin, M. I.; Jimenéz-Osés, G.; Nair, S. K.; van der Donk, W. A. Insights into AMS/PCAT transporters from biochemical and structural characterization of a double glycine motif protease. *eLife* **2019**, *8*, e42305.

(3) Vinogradov, A. A.; Chang, J. S.; Onaka, H.; Goto, Y.; Suga, H. Accurate models of substrate preferences of post-translational modification enzymes from a combination of mRNA display and deep learning. *ACS Cent. Sci.* **2022**, *8*, 814-24.

(4) Lin, T.-Y.; Goyal, P.; Girshick, R.; He, K.; Dollár, P. Focal Loss for Dense Object Detection. *arXiv* **2017**, arXiv:1708.02002

(5) Akiba, T. S., S.; Yanase, T.; Ohta, T.; Koyama, M. 2019. In *Proceedings of the 25th ACM SIGKDD International Conference on Knowledge Discovery and Data Mining* 2019.

(6) Li, B.; Sher, D.; Kelly, L.; Shi, Y.; Huang, K.; Knerr, P. J.; Joewono, I.; Rusch, D.; Chisholm, S. W.; van der Donk, W. A. Catalytic promiscuity in the biosynthesis of cyclic peptide secondary metabolites in planktonic marine cyanobacteria. *Proc. Natl. Acad. Sci. U.S.A.* **2010**, *107*, 10430-10435.
